# Wild great tits show social resilience in response to partner loss

**DOI:** 10.64898/2026.09.15.751691

**Authors:** Adelaide Daisy Abraham, Josh A. Firth, Ben C. Sheldon

## Abstract

Social relationships are foundational to animal societies, but individual social connections differ hugely in importance and strength. Key relationships, such as those between breeding partners, may be prioritised and more highly invested in by individuals, but a key question is the extent to which social flexibility allows individuals to respond to the loss of these key social relationships. Here, we use a detailed longitudinal dataset of individual breeding and social behaviour in wild great tits (*Parus major*) to characterise flexibility in social behaviour after the loss of their breeding partner (‘widowing’). We show transient increases in two measures of sociality following the widowing event. Individuals’ relationships with their replacement partner were initially weaker than those between non-widowed birds, but converged by the breeding season. Hence, we show that birds are robust to the loss of key social relationships and that great tits show a flexible ability to return to social equilibrium. Strong social resilience has implications for the response of populations to change and loss.

## Introduction

Animals show a wide diversity of social behaviours, with long-term social relationships being important for many species (Taborsky, Cant, and Komdeur 2021). An important social relationship in sexually reproducing animals is that between breeding partners, particularly in monogamous mating systems (Kvarnemo 2018). Socially monogamous pair bonding is particularly frequent in birds (∼90% of species (Choudhury 1995)) but also found in fish (18+ families (Whiteman and Côté 2004)), mammals (∼5% (Kleiman 1977), and some lizards (Bull 2000)). In such systems, the choice of a mate has implications for reproductive fitness (Jennions and Petrie 1997; Ihle, Kempenaers, and Forstmeier 2015) but also often for survival (Searcy 1982; Culina, Lachish, and Sheldon 2015). A closer or more long-term relationship with a mate has also been shown to increase reproductive success (Yokoi et al. 2016; Culina, Firth, and Hinde 2020). For these reasons, individuals may be expected to invest both time and energy into bonding with an appropriate mate. This bond-building can take place or be influenced by pre-existing social relationships formed in the non-breeding season, showing the dynamic and temporal nature of animal social bonds (Cheetham et al., 2008; Senar et al., 2013; Yokoi et al., 2016).

There is an inherent temporal nature to social relationships in almost all animal societies due to the gain and loss of individuals. While there is a large body of research into the numbers and predictors of individuals lost from a population (Bowler and Benton 2005; Reinke, Miller, and Janzen 2019; Snyder-Mackler et al. 2020), much less is known about how the loss of these individuals impacts those that remain. Because of the investment of time and energy into a potential mate, the disappearance of a partner may represent a particularly important disturbance, requiring the widowed individual to either forgo breeding or search for a new mate. While there is evidence for the effects of widowing during the breeding season on reproductive output (Duckworth 1992; Thomas and Wolff 2004), there is very little research into how the loss of a partner during the non-breeding season may impact behaviour. This lack of research is partially due to the difficulty of obtaining detailed social interaction data across time for many individuals, with associated life history and breeding data.

Great tits (*Parus major*) provide an opportunity to obtain long-term longitudinal social data across a number of social contexts. As a socially monogamous bird species with seasonal sociality they forage and raise a brood with a single partner during the breeding season. In contrast, during the winter non-breeding season, they feed in fission-fusion flocks, with a dynamic social system (Hinde 1952). The bonds great tits form in the winter can influence their pair bonding (Firth et al. 2018) and their spring pair bonds can also shape their winter foraging behaviour (Firth et al 2015; Abraham et al 2025). The earlier in the winter a pair bond forms between two great tits, the earlier in the spring they will breed on average, which is associated with higher fitness outcomes (Culina, Firth, and Hinde 2020). However, great tits also have high annual mortality (40-60% (Culina, Lachish, and Sheldon 2015)), with interspecific competition, predation, and low food availability all decreasing survival (Clobert et al. 1988). Therefore, it is likely that there will be developing pair bonds that are interrupted by the death of one of the individuals involved. Previous research has shown that great tits adjust their social associations in response to the loss of a close associate (Firth et al 2017), and that there are distinctive patterns of social behaviour in birds which are ‘divorcing’ their mate, as opposed to those maintaining a previous pair bond (Abraham, Sheldon, and Firth 2025). This makes them an appropriate species for investigating how individuals may respond to the loss of a breeding partner.

We hypothesised that, due to behavioural investment in pre-existing bonds and the importance of winter sociality for breeding success, great tits that lost their previous year’s mate would show changes in overall social behaviour. We theorised that this may occur due to the birds attempting to find a replacement for their lost mate. Previous work has suggested that great tits are resilient to population turnover (Aplin et al. 2015) and the loss of social associates (Firth et al. 2017), but that they prioritise social relationships with their breeding partner (Firth et al. 2015). We hypothesised that this prioritisation of mate relationships would lead to a social response when the partner vanished from the population.

Testing this hypothesis requires a multiple-year dataset which includes breeding pair identities and social behaviour in the non-breeding season, in order to draw connections between social processes in the two periods. The Wytham Woods long-term tit study in Oxford, UK, meets these requirements, providing a detailed dataset with which to answer questions such as these. We used three years of social network data gathered from winter feeder movements, combined with five years of breeding pair records, to identify widowed birds and analyse their social behaviour after the widowing event, to examine how individuals respond socially to the loss of a mate during the crucial pair formation phase.

## Methods

### Data preparation

The great tits in the Wytham Woods population used in this study breed almost exclusively in artificial nestboxes. These nestboxes are checked regularly during the spring (see Perrins, 1965) and British Trust for Ornithology rings and PIT (passive integrated transponder) tags are used to identify breeding individuals.

During the non-breeding season, great tits no longer engage in territorial behaviours associated with the spring and feed in foraging flocks which move around the woodland (Aplin et al. 2012). For 13 weekends (Saturday and Sunday only) across three winter seasons (3 December 2011 to 26 February 2012; 1 December 2012 to 24 February 2013; 30 November 2013 to 23 February 2014), supplementary feeders were placed throughout the woods at 250m even grid spacing. Visits of birds to the feeder were recorded with RFID (radio frequency identification) antennae that identified bird PIT tags, placed on individuals during the breeding season and through winter ringing. Feeder data was recorded every one-third of a second. Using this visit data, flocking events were identified using a Gaussian mixed model through the asnipe package (Farine 2013), as described in Farine et al. 2015. Networks were then calculated using these flocking events, with an association between individuals referring to them being observed in the same flocking event. Where used, association strength was calculated as winter association score:

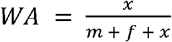

where *x* is the number of flocking events both birds are observed in, *m* is the flocking events only the male was observed in, and *f* is the number of flocking events only the female was observed in.

When identifying widowed birds, we classified a bird as ‘vanished’ if it was observed at the feeders during the winter at least once, then no longer observed visiting feeders, as well as never observed at feeders or breeding in any subsequent years (see Supplementary Information for a discussion of the accuracy of this designation). The bird’s previous mate was then classified as a ‘widow’. There may be multiple mechanisms by which a bird vanishes from our records (mortality, emigration, behavioural change) but — as we are interested in their persistence in their population — for this analysis we have classified all of them similarly as vanishing. ‘Widowing date’ refers to the last day a bird’s previous mate was observed at the feeders. As pairs were only included when one individual died during the winter experimental period, and the other was observed for the entire experimental period, data requirements were restrictive, and the number of pairs suitable for analysis was relatively small.

**Fig 1:**
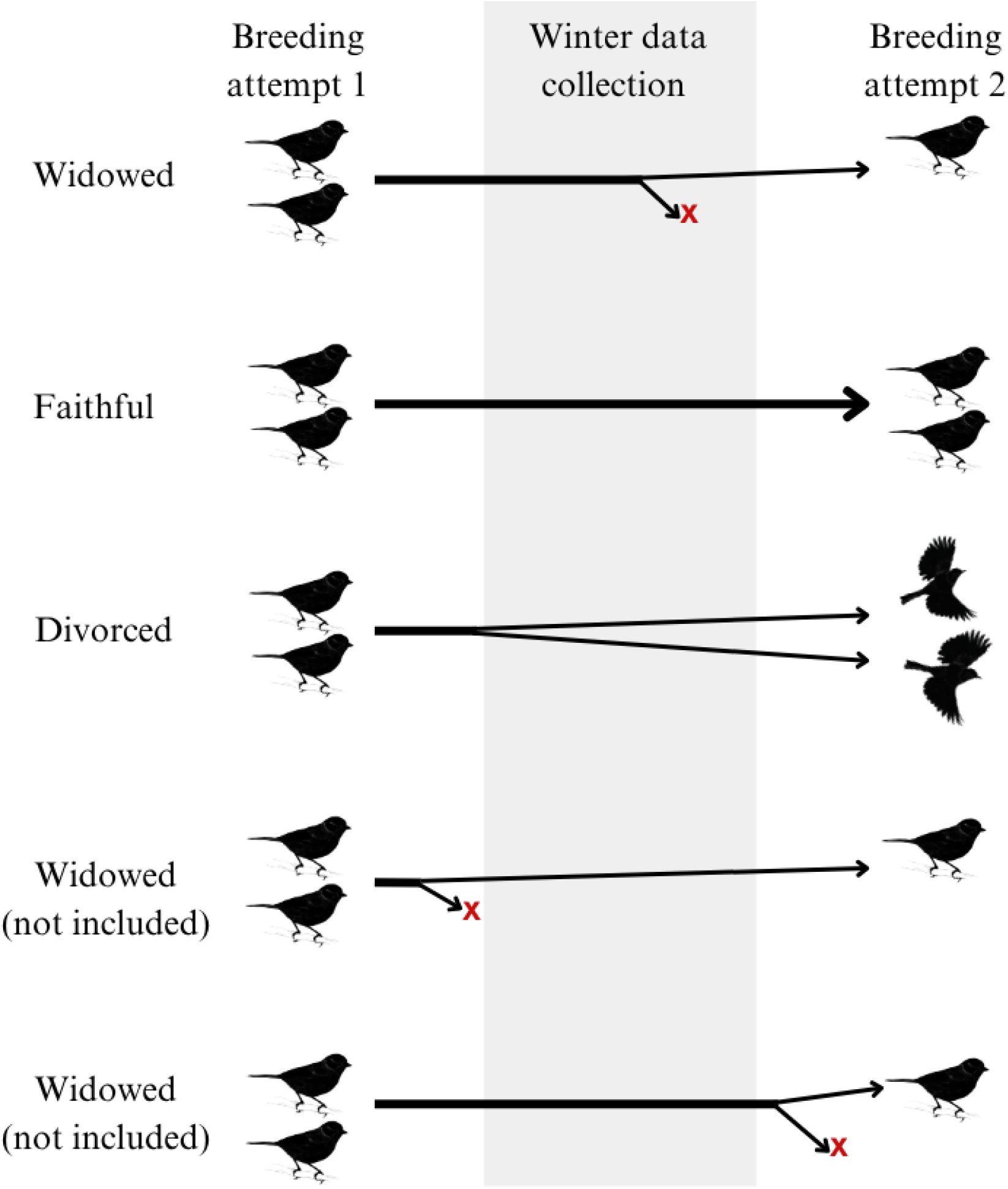
Pair classification of birds included in this paper. A red ‘X’ depicts when a bird leaves the study population through death, behavioural change, or emigration. The dates included in ‘winter data collection’ varied between the three years included in the study, but were between late November/early December to late February.

We identified bird pairs using breeding records from nestboxes, where ‘pair’ (or ‘mate’) refers to birds which were seen sharing a nestbox during breeding season. This is opposed to ‘partner’, which refers to an individual’s closest opposite-sex associate during the non-breeding season, which may or may not go on to be their mate. As found in previous research, a strong ongoing partner association between a pair is observed when they go on to breed together again, but not when they are divorced (Abraham, Sheldon, and Firth 2025). This study investigated the impact of the loss of a partner as a proxy for the loss of a faithful mate, and as such was focused on widowed pairs which had bred together previously and which we predicted would have gone on to mate together again.

In order to predict which widowed pairs would have been faithful, we built a Bayesian model to predict which pairs would have stayed together. This was a Bernoulli model trained on behavioural data from non-widowed pairs with known faithful/divorcing status, using variables describing pair and individual social behaviour and breeding success. As widowed pairs had variable amounts of winter social data available before the widowing, we assessed model accuracy using a leave-one-out test and train procedure, successively using data up to and including each experimental day. Our final model had approximately 80% prediction accuracy, increasing to 85% accuracy when trained on more than 30 days of data. When trained on more than 15 days of data, our model performed better than simply assigning all pairs faithful status. The model was then applied to the widowed pairs to predict their status. The details of this model and a sensitivity analysis of the effect of pair classification are provided in the Supplementary Information.

Analyses of behavioural changes included non-widowed divorcing and faithful pairs. As our interest was in predicted faithful widowed pairs, comparison allowed other pair types to function as controls. Predicted divorcing widowed pairs were a control where widowing occurred, but we did not predict a behavioural response as we assumed there was limited social investment between the pair. Faithful non-widowed pairs allowed us to account for pairs which were socially invested, but without widowing, and divorcing non-widowed pairs provided a further control of non-associated non-widowed pairs. For these comparative analyses, if a pre/post widowing value was required for non-widowed pairs, the ‘false’ widowing date was drawn randomly from the observed widowing dates of widowed pairs. In case our observed results were due only to the particular random distribution we used, a sensitivity analysis using multiple permutations of this random widowing date generation is provided in the Supplementary Information.

### Behavioural change metrics

We expected that losing a pair bonded partner would result in a change in the behaviour of the focal widowed individual. We analysed whether there was a significant difference in nine behavioural measures (including 7 social metrics) before and after widowing. For each model, we analysed whether there was a difference in behaviour between the entire period before widowing when compared to the entire period after widowing. In order to account for the possibility of only short-term change in behaviour, we also modelled behavioural change in the experimental weekend immediately before and immediately after the widowing date on a week-by-week basis. A short summary of each metric is provided here (and in Figure 2), with full details in the Supplementary Information.

**Fig 2:**
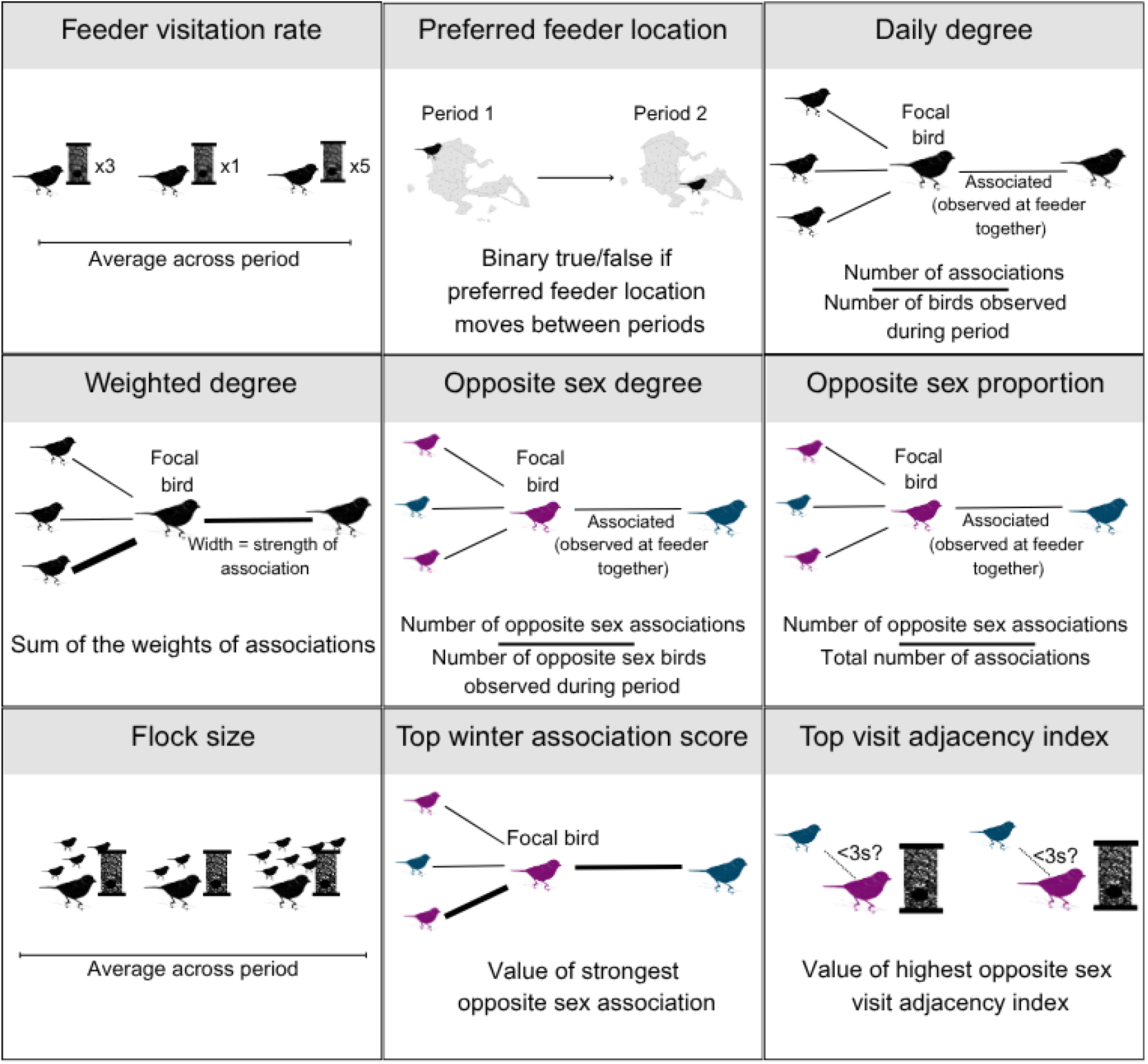
Schematic diagram of each of the behavioural measures used in this analysis.

*Feeder visitation rate*: Average feeder visits per experimental day

*Preferred feeder location*: TRUE/FALSE whether the individual changed preferred feeders after the widowing date

*Daily degree*: The number of associates the individual was observed with

*Weighted degree*: Number of associates weighted by association strength

*Opposite sex degree*: Number of associates of the opposite sex

*Opposite sex proportion*: Proportion of associates of the opposite sex (of known sex associates)

*Flock size*: Size of flocking events the individual was observed in

*Top winter association score*: Top strength of association (simple ratio index) with a bird of the opposite sex

*Top visit adjacency index*: Highest rate of visiting the feeder within 3 seconds of an opposite sex bird

### Model estimation

For models comparing behaviour before and after the widowing date, we set the average of the social behavioural measure in each period for each individual as the response variable, and analysed whether there was a difference in these averages. For the models estimating short-term responses only, we once again averaged individuals’ behavioural measures in the relevant weekends and used this as the response variable.

In all models, in order to maximise power, all four pair statuses were included (widowed faithful, widowed divorcing, divorcing, and faithful), but in the main body of this paper, only the results for widowed faithful birds are presented, as this is the relevant focal class of the study. Faithful non-widowed birds were used as a reference level in all models, so that a significant effect for faithful widowed birds represents a significant difference from the control faithful non-widowed pairs. The divorcing classes provide a useful additional comparison and control. Additional results, and models split by sex, are included in the Supplementary Information.

All models were run using the glmmTMB (Brooks et al. 2017) package in R (R Core Team 2021). In all cases except where specified in the Supplementary Information, models were run as lognormal, including a baseline zero inflation term to account for the presence of zeroes in the data. The basic model structure was pair status (faithful, divorcing, faithful widowed, divorcing widowed) in interaction with widowing stage (pre-widowing, post-widowing), and year and ID as random effects. Models were checked using residuals generated with the DHARMa package (Hartig 2018) as well as visual checks of predictions.

## Results

Of the 627 breeding pairs recorded during the period of this analysis, 228 (36%) had some winter feeder data for both individuals, allowing a joint analysis of breeding and non-breeding behaviour (this proportion is broadly as expected given individual annual survival of ∼50%). 98 of these pairs were unable to be used as they were newly formed and therefore didn’t have a previous breeding season. Of the remaining pairs, 27 were suitable for analysis as widowed pairs, as they had one individual survive to be observed again and one individual vanish during the winter non-breeding season. The Bayesian faithfulness model predicted that 18 (66.67%) would have been faithful, and that 9 (33.33%) would have been divorced (detailed model results in Supplementary Information). This is compared to 77 (74.8%) non-widowed faithful pairs and 26 (25.2%) non-widowed divorcing pairs over the same period. As each widowed pair had one focal individual able to be analysed while non-widowed pairs had two, the final number of analysed individuals was 18 faithful widowed, 9 divorcing widowed, 154 faithful non-widowed, and 52 divorcing non-widowed.

### Social behavioural change in faithful widowed birds

When the entire period before the widowing date was compared to the entire period after, no significant difference in feeder visitation rate (coef=-0.067 ± s.e=0.100, *p=*0.503), feeder location (0.386±0.793, 0.626), weighted degree (0.028±0.095, 0.769), opposite sex proportion (-0.014±0.017, 0.412), flock size (-0.354±0.896, 0.693), or top visit adjacency index (0.000±0.142, 1.00) was seen after predicted faithful individuals were widowed. There was still no significant effect in these metrics when only the weekends before and after widowing were analysed. Full coefficient tables and plots for these results are provided in the Supplementary Information.

In the weekend immediately after their partner vanished, widowed birds showed a near significant increase in normalised degree (coef=0.231 ± s.e=0.132, *p=*0.081; Figure 3). However, they showed no similar increase in degree over the entire period after their widowing (0.089±0.097, 0.360).

**Fig 3:**
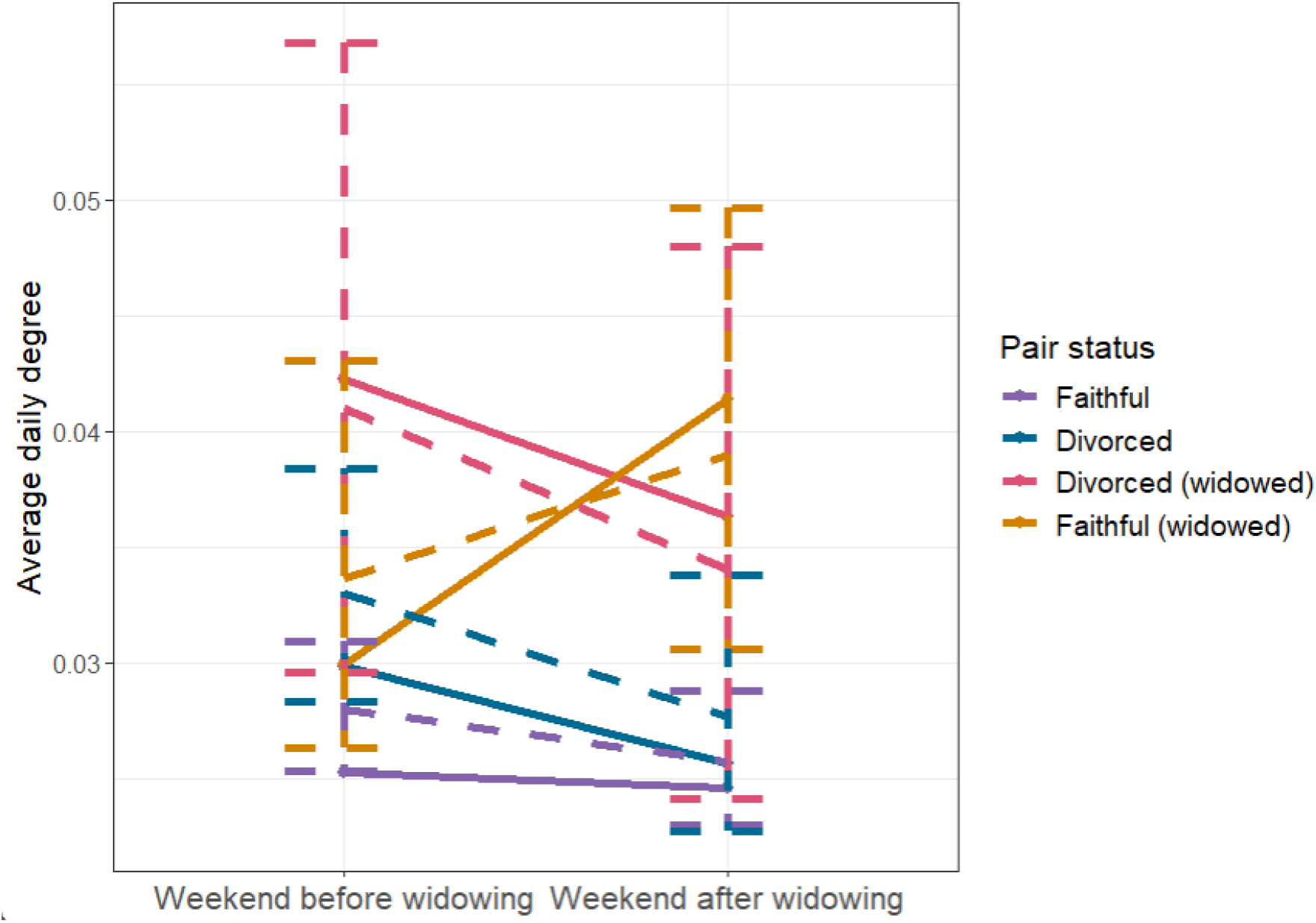
Predicted and observed difference between average daily degree in the weekend before and after the loss of a bird’s mate (widowing). Solid points and lines are averages from the raw data, while dashed error bars and lines are predictions from a log normal generalised linear mixed model, with upper and lower bars representing a 95% confidence interval. Predictions were made using ggeffects, accounting for random effects and a zero-inflation term (Lüdecke, 2018).

Their opposite-sex degree also significantly increased in the weekend immediately after their partner vanished (coef=0.285 ± s.e=0.125, *p=*0.023; Figure 4). As no increase was seen in proportion of opposite sex associates, this increase is likely driven by an increase in associations overall, rather than by a focus on opposite sex interactions.

**Fig 4:**
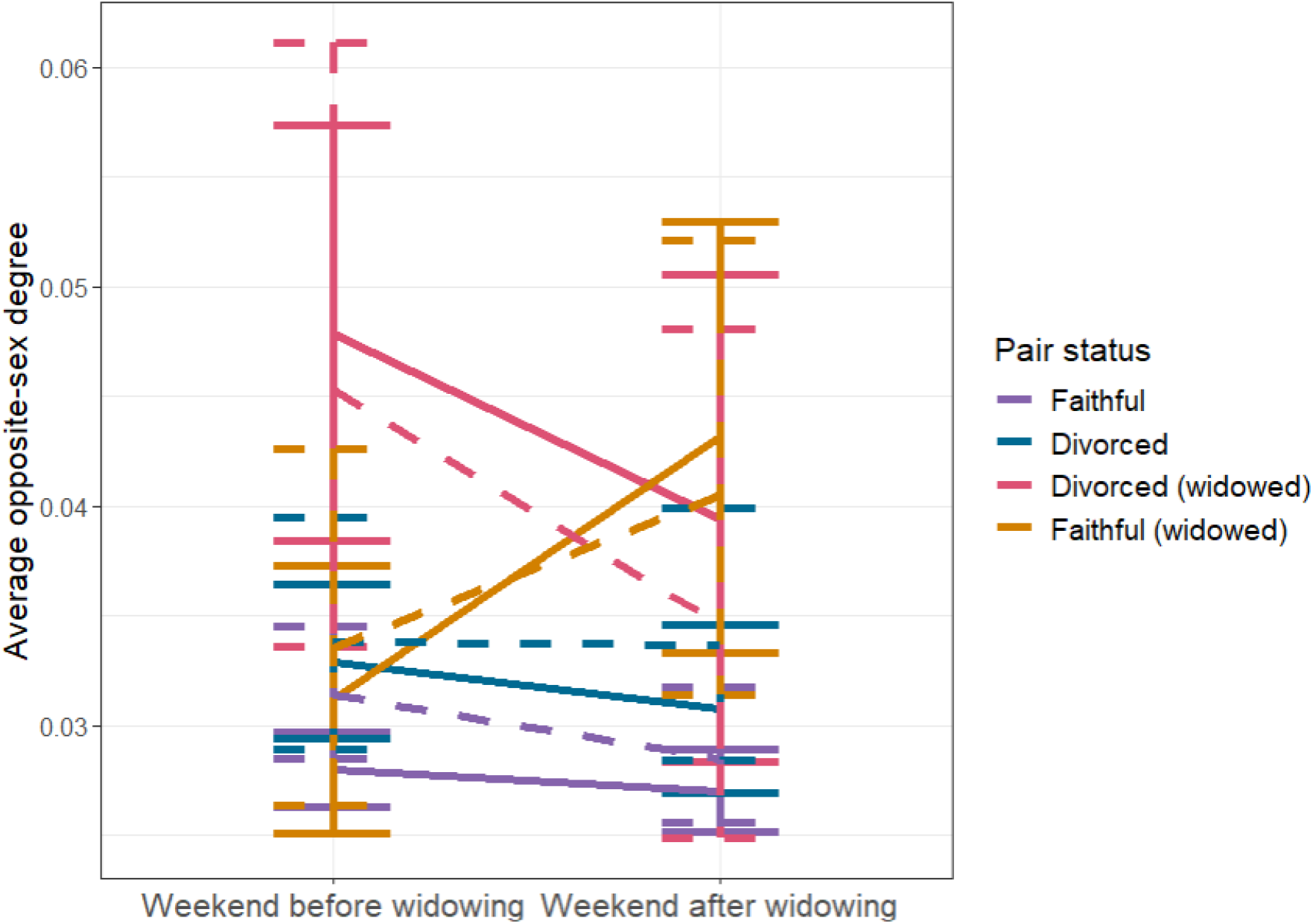
Predicted and observed difference between average opposite-sex degree in the weekend before and after the loss of a bird’s mate (widowing). Solid points and lines are averages from the raw data, while dashed error bars and lines are predictions from a log normal generalised linear mixed model, with upper and lower bars representing a 95% confidence interval. Predictions were made using ggeffects, accounting for random effects and a zero-inflation term (Lüdecke, 2018).

Top winter association score was lower in the post-widowing period for faithful widowed birds in the overall period before and after (coef=-0.191 ± s.e=0.084, p=0.023), but not in the weekend immediately before and after (-0.137±0.122, 0.259). This suggests that the top winter association score decreases after widowing in the long term, especially when contrasted with the lack of a similar pattern in other pair types (Figure 5).

**Fig 5:**
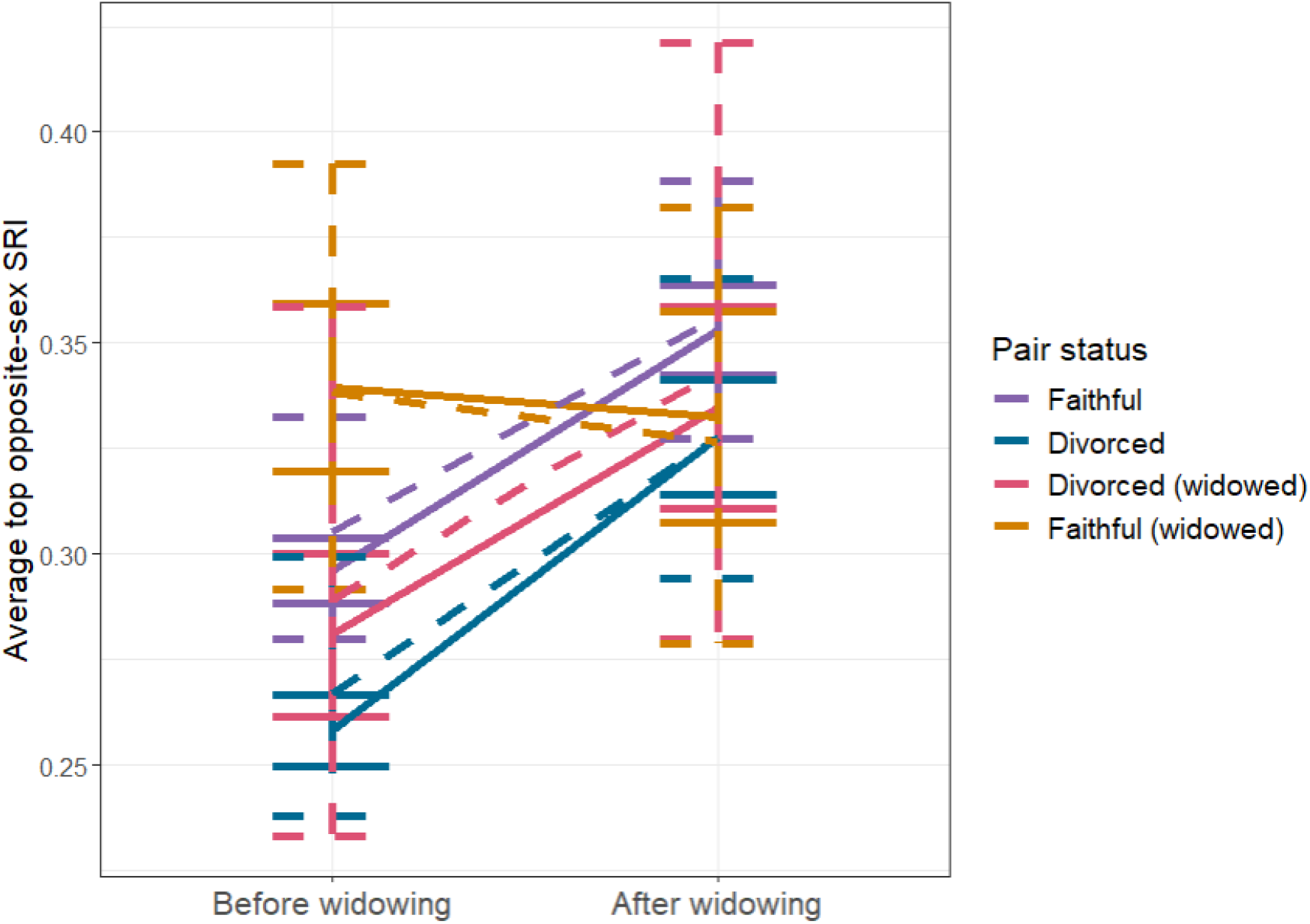
Predicted and observed difference between average top winter association score before and after the loss of a bird’s mate (widowing). Solid points and lines are averages from the raw data, while dashed error bars and lines are predictions from a log normal generalised linear mixed model, with upper and lower bars representing a 95% confidence interval. Predictions were made using ggeffects, accounting for random effects and a zero-inflation term (Lüdecke, 2018).

Analysis of top winter association in the final weekend of data collection shows that any difference in top winter association score between the pair statuses is no longer significant by the end of the experimental period (coef=-0.158 ± s.e=0.099, *p*=0.110; full coefficient table in Supplementary Information).

## Discussion

Within populations of social animals, population turnover means that individuals will often have to experience the loss of social associates. We know little about what form their response to this loss may take, even in cases where the lost associate is a key social counterpart such as a mate. In previous research, wild great tits have shown some response to the loss of an associate (Aplin et al. 2015; Firth et al. 2017). We predicted that a strong response would be observed in cases of individuals losing a previous mate, due to research demonstrating that they may shape their winter foraging decisions around their breeding partners (Firth et al 2015; Abraham, Sheldon, and Firth 2025).

Across the majority of social measures, we saw no significant post-widowing shift in social behaviour, suggesting striking social resilience to these losses. However, there was evidence that birds may increase their number of associates in the weekend immediately after the loss of their mate. This was not driven by an increase in average flock size, proportion of associates which were of the opposite sex, or rate of visits to the supplementary feeders. As this change in behaviour was only observed in the short-term, it suggests an immediate response to the vanishing of a partner. This may indicate an attempt by the widowed individual to buffer the loss of their partner, which could involve trying to develop a new pair bond relationship.

We also found that widowing reduced an individual’s top opposite sex association score over the remainder of the season. If the pair bonding process is impacted in this way, we might expect to see a longer term adjustment of social behaviour to buffer this effect, but this was not observed. Previous research has shown that familiarity with a mate is beneficial for breeding, both directly (possibly through a mechanism of increased coordination/cooperation) and indirectly (via allowing an earlier laydate) (Culina, Firth, and Hinde 2020). Therefore, it may not be valuable for birds to attempt to adjust their social network to meet new potential mates, as meeting their new partner that late could limit breeding success. If they were doing this, we would expect to see social behavioural changes such as a longer-term increase in degree, or a shift towards a network containing more opposite sex conspecifics. Hence, individuals may focus on drawing a new partner from their already existing social network, therefore not adjusting their overall social behaviour after widowing. More detailed individual level analyses are required to investigate this hypothesis.

While we used individual social behavioural measures as our object of analysis here, pair bonding is a dyadic process, requiring two individuals. The shift in winter association score post-widowing may be insurmountable by the individual, particularly if widowing occurs late in the season, when pair bonds may already be developing for many birds. Remaining individuals to pair with could be of lower quality, or it may be more difficult to find a new partner. Further, socially prioritising a new partner to the extent required to raise association score might require the deprioritisation of non-partner social associations. These connections have a wide range of possible benefits, such as information transfer and predation dilution, so losing strength in these connections could have negative effects on the individual. Prioritising a new partner could also leave birds vulnerable to another widowing event, requiring further social network adjustment.

The dyadic nature of pair bonds may also limit our conclusions, as vanishing is likely non-independent within pairs, and a vanishing event may reflect an aspect of behaviour or status in the remaining individual. For example, many disappearances of individuals from the population can probably be attributed to age-related decline, low food availability, or predation (Clobert et al. 1988). Great tits show age assortative mating (Woodman et al. 2023), where individuals are more likely to breed with a mate of a similar age. There is also some evidence of social aging, with adult birds being more repeatable in their social behaviour than juveniles (Aplin et al. 2015). If both of these occur, and older birds show higher mortality, then widowed birds may show different social behaviour than non-widowed by virtue of being older. However, this is unlikely to explain the immediate, short-term effects of widowing. As faithful mates have been shown to associate more often during the winter (Abraham, Sheldon, and Firth 2025), they are likely to experience similar environmental effects, such as food availability. This is a further confounder, which may influence behaviour at the feeders in a similar way for both a vanished bird and its living partner, despite the living individual’s survival. Finally, experiencing a near-miss predation event has been shown to change tit social behaviour in the short term (Voelkl, Firth, and Sheldon 2016). If a bird experiences the predation event which leads to the vanishing of its mate, it could explain short-term behavioural effects such as the ones we observed.

An explanation for the lack of long-term behavioural response despite the difference in winter association score is that it may not be impactful enough on breeding success to select for a potentially costly social behavioural response. As seen, the difference between association scores at the end of the data collection period (as winter draws to a close) is no longer statistically significant between pair types. There are also weeks between the end of data collection and the beginning of breeding proper for pairs to close this gap in association scores. Additionally, top winter association score refers to the partner of the individual - that is, their top winter associate, who they may not go on to mate with. For 44.5% of faithful widowed birds, their winter partner (excluding their vanished mate) does not go on to become their next mate (calculated using data from this analysis). There are likely aspects of the post-widowing pair bonding process that are not being picked up by a focus on winter partner and association scores alone. For instance, spatial behaviour, nestbox choice and the complexity of dyadic mate choice. This provides an exciting direction for further analysis, investigating whether and how winter partners become breeding pairs, and if this is impacted by events such as widowing.

As an interesting addition to previous work in this system (Firth et al. 2017; Abraham, Sheldon, and Firth 2025) and other systems considering pair bonded individuals (Duckworth 1992; Thomas and Wolff 2004), wild great tits showed only a short-term increase in degree after the loss of a breeding mate, but were less close with their post-widowing partner during the non-breeding season. In order to understand how birds respond after being widowed, future analysis could investigate in more detail the process of how birds select their new mate, and the potential fitness consequences in relation to the timing of experiencing widowing, as well as possible non-independence in pair mortality. This will provide more insight into how individuals in natural social systems respond to the loss of others, helping us to predict how populations respond to change.

