## Supplementary Information for "Wild great tits show social resilience in response to partner loss"

Calculating behavioural metrics

Estimating accuracy of vanishing

Models without final weekend (78/79 vanishers)

Status prediction information and sensitivity analysis

Full models

Sensitivity of non-widowed ‘widowing date’

Split sex models

#### Calculating behavioural metrics

Feeder visitation rate

We calculated feeder visitation rate before and after the widowing date. This was calculated as the average feeder visits per experimental day for each relevant period (including days where the bird was not observed in the average). A 1:n overdispersion random effect was included in the models for feeder visitation rate, due to non-normality in the residuals.

Preferred feeder location

In case the birds responded spatially while maintaining the same overall social structure, we investigated whether preferred feeder location changed before and after widowing date. For preferred feeder location, we used the feeder which an individual visited most often for each period.

This measure was modelled as binomial, with the response variable being whether a bird did or did not change its preferred location after widowing, and pair status being the only fixed effect. Year and ID were still included as random effects.

Daily degree

We calculated daily degree as one measure of general social behaviour. Degree measures the number of associations an individual has (Zhang and Luo 2017). We assumed that widowing may lead to a change in daily degree if birds responded to widowing by increasing their number of associations, in order to increase their pool of potential replacement partners. Degree was calculated as the number of birds an individual was observed associating with on a day, scaled by the number of birds observed in total on that day.

Weighted degree

Weighted degree was also used, which is the weighted sum of the winter association score of an individual’s associations to all its social partners. Previous research into social responses to associate loss had shown a response only in weighted degree, which may reflect a change in association intensity beyond any change in association numbers (Firth et al. 2017). That is, in response to a loss, individuals may strengthen their connections to existing associations, without forming more.

Opposite sex degree

We also calculated opposite sex degree, which is the number of associations an individual has with birds of the opposite sex, normalised by the number of birds of the opposite sex observed during that day. This was included for similar reasons to weighted and daily degree, but in order to assess whether birds are specifically changing their social behaviour in regards to potential mates.

Opposite sex proportion

Opposite sex proportion was included as an alternative measure of the extent to which an individual was prioritising connections to potential mates. It was calculated as the number of opposite sex connections as a proportion of total connections. Some birds have unknown sex, and these were included as total associations, but not as opposite sex associations.

Flock size

Average flock size before and after widowing was used to investigate the way in which birds might adjust their social environment. For example, if a bird was increasing its number of associates, it could do this both through increasing the number of flocks it interacted with, or by increasing the size of those flocks. Flock size was simply calculated as the average number of birds observed in flocks where the focal individual was seen.

Top winter association score

To investigate whether widowing had an impact on strength of association with a bird’s partner, we investigated individuals’ top winter association score with opposite sex associates in the period before and after widowing.

As top winter association score was only calculated with opposite sex associates, it was recorded as a zero if an individual was observed at the feeders but without any opposite sex associates. If an individual was not observed at the feeders at all in a given period, top winter association score was recorded as NA. Birds were only counted as opposite sex if their sex was known, so birds with unknown sex were not included in these calculations.

Top visit adjacency index

Previous work (Firth et al. 2015; Abraham, Sheldon, and Firth 2025) has shown that following behaviour might be relevant as an indicator of pair bonding at a smaller spatiotemporal scale than shared flocking events (i.e winter association score). For this reason we also calculated individuals’ top visit adjacency index with opposite sex associates in the period before and after.

Visit adjacency index represents the proportion of visits to the feeders within 3 seconds of another bird. It can be represented mathematically as:

$$VAI = \frac{x}{x + y}$$

where *x* is a visit within 3 seconds of a given opposite sex associate and *y* is all other recorded visits.

For each focal individual, we calculated VAI with all opposite sex associates, then selected the top value of VAI. Birds were only counted as opposite sex if their sex was known, so birds with unknown sex were not included in these calculations.

#### Estimating accuracy of vanishing

In this analysis, ‘widowing’ was used to refer to an individual no longer being present in the population. This was defined through a combination of not being observed at the feeders for the remainder of the winter, and not being observed to breed in any later breeding seasons. However, in both of these measures it is possible for a bird which is still present in the population to go unobserved, and therefore be incorrectly recorded as vanished. Here, we try to estimate the accuracy of our vanishing dates (Figure A1).

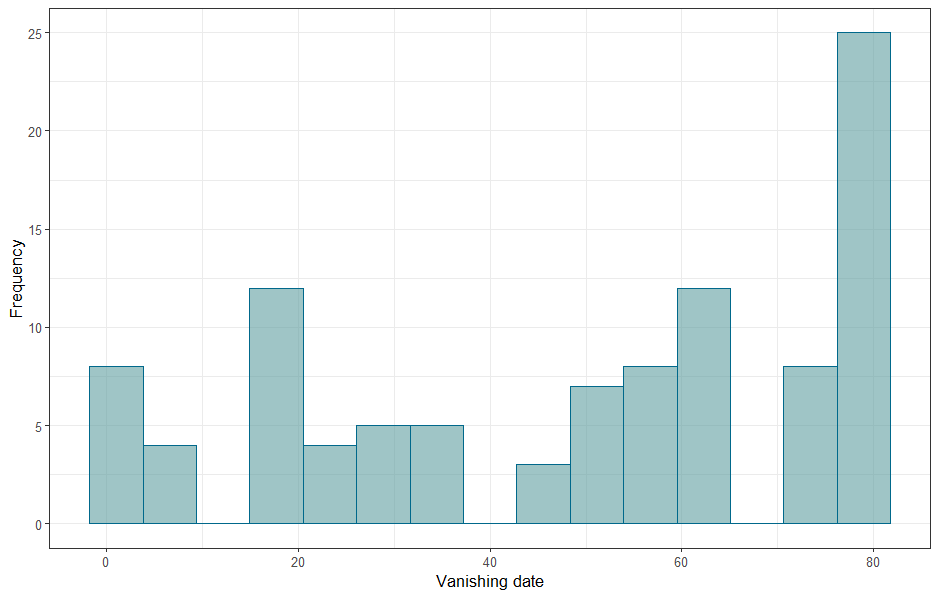

*Figure A1: Estimated vanishing date (last day observed during the winter) for the vanished birds identified in this analysis.*

The majority of birds did not visit the feeders every day (Figure A2). Therefore, particularly in vanishing dates calculated as late in the winter, such as the seemingly inflated number of birds whose last day observed was day 79 (28 (24.6%)), the individual not being observed again could be due to chance based on their visitation rate.

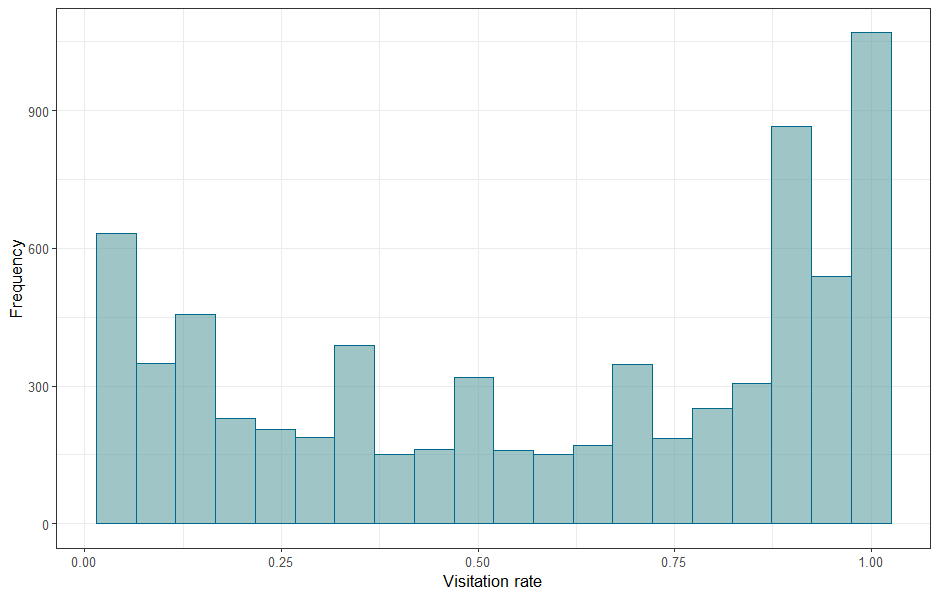

*Figure A2: The visitation rate of all observed birds to the feeders, measured as the proportion of days during the data collection period where they were observed at least once.*

For each individual classed as vanished, we calculated their visitation rate up to their vanishing date (Figure A3). The distribution of visitation rates is different from the distribution for all birds in the population, without a less strong bimodal shape. This might be due to visitation rates being calculated for these birds only before vanishing date, or because of the selection criteria of having been observed in a breeding pair the year before, therefore removing new immigrants and juveniles from the analysis.

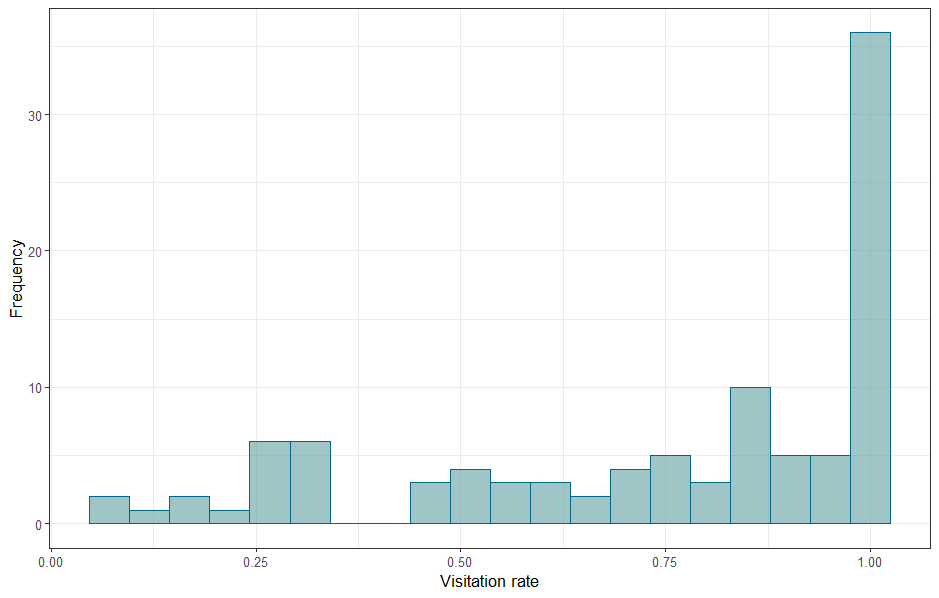

*Figure A3: The visitation rate of all vanished birds to the feeders, measured as the proportion of days before their final observed day where they were observed at least once.*

We then treated their visitation rate as a binomial probability, representing the chance that they would visit the feeders on a given day. This probability was then used to calculate the chance that they would have not visited the feeders for every day after their calculated vanishing date, given their previous visitation rate. For example, if a bird’s vanishing date was calculated as day 58, and its previous visitation rate was 0.85, the chance of it not being observed for every day after day 58 would be 1.708594e-06, so we can be fairly confident of our vanishing date estimate. The results of this analysis for each individual (Figure A4) showed very low probabilities of that length of vanishing being observed by chance for most birds, although two birds had greater than 10% chance of being unobserved for that long by chance, given their previous visitation rate. These birds were not concentrated at a particular vanishing date.

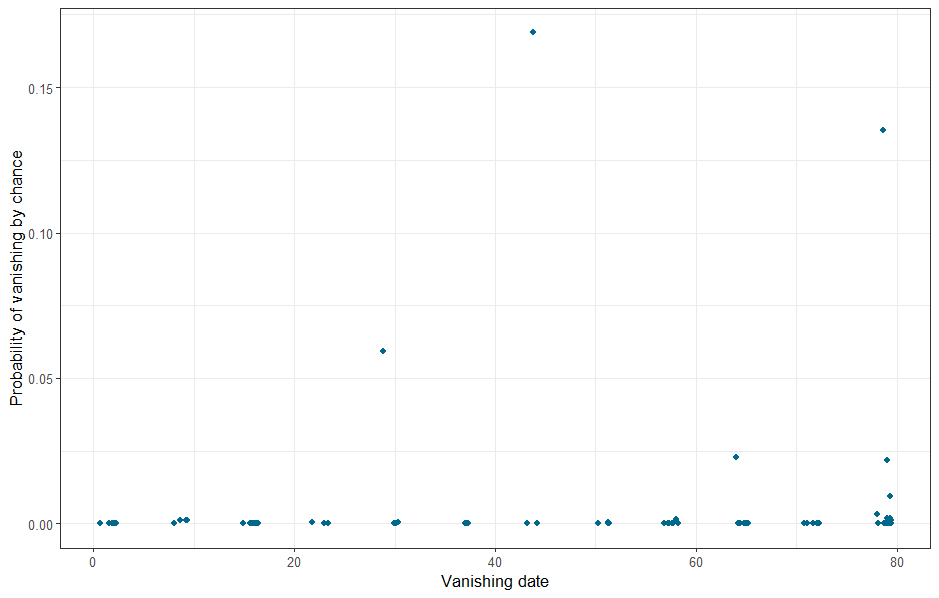

*Figure A4: The binomially estimated probability of a bird having not been observed at the feeders by chance, plotted against our estimated vanishing date. Points are jittered.*

To check the above results, we compared them to the same calculations done for birds which we know to be alive (were observed breeding later), but which also vanished at some point in the winter. The vanishing dates for these birds (Figure A5) were similar for those observed in the birds used in our analysis, including a peak at day 16, suggesting a possible environmental driver for birds stopping coming to the feeders.

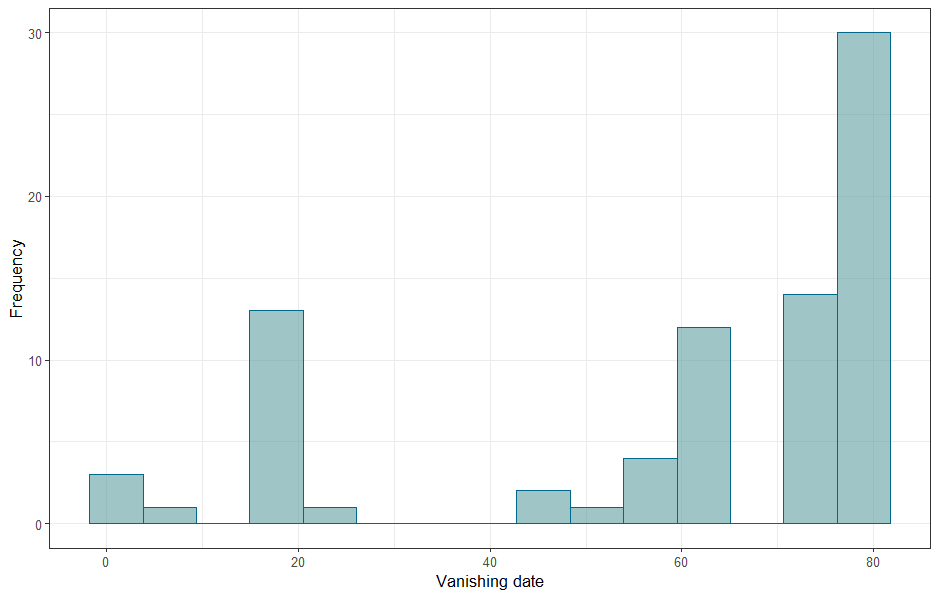

*Figure A5: The estimated vanishing date for birds which ‘vanished’, but went on to later be observed breeding.*

The distribution of probabilities for these results being observed by chance (Figure A6) is also very similar to that observed in birds which we considered to truly have vanished. This suggests that our confidence in the accuracy of our vanishing dates may be inflated, and that stopping visiting the feeders does not necessarily suggest death. However, it does not mean that the birds included in our analysis did not truly vanish then, as a disappearance from the feeders is likely to reflect a true change in behaviour, whether or not a bird has died. It does raise the possibility that our inability to detect a behavioural change in widows may be due to an inability to accurately identify widowed birds or true vanishing date.

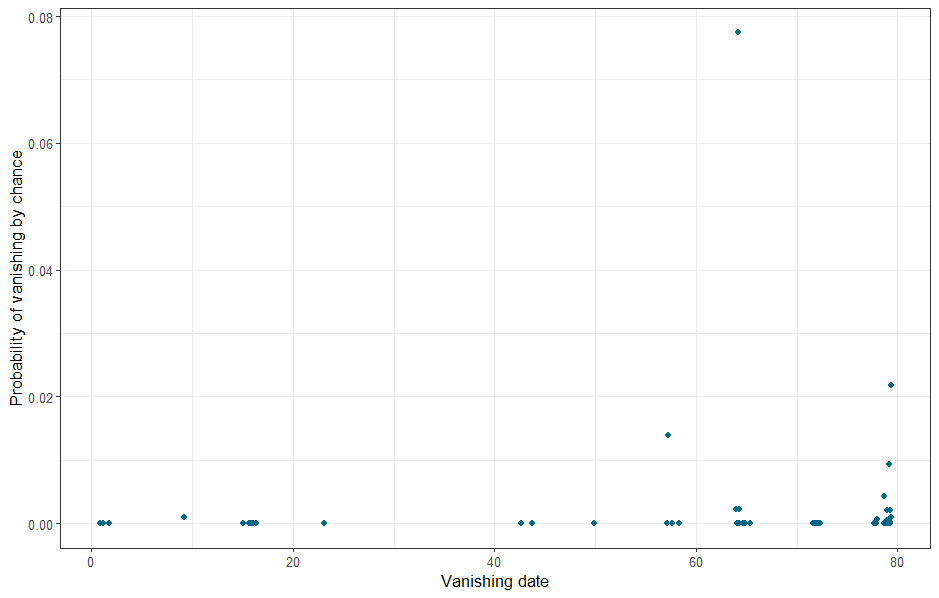

*Figure A6: The binomially estimated probability of a bird having not been observed at the feeders by chance, plotted against our estimated vanishing date, for birds which ‘vanished’ but were later observed breeding. Points are jittered.*

The fact that widowed birds failed to appear during the breeding season is not meaningless, but only a minority of birds in this dataset were observed breeding again the year after their first attempt (447 (35.7%)). Many of these birds will have died at some point between breeding seasons, making their partner truly widowed. However, data collection during the breeding season has some error, allowing some birds to go unidentified even during a breeding attempt. Assuming that all great tits in the woods choose to breed in the provided nestboxes (see Perrins 1979 for an estimate of the low numbers of birds choosing to breed in natural cavities), there is still room for adults to die or nests to fail before the point of adult identification, or for them to fail to be identified at all. Due to the use of RFID readers, it is less likely for previously tagged birds to not be identified if they reach the adult identification stage, but still possible. Kidd et al. 2015 found an early failure rate of 17% in their sample. However, this rate was highest in young unringed immigrants, which would not have been included in our analysis. This can make us more confident in a bird not being observed during the breeding season.

There may also be yearly differences in confidence. The distribution of vanishing dates for each year varies between vanished birds used in our analysis (Figure A7), and those ‘false’ vanishers who reappear in the breeding season (Figure A8), despite the overall distribution of vanishing dates being similar. For example, the peak of vanishings at day 16 is formed across all three years for ‘true’ vanishers, but only present in 2013 for ‘false’ vanishers. The majority of ‘false’ vanishers are concentrated in 2012, in the last few weekends. There is a similar pattern in ‘true’ vanishers, but it is less visually notable when compared to other years.
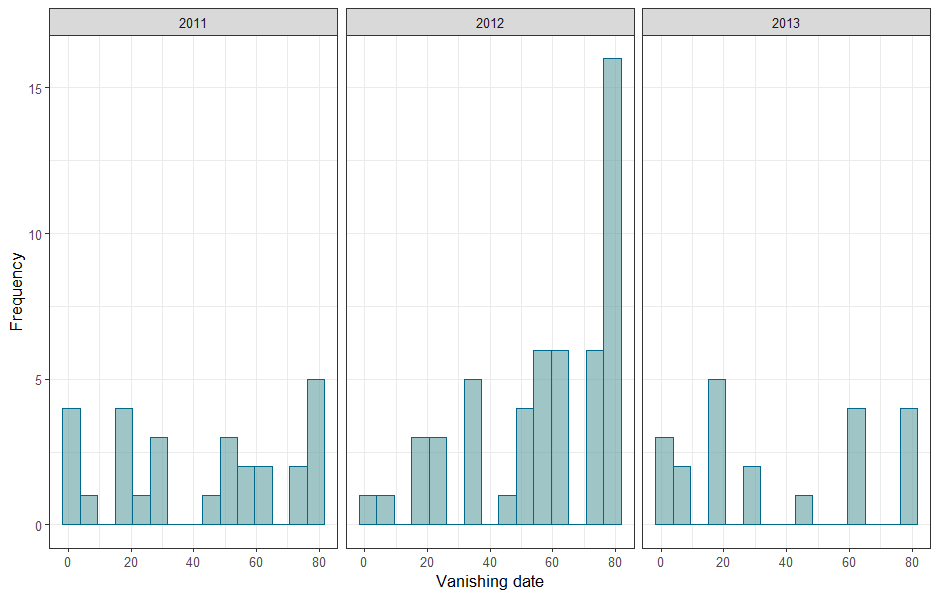

*Figure A7: Vanishing date distribution across the years for birds which vanished at some point in the winter, and were not observed again during the breeding season (‘true’ vanishers).*

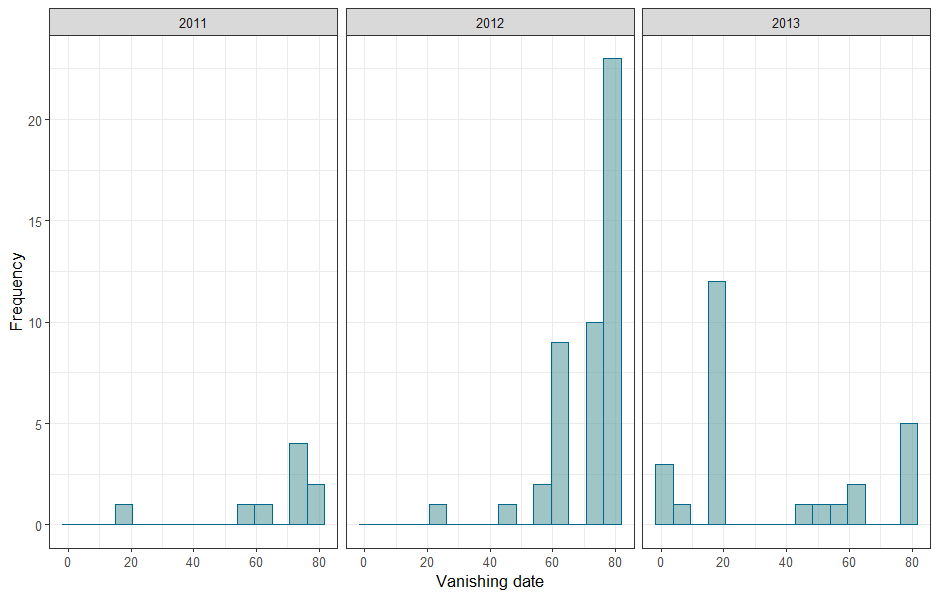

*Figure A8: Vanishing date distribution across the years for birds which vanished at some point in the winter, and were observed again during the breeding season (‘false’ vanishers).*

Based on a simple linear model of the effect of an interaction between year and day on observation number, the numbers of observations (feeder visits) are significantly different between years, with significantly different trends (Figure A9). However, if this was the full explanation for the trends observed in vanishing dates, we would expect there to also be higher vanishing dates in the final year, which has lower observations than both other years.

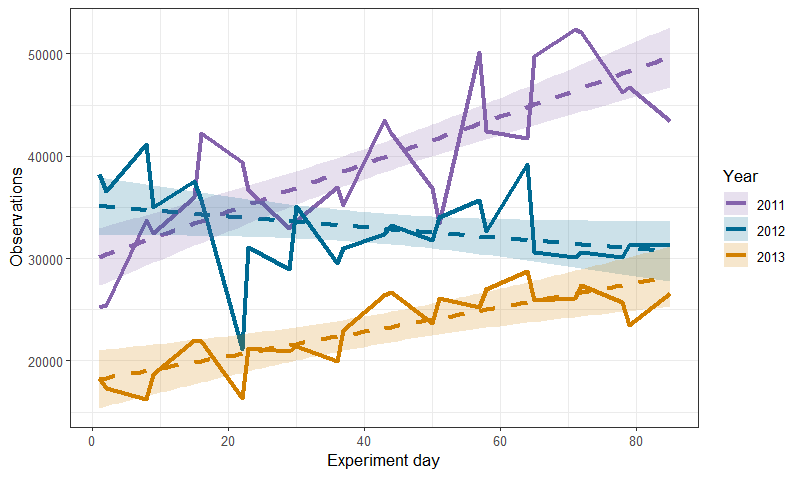

*Figure A9: Predicted and raw data for the number of observations at the feeders across three winters. Solid lines are the real numbers of observations, while dashed lines and ribbons are the predicted values from a linear model and 95% confidence intervals around the predictions.*

Retention rate (the chance of a bird being observed again in the breeding season if it was seen in the preceding winter) was significantly lower in 2012 (coef=-0.142±0.068, *p*=0.037) than 2011, and significantly higher in 2013 (0.343±0.065, <0.001) (Figure A10).

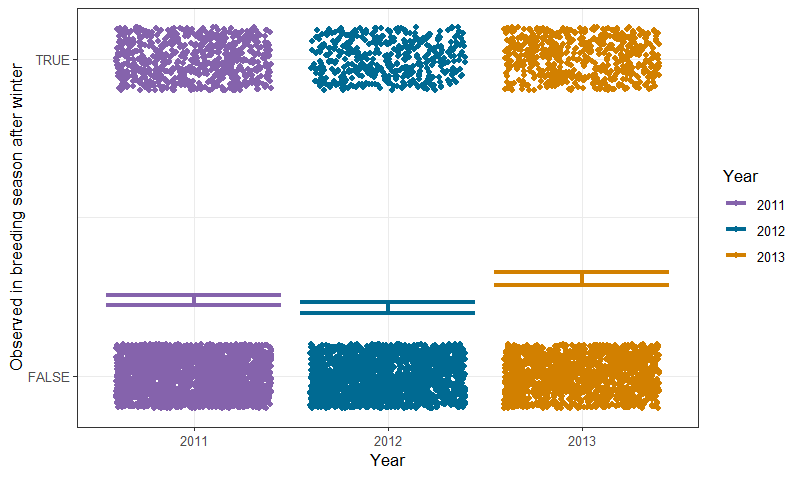

*Figure A10: Predicted and raw values from a binomial generalised linear model of the chance of a bird being observed in both a winter and the following breeding season. Points represent individual birds, while error bars are the 95% confidence interval of the model estimate.*

With only three years of data, it is hard to draw conclusions from these patterns in vanishing date. However, it is interesting that this distinct pattern of behaviour at the feeder (i.e vanishing) may relate to a behavioural change caused by something which appears to vary between years.

Another way to investigate our confidence in the accuracy of our vanishing estimates is to account for the potential non-independence of vanishing. In whales, Whitehead and Gero (2013) showed improved mortality estimates when accounting for whether an individual’s close associates also vanished from the population - if they did, it suggested a change in behaviour for the group rather than the death of an individual.

By dividing birds into groups based on their appearance or disappearance in different parts of the season, we can start to investigate non-independent patterns of disappearance. The groups are as follows:

- Alive: Seen breeding in two consecutive springs.
- Vanished before winter: Seen breeding, then not seen in the following winter, or any further breeding seasons.
- Vanished during winter: Seen breeding, then at least once in the winter, but stopped visiting feeders at some point during the winter, and is not observed in following breeding seasons.
- Vanished after winter: Seen breeding, then during the entire winter, then not observed at following breeding seasons.
- ‘False’ vanish: Seen breeding, then at least once in the winter, but stopped visiting feeders at some point during the winter, then was observed in following breeding seasons.

As shown in Figure A11, the distribution of mate statuses differs between ‘true’ and ‘false’ vanishers. Particularly, the birds who vanish in the winter and are not observed in the breeding season are far less likely to have a mate who also vanishes, but is then observed breeding later. If we can assume a similar non-independence as was observed in Whitehead and Gero (2013), this suggests a meaningful difference between birds who vanished, only to reappear, and birds who did not reappear.

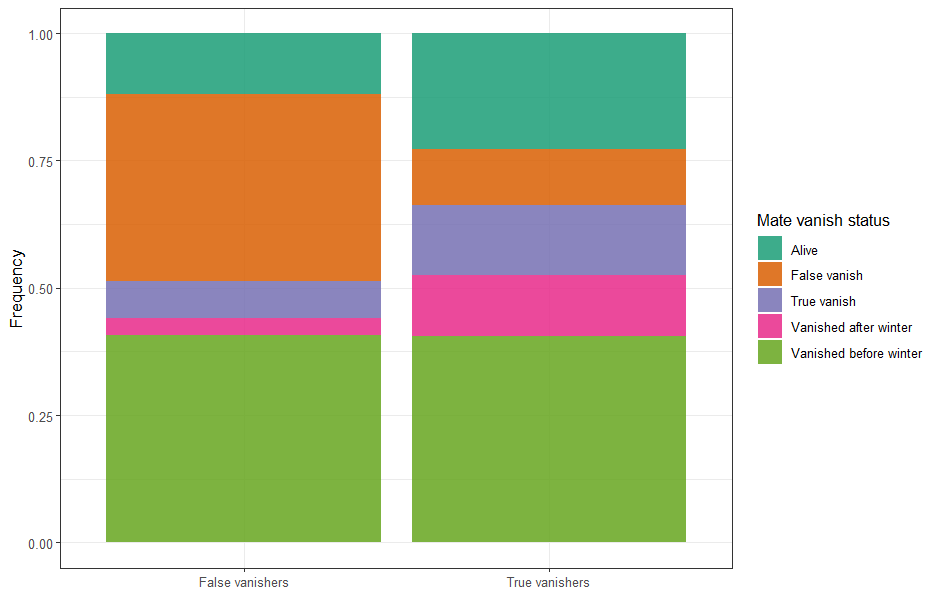

*Figure A11: The vanishing status of the mates of birds who vanished in the winter, but were observed breeding the following spring (‘True’ vanishers), contrasted with birds who vanished in the winter, and were not observed breeding again (‘False’ vanishers).*

#### Models without final weekend (78/79 vanishers)

The most common vanishing date in the dataset was day 79. For widows of birds estimated as vanishing on day 78 or 79, there was only one weekend of data after the vanishing for any changes in behaviour to be observed. In case this impacted the models, we repeated models from the main body of the text but only with birds who vanished before days 78/79, including re-generating the fake vanishing dates of non-widowed birds. We only ran the models analysing the entire period before and after widowing, as the models only analysing the weekend before and after had the same amount of pre- and post-widowing data for all birds, regardless of vanishing date. All models without the late vanishing birds supported the conclusions from the reported models (coefficient tables below).

Daily degree - period before/after

*Table A1: Fixed effect coefficient estimates from a lognormal GLMM of the effect of pair status and widowing on daily degree, without pairs with a widowing date of 78 or 79. Significant coefficients are indicated in italics.*

| Coefficient | Estimate | Standard error | *p*-value |
| --- | --- | --- | --- |
| *Intercept* | *-3.813* | *0.047* | *<0.001* |
| Pair status Divorced | 0.011 | 0.073 | 0.876 |
| Pair status Divorced (widowed) | 0.045 | 0.182 | 0.806 |
| Pair status Faithful (widowed) | 0.174 | 0.127 | 0.169 |
| Post-widowing | 0.052 | 0.035 | 0.133 |
| *Pair status Divorced:Post-widowing* | *0.142* | *0.070* | *0.043* |
| Pair status Divorced (widowed):Post-widowing | 0.264 | 0.181 | 0.145 |
| Pair status Faithful (widowed):Post-widowing | 0.088 | 0.097 | 0.362 |

Weighted degree - period before/after

*Table A2: Fixed effect coefficient estimates from a lognormal GLMM of the effect of pair status and widowing on weighted degree, without pairs with a widowing date of 78 or 79. Significant coefficients are indicated in italics.*

| Coefficient | Estimate | Standard error | *p*-value |
| --- | --- | --- | --- |
| *Intercept* | *2.099* | *0.071* | *<0.001* |
| Pair status Divorced | 0.055 | 0.070 | 0.431 |
| Pair status Divorced (widowed) | -0.063 | 0.186 | 0.735 |
| Pair status Faithful (widowed) | 0.133 | 0.112 | 0.238 |
| *Post-widowing* | *0.074* | *0.032* | *0.020* |
| Pair status Divorced:Post-widowing | 0.071 | 0.062 | 0.252 |
| Pair status Divorced (widowed):Post-widowing | 0.237 | 0.150 | 0.115 |
| Pair status Faithful (widowed):Post-widowing | 0.042 | 0.091 | 0.644 |

Opposite sex degree - period before/after

*Table A3: Fixed effect coefficient estimates from a lognormal GLMM of the effect of pair status and widowing on opposite sex degree, without pairs with a widowing date of 78 or 79. Significant coefficients are indicated in italics.*

| Coefficient | Estimate | Standard error | *p*-value |
| --- | --- | --- | --- |
| *Intercept* | *-3.528* | *0.044* | *<0.001* |
| Pair status Divorced | 0.006 | 0.065 | 0.923 |
| *Pair status Divorced (widowed)* | *0.439* | *0.141* | *0.002* |
| Pair status Faithful (widowed) | 0.110 | 0.105 | 0.297 |
| Post-widowing | 0.001 | 0.030 | 0.972 |
| *Pair status Divorced:Post-widowing* | *0.175* | *0.056* | *0.002* |
| Pair status Divorced (widowed):Post-widowing | 0.040 | 0.105 | 0.711 |
| Pair status Faithful (widowed):Post-widowing | 0.050 | 0.084 | 0.552 |

Opposite sex proportion - period before/after

*Table A4: Fixed effect coefficient estimates from a Gaussian GLMM of the effect of pair status and widowing on opposite sex proportion, without pairs with a widowing date of 78 or 79. Significant coefficients are indicated in italics.*

| Coefficient | Estimate | Standard error | *p*-value |
| --- | --- | --- | --- |
| *Intercept* | *0.375* | *0.008* | *<0.001* |
| Pair status Divorced | -0.015 | 0.012 | 0.196 |
| Pair status Divorced (widowed) | 0.010 | 0.030 | 0.748 |
| Pair status Faithful (widowed) | -0.007 | 0.019 | 0.723 |
| Post-widowing | 0.006 | 0.005 | 0.232 |
| Pair status Divorced:Post-widowing | 0.017 | 0.010 | 0.107 |
| Pair status Divorced (widowed):Post-widowing | 0.002 | 0.026 | 0.935 |
| Pair status Faithful (widowed):Post-widowing | -0.013 | 0.019 | 0.480 |

Average flock size - period before/after

*Table A5: Fixed effect coefficient estimates from a Gaussian GLMM of the effect of pair status and widowing on opposite sex proportion, without pairs with a widowing date of 78 or 79. Significant coefficients are indicated in italics.*

| Coefficient | Estimate | Standard error | *p*-value |
| --- | --- | --- | --- |
| *Intercept* | *11.720* | *0.854* | *<0.001* |
| Pair status Divorced | 0.463 | 0.587 | 0.437 |
| *Pair status Divorced (widowed)* | *3.945* | *1.518* | *0.010* |
| Pair status Faithful (widowed) | 1.760 | 0.967 | 0.069 |
| Post-widowing | 0.073 | 0.269 | 0.787 |
| *Pair status Divorced:Post-widowing* | *1.306* | *0.529* | *0.014* |
| Pair status Divorced (widowed):Post-widowing | -0.268 | 1.311 | 0.838 |
| Pair status Faithful (widowed):Post-widowing | -0.516 | 0.941 | 0.583 |

Feeder visitation rate - period before/after

*Table A6: Fixed effect coefficient estimates from a lognormal GLMM of the effect of pair status and widowing on feeder visitation rate, without pairs with a widowing date of 78 or 79. Significant coefficients are indicated in italics.*

| Coefficient | Estimate | Standard error | *p*-value |
| --- | --- | --- | --- |
| *Intercept* | *2.591* | *0.072* | *<0.001* |
| Pair status Divorced | 0.101 | 0.080 | 0.207 |
| Pair status Divorced (widowed) | -0.106 | 0.214 | 0.621 |
| Pair status Faithful (widowed) | 0.209 | 0.127 | 0.101 |
| *Post-widowing* | *0.100* | *0.031* | *0.002* |
| Pair status Divorced:Post-widowing | 0.037 | 0.057 | 0.524 |
| Pair status Divorced (widowed):Post-widowing | 0.227 | 0.173 | 0.188 |
| Pair status Faithful (widowed):Post-widowing | -0.005 | 0.088 | 0.959 |

Preferred feeder location - period before/after

*Table A7: Fixed effect coefficient estimates from a Bernoulli GLMM of the effect of pair status on the chance of changing feeder location after widowing, without pairs with a widowing date of 78 or 79. Significant coefficients are indicated in italics.*

| Coefficient | Estimate | Standard error | *p*-value |
| --- | --- | --- | --- |
| *Intercept* | *1.792* | *0.248* | *<0.001* |
| Pair status Divorced | -0.483 | 0.434 | 0.265 |
| Pair status Divorced (widowed) | -0.182 | 1.123 | 0.871 |
| Pair status Faithful (widowed) | -0.182 | 0.813 | 0.823 |

Top winter association score - period before/after

*Table A8: Fixed effect coefficient estimates from a lognormal GLMM of the effect of pair status and widowing on top winter association score, without pairs with a widowing date of 78 or 79. Significant coefficients are indicated in italics.*

| Coefficient | Estimate | Standard error | *p*-value |
| --- | --- | --- | --- |
| *Intercept* | *-1.238* | *0.036* | *<0.001* |
| *Pair status Divorced* | *-0.157* | *0.049* | *0.002* |
| Pair status Divorced (widowed) | -0.055 | 0.114 | 0.628 |
| *Pair status Faithful (widowed)* | *0.182* | *0.072* | *0.011* |
| *Post-widowing* | *0.148* | *0.024* | *<0.001* |
| Pair status Divorced:Post-widowing | 0.072 | 0.050 | 0.151 |
| Pair status Divorced (widowed):Post-widowing | 0.018 | 0.111 | 0.871 |
| *Pair status Faithful (widowed):Post-widowing* | *-0.187* | *0.077* | *0.015* |

Top visit adjacency index - period before/after

*Table A9: Fixed effect coefficient estimates from a lognormal GLMM of the effect of pair status and widowing on top visit adjacency index, without pairs with a widowing date of 78 or 79. Significant coefficients are indicated in italics.*

| Coefficient | Estimate | Standard error | *p*-value |
| --- | --- | --- | --- |
| *Intercept* | *-6.183* | *0.057* | *<0.001* |
| Pair status Divorced | 0.046 | 0.065 | 0.478 |
| Pair status Divorced (widowed) | -0.074 | 0.133 | 0.579 |
| Pair status Faithful (widowed) | -0.035 | 0.095 | 0.716 |
| Post-widowing | -0.070 | 0.046 | 0.129 |
| Pair status Divorced:Post-widowing | -0.012 | 0.090 | 0.169 |
| Pair status Divorced (widowed):Post-widowing | 0.023 | 0.194 | 0.904 |
| Pair status Faithful (widowed):Post-widowing | 0.000 | 0.142 | 1.000 |

#### Status prediction information and sensitivity analysis

In order to predict faithfulness of widowed pairs, we built models using a test/train procedure to assess accuracy on pairs for which their faithfulness status was known. While a more accurate model could likely be produced with a more extensive model building procedure and variable exploration, we considered the accuracy achieved by our model acceptable given that predicting faithfulness/divorce was not the core goal of this analysis. A sensitivity analysis is provided below to investigate how inaccuracies in the model might have affected the results of the analysis.

As widowing occurred at different dates throughout the winter, we wanted an estimate of accuracy with different amounts of data, so tested the model on censored data from the known status pairs. The model building and running procedure went as follows:

- For a dataset of 103 pairs with known breeding status (faithful/divorced), we calculated the following variables of interest:
  - Clutch size (number of eggs laid by the pair)
  - Mean of the first day the male and the female were observed in the winter season following their first breeding attempt
  - Mean of the distance between previous nestbox and preferred supplementary feeder for the male and the female
  - Mean of the relative association index rank for the male and the female (i.e 1 for the male would mean the female was his closest associate, and 0.5 for the female would mean the male was halfway ranked in her associates, then the two values are averaged)
  - The mean coefficient of variation (not including zeroes) of relative association index rank for the male and female. This number represents how variable the ranks of the birds were to each other
  - The minimum daily difference between relative association index rank of the male and the female across the time period analysed
  - A factor set as TRUE when one or both individuals had their mate as their closest social associate, and FALSE when neither did.
- A test/train procedure across different potential widowing dates was then run as follows, in order to estimate accuracy:
  - Filter social network data to only include up to a given day (beginning at day 2, as day 1 does not have enough data to provide variation estimates).
  - Recalculate the variables above for each of the known status pairs which have enough data up to that day to be included, using the social data from the specified time frame.
  - Run test/train for each known status pair
    - Remove pair 1 from the data frame
    - Run a Bayesian generalised mixed model with binary response term on the remaining pairs, using the above variables as fixed effects with Normal(0,1) priors.
    - Use the model to predict the status of pair 1 100 times
    - Record the mean of those predictions.
      - The mean is a number from 0 to 1, where 1 is a pair always being assigned faithful and 0 is a pair always being assigned divorced.
      - Results were recorded at different levels of confidence, where 50% confidence means every pair given a value above 0.5 is assigned faithful, 60% confidence means every pair above 0.6 is assigned faithful, and so on.
    - Repeat for each pair
  - Using results from the test/train, estimate accuracy of the model for pairs in the given time frame.
  - Repeat for each day of data collection.
- Using the accuracy results, select the confidence level that will be used in the prediction for widowed pairs (we selected 60% confidence).
- The following prediction procedure was then performed on the widowed pairs to estimate their faithfulness status:
  - Filter social network data up to the vanishing date for the widowed pair.
  - Recalculate the variables above for each of the known status pairs which have enough data up to that day to be included, using the social data from the specified time frame.
  - Run the model on the known status pairs.
  - Use the model results to predict the faithfulness status for the widowed pair.
  - Repeat for each widowed pair.

As can be seen below (Figure A12), accuracy at all confidence levels increased with the inclusion of more days of data, which is to be expected, except for the ‘All faithful’ level, which is the accuracy achieved when all pairs are assigned faithful by default. The reason that the accuracy of all faithful changes over time is that more pairs are included over time, as some pairs don’t have enough data from the early winter to be included in the modelling.

We selected the 60% confidence level to use in our predictions of faithfulness in widowed pairs, as it consistently outperformed 50% confidence, and reduced Type 2 error (birds incorrectly being assigned as faithful). While 70% did outperform 60% confidence at times, and had reduced Type 2 error, we didn’t want to be overly strong in our selection criteria, as the data may take a slightly different form in the widowed pairs. The output of the predictions for the widowed pairs also seemed to have a natural break at ~60% (Figure A13), so we took that as a further argument to use 60% confidence as the cutoff.

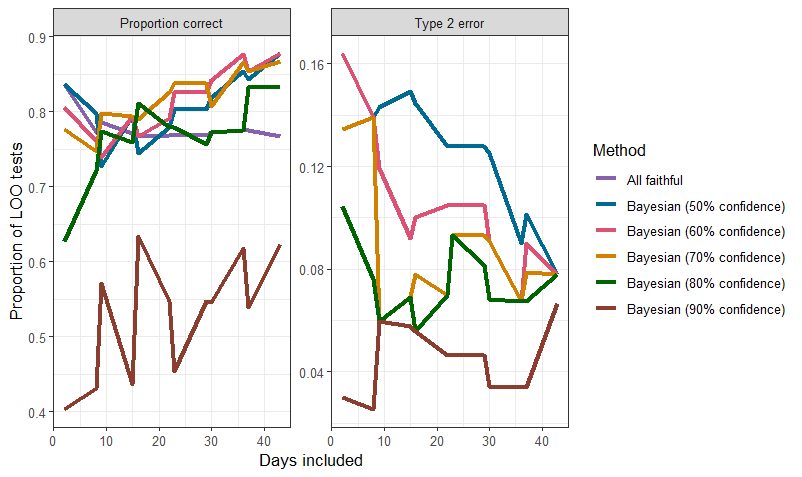

*Figure A12: Proportion of leave-one-out tests which assigned pairs correctly (proportion correct) or incorrectly classified divorced pairs as faithful (Type 2 error). Results were generated using a Bayesian model to predict pair status, with ‘Method’ being the requirement for a pair to be classified as faithful.*

Figure A13 shows a histogram of prediction values for the 27 widowed pairs included in the analysis. Using the 60% confidence cutoff, 9 (33.33%) pairs were assigned divorced status. This is higher than the divorce rate of 25.25% in pairs of known faithful/divorced status.

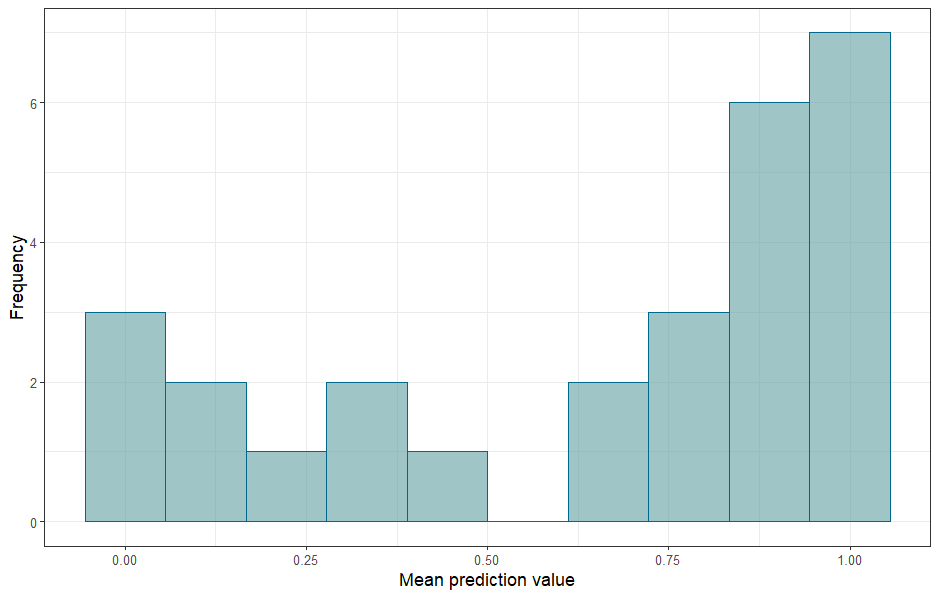

*Figure A13: Prediction values from a Bayesian model of pair status, for the widowed pairs in the analysis. Predictions (1=faithful, 0=divorced) were averaged across 100 runs, generating the mean prediction value.*

Figure A14 shows the standard deviations of the predictions generated by the model, which shows a relatively uniform distribution. A t-test of the groups has a *p*-value of 0.649 for the alternative hypothesis that mean prediction standard deviation is different between the prediction classes, suggesting that the confidence around individual predictions is similar between the prediction classes.

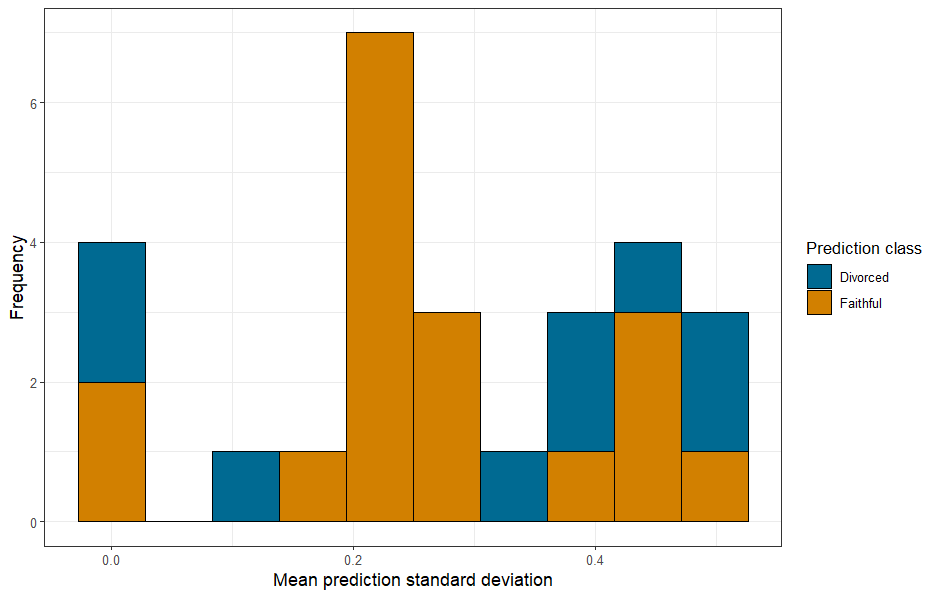

*Figure A14: Standard deviations from a Bayesian model of pair status, for the widowed pairs in the analysis, categorised by their predicted pair status. Predictions (1=faithful, 0=divorced) were generated across 100 runs, from which the standard deviation was calculated.*

The presence of relatively high standard deviation values suggests that, were the process to be run again, some pairs (particularly those close to the 0.6 cutoff) might be assigned a different status. With a relatively small sample size, these differences could lead to changes in model results, and therefore conclusions. In order to test whether the specific set of predictions used in our analysis had an outsized effect on our conclusions, we planned to conduct a sensitivity test, involving repredicting pair statuses. However, after repredicting pair statuses 500 times, only two individuals of 27 ever changed prediction (Table A10), with both being heavily biased towards their original pair status.

*Table A10: Original and rerun status predictions for the three individuals whose predicted status changed across 100 predictions from a Bayesian model of pair status.*

| **Pair number** | **Original model status** | **Predicted ‘faithful’ (/500)** | **Predicted ‘divorced’ (/500)** |
| --- | --- | --- | --- |
| 16 | Faithful | 481 | 19 |
| 17 | Divorced | 3 | 497 |

Models were rerun with all possible combinations of these changed pair statuses (three permutations in total). All models retained the same conclusions, although effect estimates for weighted degree in the weekend before/after and top opposite-sex SRI in the weekend before/after were almost significant in all three non-original permutations. As a significant result for these models would only support the conclusions in the main body of the text, this result is acceptable within the boundaries of the sensitivity analysis, and the reported result was retained as the conservative non-significant estimate.

An extension of this sensitivity analysis would be to perform it in a Bayesian framework which accounted for the uncertainty around the estimates.

As a further test of sensitivity to model results, we performed a permutation null test, rerunning the models 500 times with pair statuses randomly assigned between the pairs. This was in order to investigate the likelihood of our results being obtained by chance alone, or by the sample size effect of having only 9 divorced widowed and 18 faithful widowed pairs. As can be seen in Figure A15, the significant association score results presented in this analysis (topsri, degree.wknd, oppsex.wknd) were unlikely to be due only to sample size.

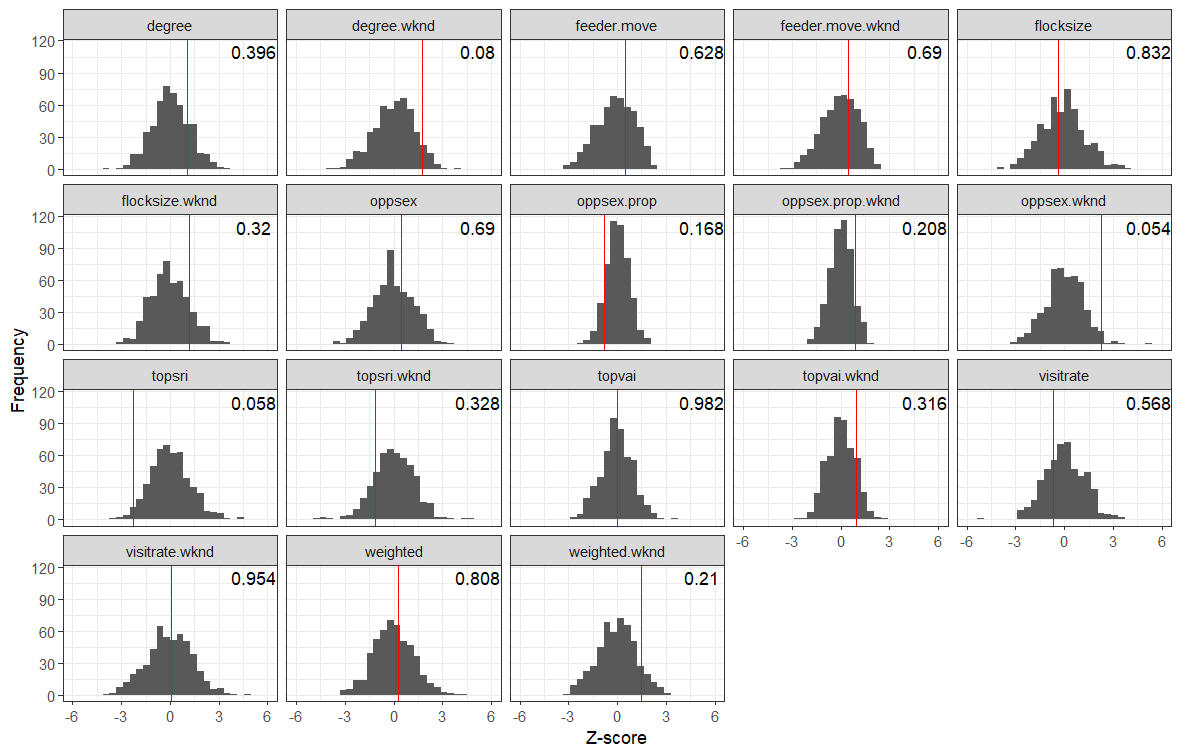

*Figure A15: Permutation test of analysis models, generated by rerunning the models 500 times with randomly assigned faithful/divorced status, while retaining the pair status sample sizes. Red lines represent the Z-score of the ‘Faithful (widowed):Pre-widowing’ coefficient in the original model, as presented in the main text. Text in the right hand side is the two-sided p-value of that original Z-score given the distribution of permuted results. The x axis has been constrained to -6, 6 for visualisation, which has removed seven extreme values, which were included in p-value calculation.*

#### Full models

Daily degree model - period before/after

*Table A11: Fixed effect coefficient estimates from a lognormal GLMM of the effect of pair status and widowing on daily degree. Significant coefficients are indicated in italics.*

| Coefficient | Estimate | Standard error | *p*-value |
| --- | --- | --- | --- |
| *Intercept* | *-3.85* | *0.05* | *<0.001* |
| Pair status Divorced | 0.024 | 0.074 | 0.748 |
| Pair status Divorced (widowed) | 0.105 | 0.171 | 0.539 |
| Pair status Faithful (widowed) | 0.138 | 0.119 | 0.246 |
| Post-widowing | 0.062 | 0.038 | 0.104 |
| *Pair status Divorced:Post-widowing* | *0.155* | *0.074* | *0.036* |
| Pair status Divorced (widowed):Post-widowing | 0.247 | 0.175 | 0.158 |
| Pair status Faithful (widowed):Post-widowing | 0.089 | 0.097 | 0.360 |

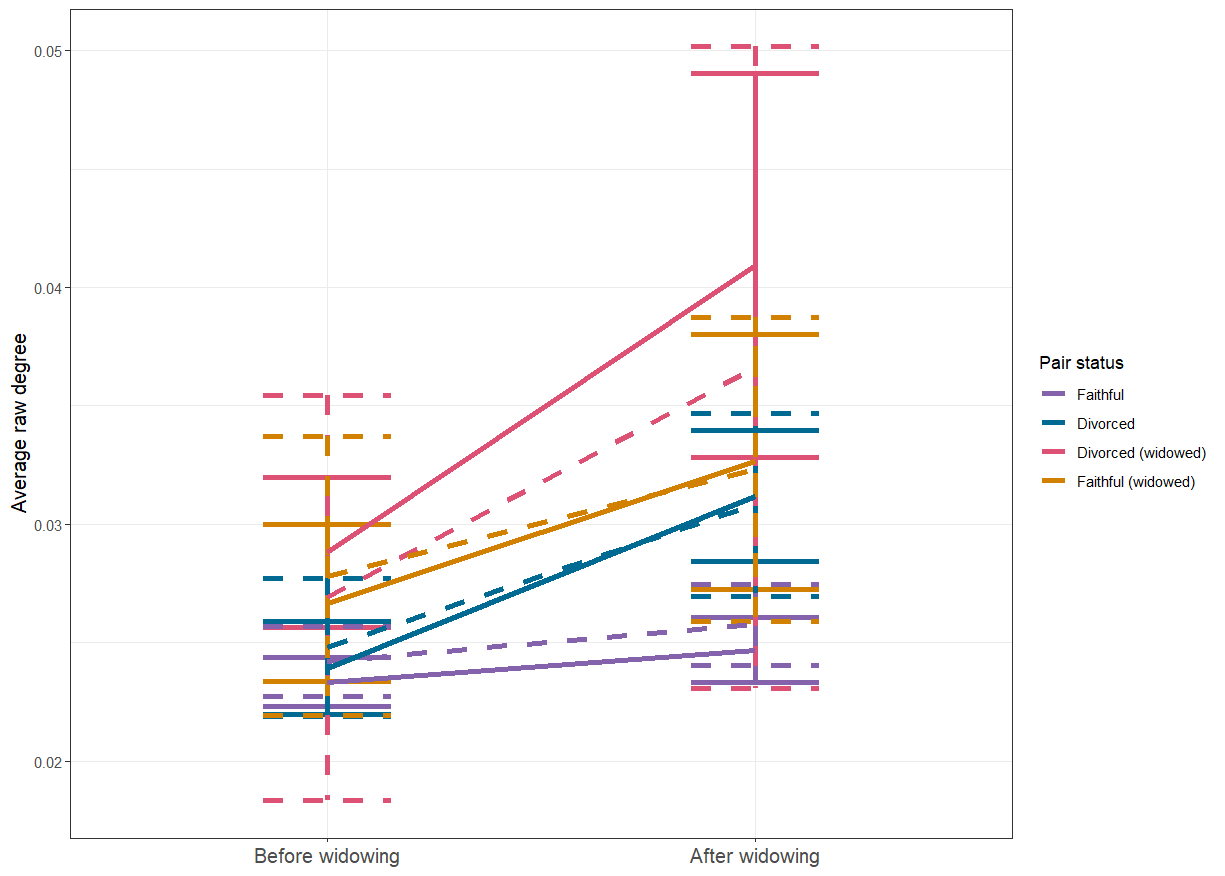

*Figure A16: Predicted and observed difference between average daily degree before and after the loss of a bird’s mate (widowing). Solid points and lines are averages and standard error from the raw data, while dashed error bars and lines are predictions from a log normal generalised linear mixed model, with upper and lower bars representing a 95% confidence interval. Predictions were made using ggeffects, accounting for random effects and a zero-inflation term (Lüdecke, 2018).*

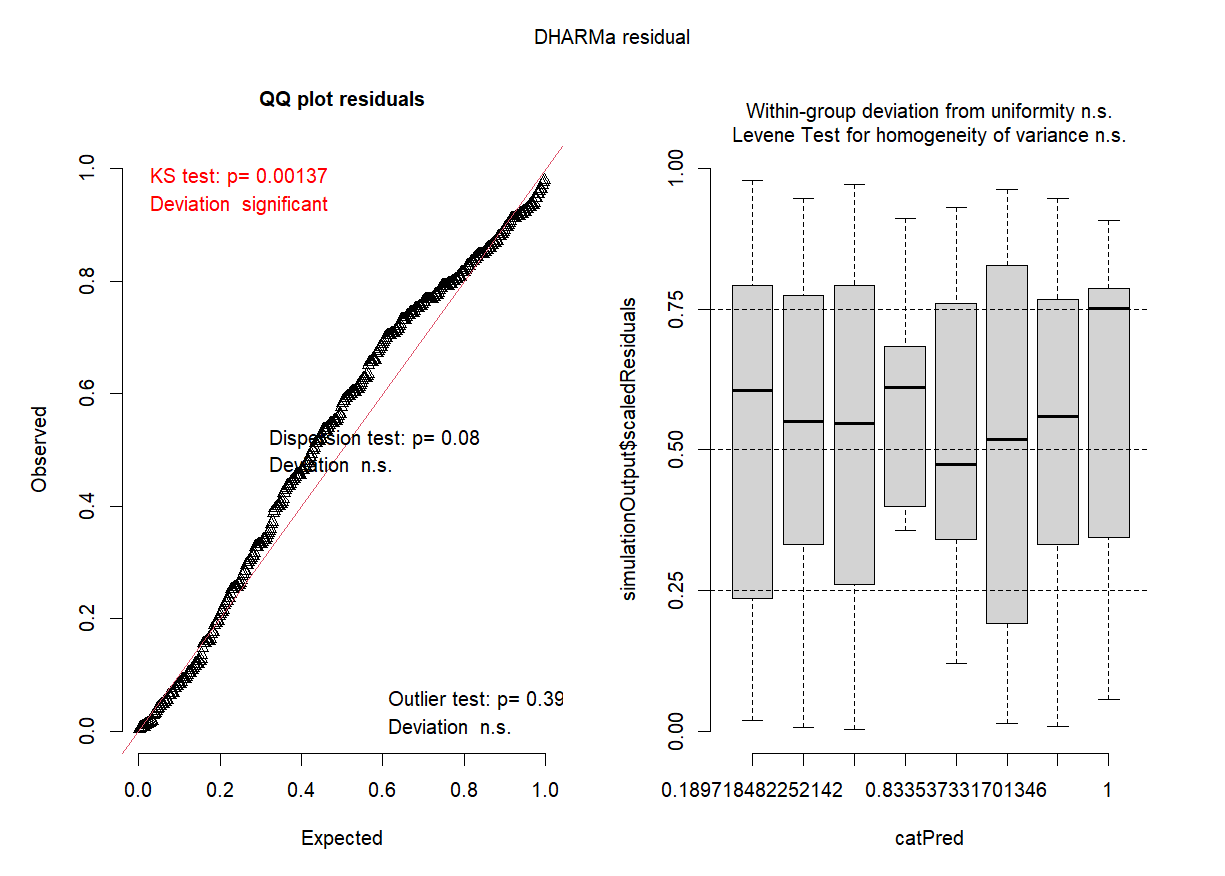

*Figure A17: Residuals from a lognormal GLMM of the effect of pair status and widowing on daily degree, as calculated by DHARMa (Hartig, 2018).*

Daily degree model - weekend before/after

*Table A12: Fixed effect coefficient estimates from a lognormal GLMM of the effect of pair status and widowing on daily degree in the weekend before and after widowing. Significant coefficients are indicated in italics.*

| Coefficient | Estimate | Standard error | *p*-value |
| --- | --- | --- | --- |
| *Intercept* | *-3.576* | *0.051* | *<0.001* |
| Pair status Divorced | 0.164 | 0.085 | 0.054 |
| *Pair status Divorced (widowed)* | *0.382* | *0.175* | *0.029* |
| Pair status Faithful (widowed) | 0.184 | 0.131 | 0.161 |
| Post-widowing | -0.084 | 0.053 | 0.114 |
| Pair status Divorced:Post-widowing | -0.091 | 0.117 | 0.434 |
| Pair status Divorced (widowed):Post-widowing | -0.102 | 0.198 | 0.606 |
| Pair status Faithful (widowed):Post-widowing | 0.231 | 0.132 | 0.081 |

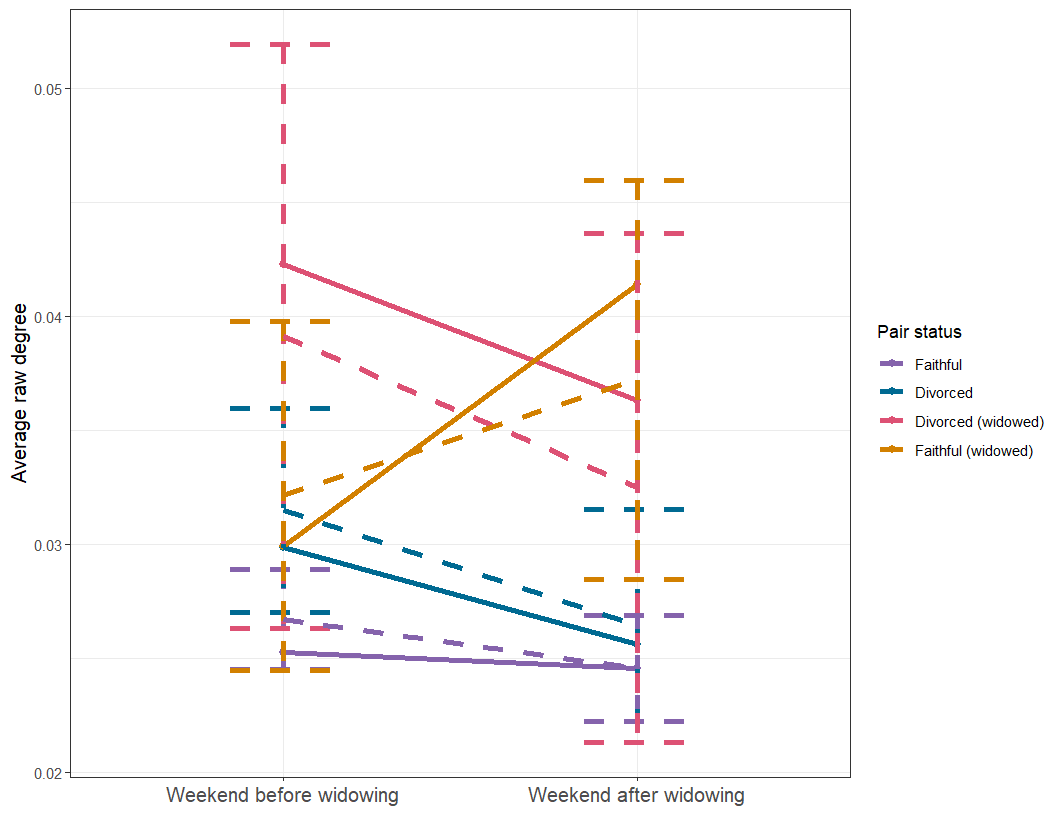

*Figure A18: Predicted and observed difference between average daily degree in the weekend before and after the loss of a bird’s mate (widowing). Solid points and lines are averages and standard error from the raw data, while dashed error bars and lines are predictions from a log normal generalised linear mixed model, with upper and lower bars representing a 95% confidence interval. Predictions were made using ggeffects, accounting for random effects and a zero-inflation term (Lüdecke, 2018).*

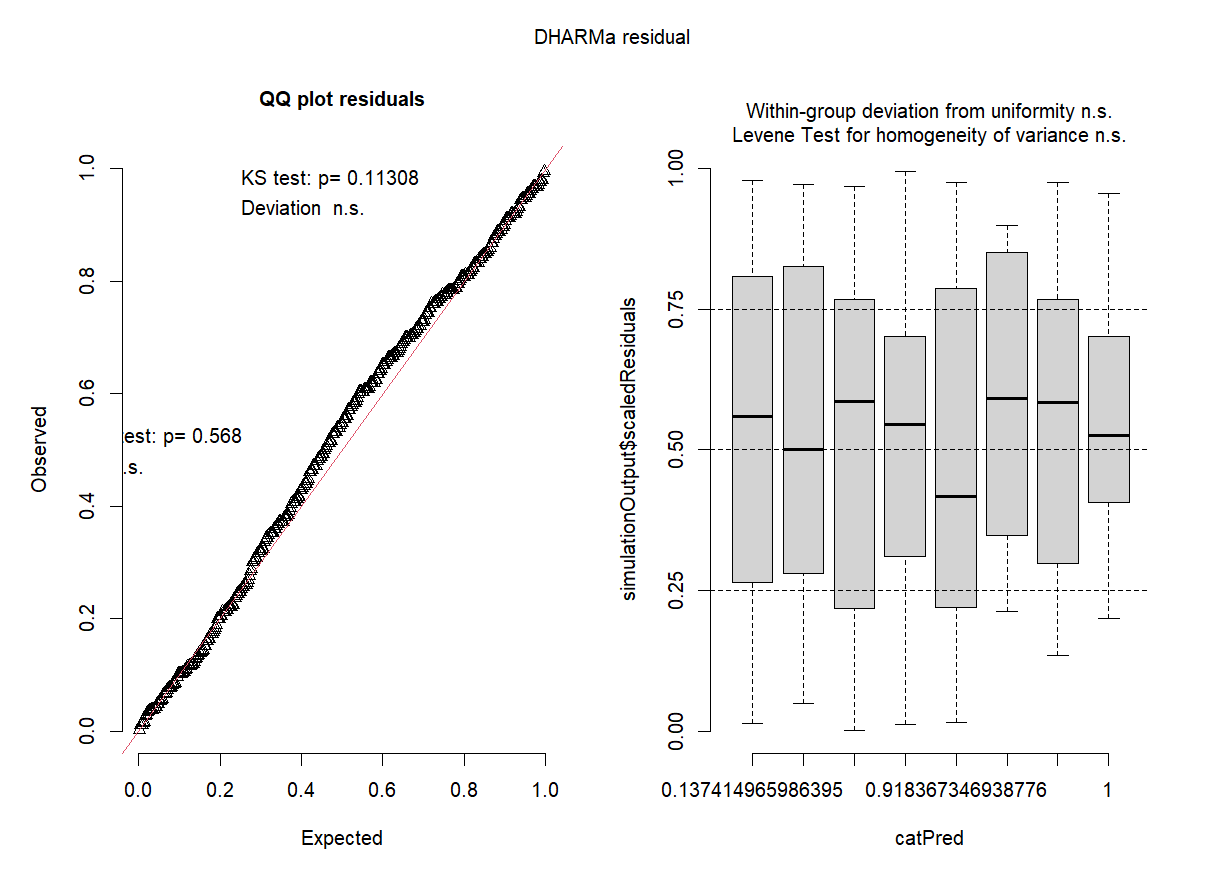

*Figure A19: Residuals from a lognormal GLMM of the effect of pair status and widowing on daily degree in the weekend before and after widowing, as calculated by DHARMa (Hartig, 2018).*

Weighted model - period before/after

*Table A13: Fixed effect coefficient estimates from a lognormal GLMM of the effect of pair status and widowing on weighted degree. Significant coefficients are indicated in italics.*

| Coefficient | Estimate | Standard error | *p*-value |
| --- | --- | --- | --- |
| *Intercept* | *2.086* | *0.04* | *<0.001* |
| Pair status Divorced | 0.025 | 0.074 | 0.734 |
| Pair status Divorced (widowed) | 0.077 | 0.190 | 0.686 |
| Pair status Faithful (widowed) | 0.066 | 0.108 | 0.538 |
| *Post-widowing* | *0.099* | *0.036* | *0.006* |
| Pair status Divorced:Post-widowing | 0.048 | 0.068 | 0.486 |
| Pair status Divorced (widowed):Post-widowing | 0.164 | 0.146 | 0.261 |
| Pair status Faithful (widowed):Post-widowing | 0.028 | 0.095 | 0.769 |

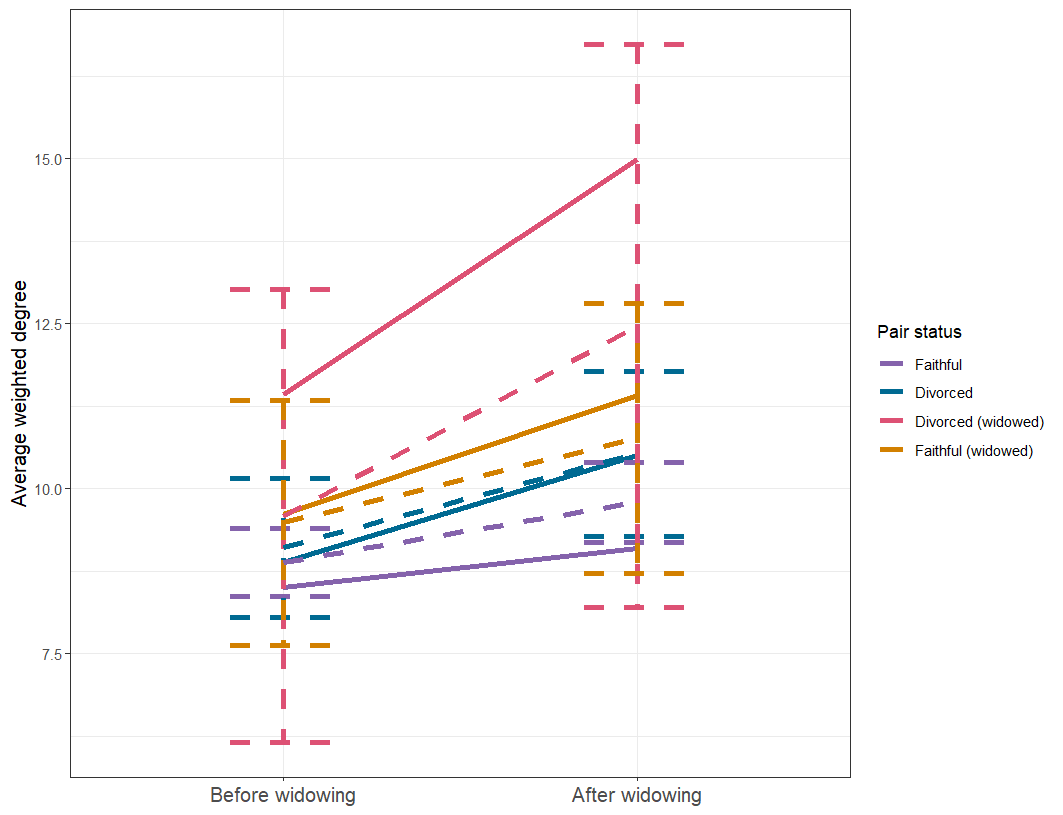

*Figure A20:* *Predicted and observed difference between average weighted degree before and after the loss of a bird’s mate (widowing). Solid points and lines are averages and standard error from the raw data, while dashed error bars and lines are predictions from a log normal generalised linear mixed model, with upper and lower bars representing a 95% confidence interval. Predictions were made using ggeffects, accounting a zero-inflation term (Lüdecke, 2018). Predictions were made without random effects for visualisation purposes.*

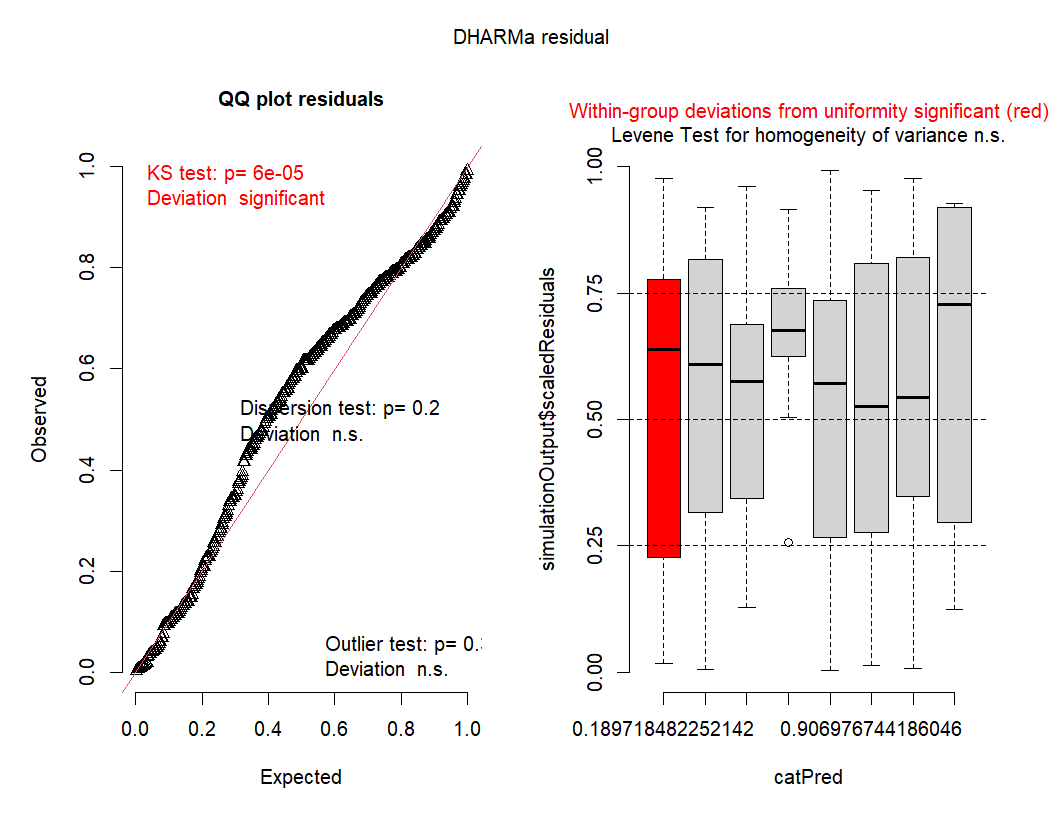

*Figure A21: Residuals from a lognormal GLMM of the effect of pair status and widowing on weighted degree, as calculated by DHARMa (Hartig, 2018).*

Weighted model - weekend before/after

*Table A14: Fixed effect coefficient estimates from a lognormal GLMM of the effect of pair status and widowing on weighted degree in the weekend before and after widowing. Significant coefficients are indicated in italics.*

| Coefficient | Estimate | Standard error | *p*-value |
| --- | --- | --- | --- |
| *Intercept* | *2.396* | *0.043* | *<0.001* |
| *Pair status Divorced* | *0.159* | *0.076* | *0.036* |
| *Pair status Divorced (widowed)* | *0.325* | *0.156* | *0.037* |
| Pair status Faithful (widowed) | 0.129 | 0.112 | 0.250 |
| *Pre-widowing* | *-0.101* | *0.049* | *0.038* |
| Pair status Divorced:Pre-widowing | -0.103 | 0.100 | 0.304 |
| Pair status Divorced (widowed):Pre-widowing | -0.041 | 0.182 | 0.822 |
| Pair status Faithful (widowed):Pre-widowing | 0.183 | 0.124 | 0.142 |

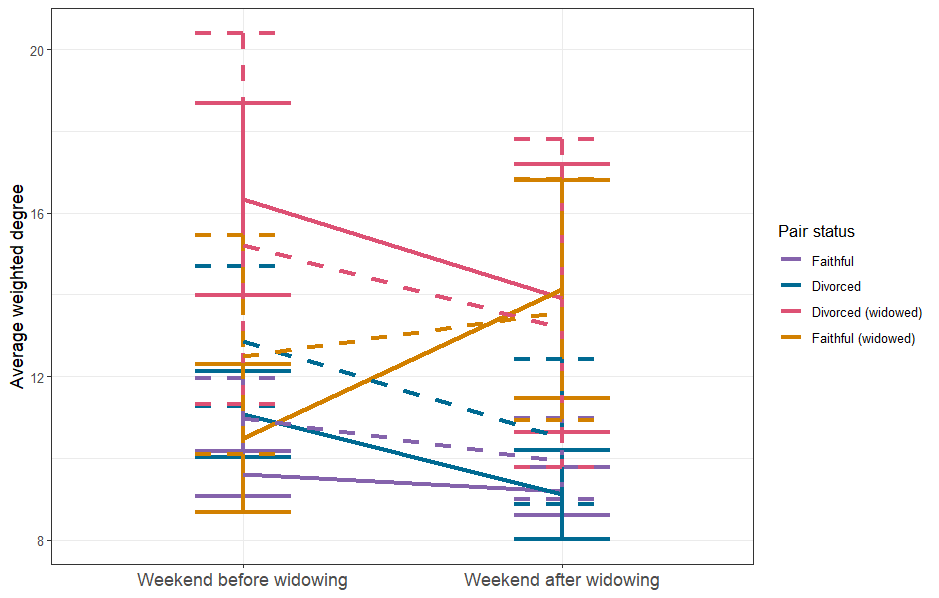

*Figure A22: Predicted and observed difference between average weighted degree in the weekend before and after the loss of a bird’s mate (widowing). Solid points and lines are averages and standard error from the raw data, while dashed error bars and lines are predictions from a Gaussian generalised linear mixed model, with upper and lower bars representing a 95% confidence interval. Predictions were made using ggeffects, accounting for random effects and a zero-inflation term (Lüdecke, 2018).*

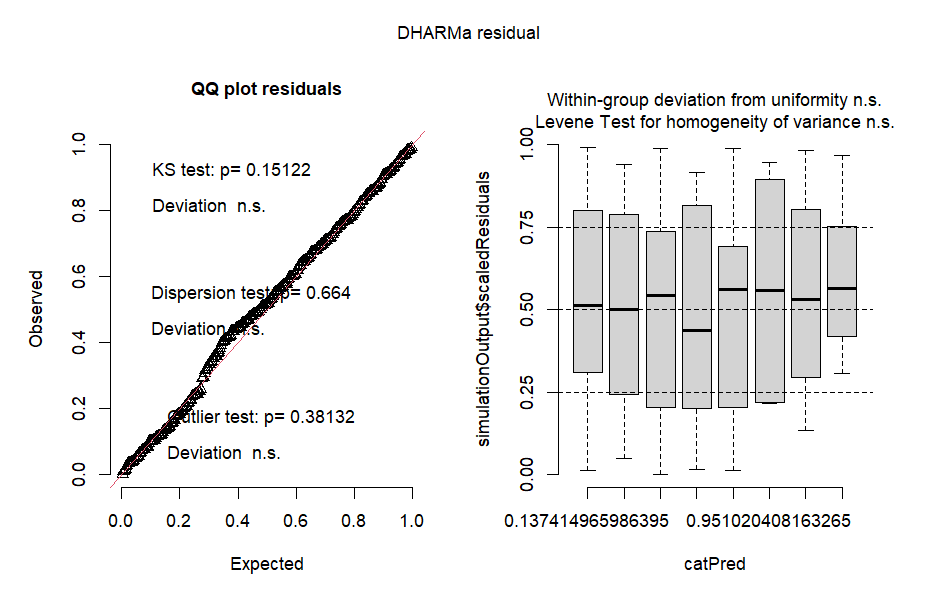

*Figure A23: Residuals from a Gaussian GLMM of the effect of pair status and widowing on weighted degree in the weekend before and after widowing, as calculated by DHARMa (Hartig, 2018).*

Opposite sex degree - period before/after

*Table A15: Fixed effect coefficient estimates from a lognormal GLMM of the effect of pair status and widowing on opposite sex degree. Significant coefficients are indicated in italics.*

| Coefficient | Estimate | Standard error | *p*-value |
| --- | --- | --- | --- |
| Intercept | -3.510 | 0.042 | <0.001 |
| Pair status Divorced | -0.012 | 0.065 | 0.849 |
| *Pair status Divorced (widowed)* | *0.323* | *0.123* | *0.009* |
| Pair status Faithful (widowed) | 0.065 | 0.097 | 0.501 |
| Post-widowing | -0.015 | 0.032 | 0.634 |
| *Pair status Divorced:Post-widowing* | *0.188* | *0.059* | *0.002* |
| Pair status Divorced (widowed):Post-widowing | 0.086 | 0.106 | 0.418 |
| Pair status Faithful (widowed):Post-widowing | 0.037 | 0.082 | 0.650 |

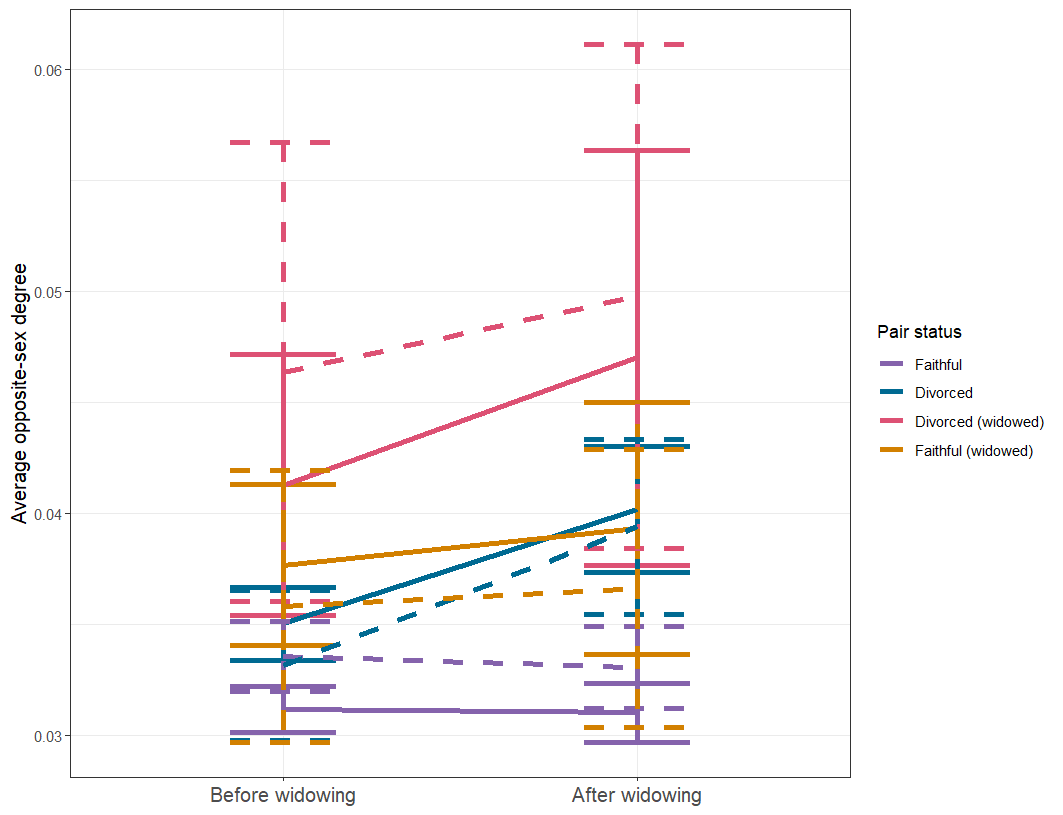

*Figure A24: Predicted and observed difference between average opposite sex degree before and after the loss of a bird’s mate (widowing). Solid points and lines are averages and standard error from the raw data, while dashed error bars and lines are predictions from a log normal generalised linear mixed model, with upper and lower bars representing a 95% confidence interval. Predictions were made using ggeffects, accounting for random effects and a zero-inflation term (Lüdecke, 2018).*

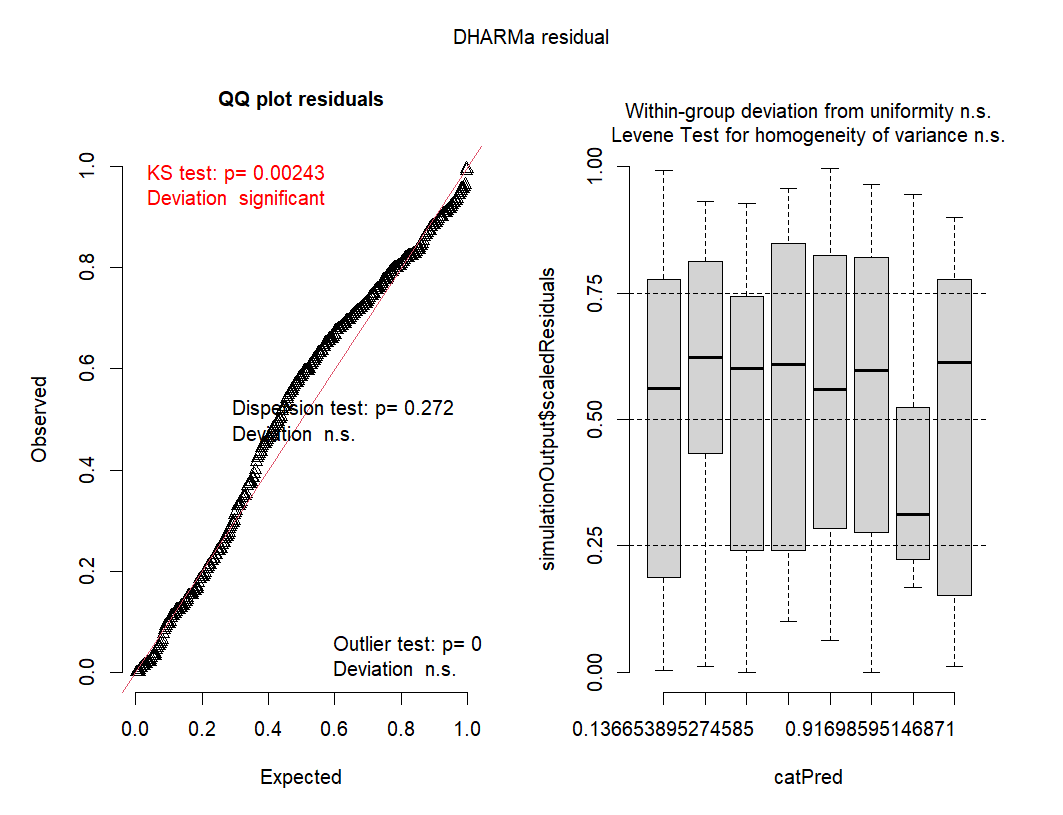

*Figure A25: Residuals from a lognormal GLMM of the effect of pair status and widowing on opposite sex degree, as calculated by DHARMa (Hartig, 2018).*

Opposite sex model - weekend before/after

*Table A16: Fixed effect coefficient estimates from a lognormal GLMM of the effect of pair status and widowing on opposite sex degree in the weekend before and after widowing. Significant coefficients are indicated in italics.*

| Coefficient | Estimate | Standard error | *p*-value |
| --- | --- | --- | --- |
| *Intercept* | *-3.462* | *0.049* | *<0.001* |
| Pair status Divorced | 0.074 | 0.085 | 0.381 |
| *Pair status Divorced (widowed)* | *0.367* | *0.160* | *0.022* |
| Pair status Faithful (widowed) | 0.066 | 0.128 | 0.610 |
| *Post-widowing* | *-0.096* | *0.049* | *0.049* |
| Pair status Divorced:Post-widowing | 0.093 | 0.100 | 0.352 |
| Pair status Divorced (widowed):Post-widowing | -0.174 | 0.182 | 0.339 |
| *Pair status Faithful (widowed):Post-widowing* | *0.285* | *0.125* | *0.023* |

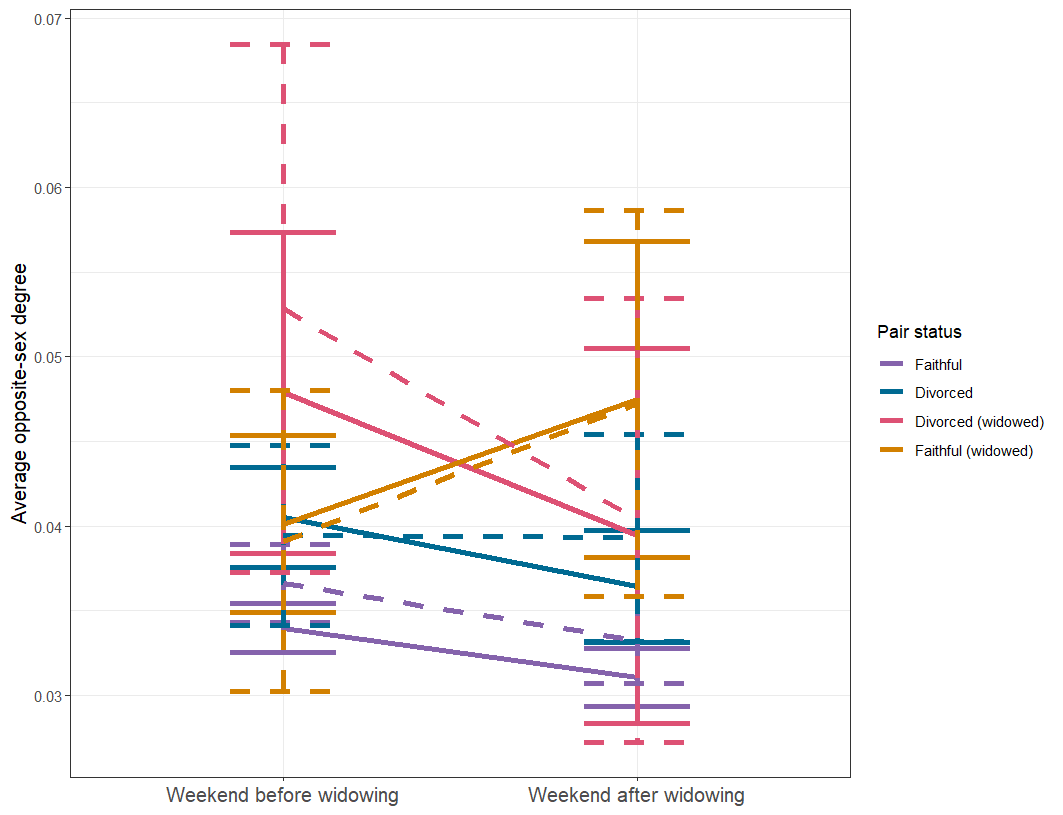

*Figure A26: Predicted and observed difference between average opposite sex degree in the weekend before and after the loss of a bird’s mate (widowing). Solid points and lines are averages and standard error from the raw data, while dashed error bars and lines are predictions from a log normal generalised linear mixed model, with upper and lower bars representing a 95% confidence interval. Predictions were made using ggeffects, accounting for random effects and a zero-inflation term (Lüdecke, 2018).*

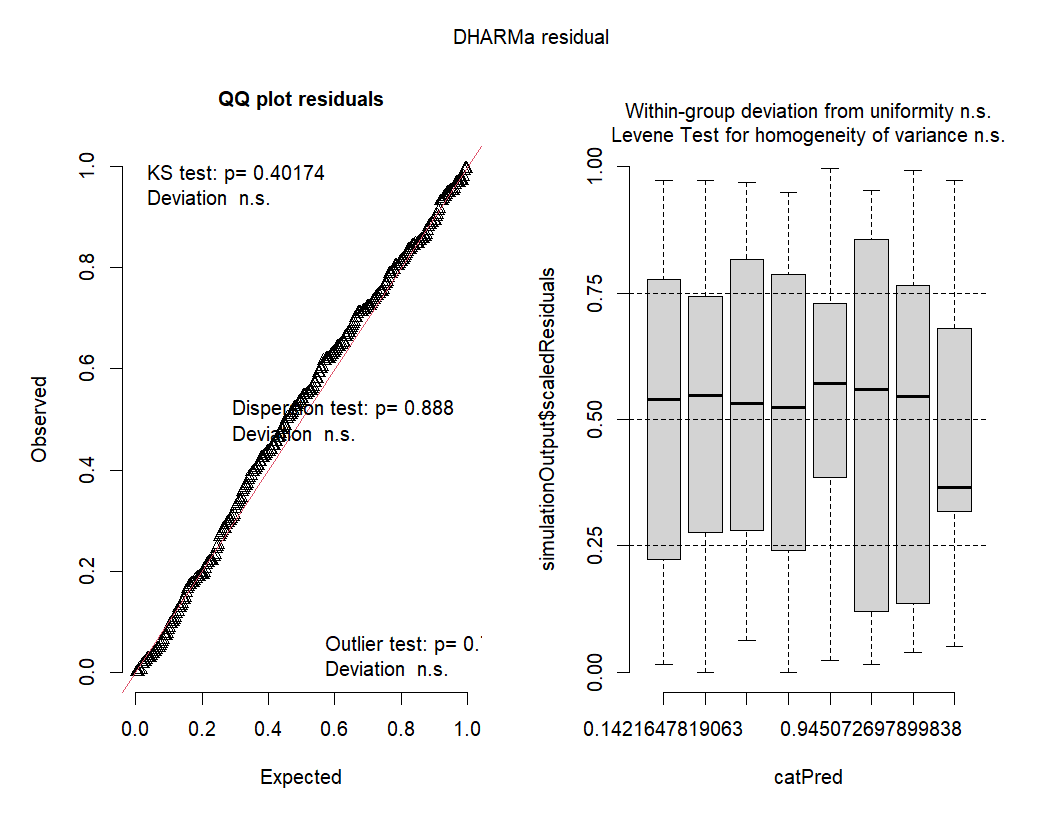

*Figure A27: Residuals from a lognormal GLMM of the effect of pair status and widowing on weighted degree in the weekend before and after widowing, as calculated by DHARMa (Hartig, 2018).*

Opposite sex proportion - period before/after

*Table A17: Fixed effect coefficient estimates from a Gaussian GLMM of the effect of pair status and widowing on opposite sex proportion. Significant coefficients are indicated in italics.*

| Coefficient | Estimate | Standard error | *p*-value |
| --- | --- | --- | --- |
| *Intercept* | *0.376* | *0.008* | *<0.001* |
| Pair status Divorced | -0.014 | 0.012 | 0.215 |
| Pair status Divorced (widowed) | 0.007 | 0.024 | 0.777 |
| Pair status Faithful (widowed) | 0.003 | 0.016 | 0.855 |
| Post-widowing | 0.011 | 0.006 | 0.049 |
| Pair status Divorced:Post-widowing | -0.005 | 0.011 | 0.642 |
| Pair status Divorced (widowed):Post-widowing | -0.002 | 0.024 | 0.948 |
| Pair status Faithful (widowed):Post-widowing | -0.014 | 0.017 | 0.412 |

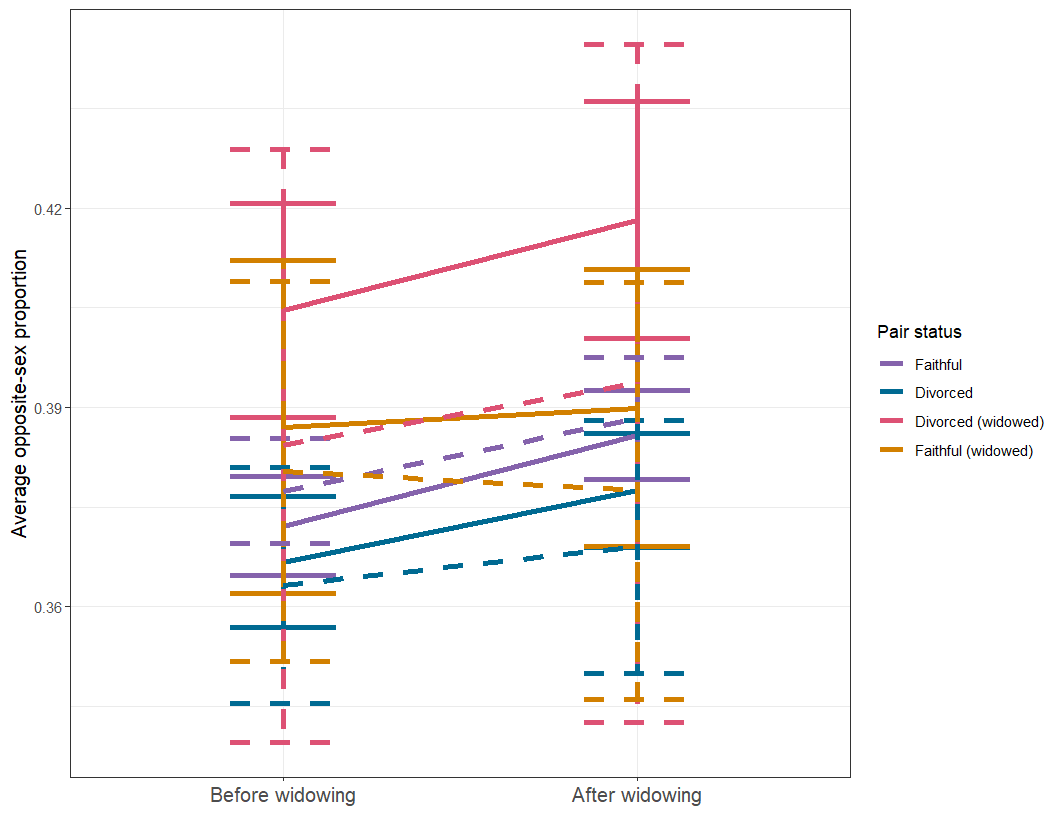

*Figure A28: Predicted and observed difference between average opposite sex proportion before and after the loss of a bird’s mate (widowing). Solid points and lines are averages and standard error from the raw data, while dashed error bars and lines are predictions from a Gaussian generalised linear mixed model, with upper and lower bars representing a 95% confidence interval. Predictions were made using ggeffects, accounting for random effects and a zero-inflation term (Lüdecke, 2018).*

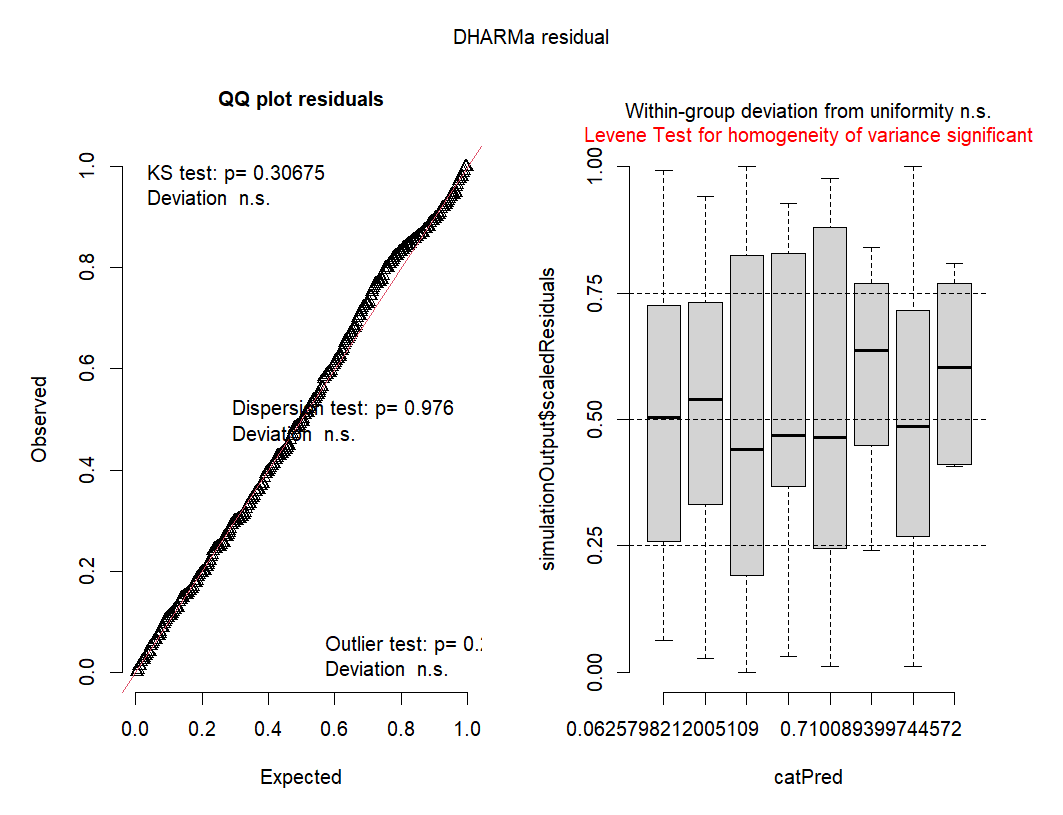

*Figure A29: Residuals from a Gaussian GLMM of the effect of pair status and widowing on opposite sex proportion, as calculated by DHARMa (Hartig, 2018).*

Opposite sex proportion - weekend before/after

*Table A18: Fixed effect coefficient estimates from a Gaussian GLMM of the effect of pair status and widowing on opposite sex proportion in the weekend before and after widowing. Significant coefficients are indicated in italics.*

| **Coefficient** | **Estimate** | **Standard error** | ***p*-value** |
| --- | --- | --- | --- |
| *Intercept* | *0.378* | *0.012* | *<0.001* |
| Pair status Divorced | 0.001 | 0.014 | 0.934 |
| Pair status Divorced (widowed) | 0.026 | 0.029 | 0.360 |
| Pair status Faithful (widowed) | -0.014 | 0.019 | 0.470 |
| Post-widowing | -0.005 | 0.007 | 0.503 |
| Pair status Divorced:Post-widowing | 0.003 | 0.014 | 0.856 |
| Pair status Divorced (widowed):Post-widowing | -0.035 | 0.028 | 0.210 |
| Pair status Faithful (widowed):Post-widowing | 0.020 | 0.022 | 0.373 |

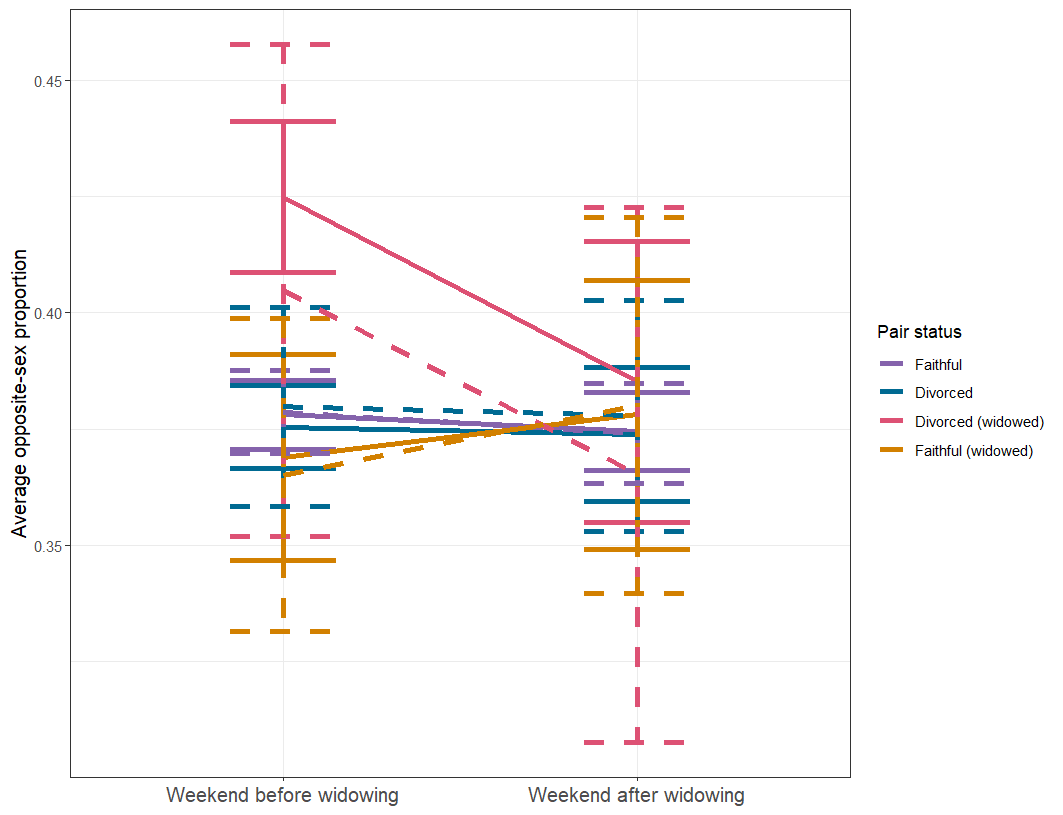

*Figure A30: Predicted and observed difference between average opposite sex proportion in the weekend before and after the loss of a bird’s mate (widowing). Solid points and lines are averages and standard error from the raw data, while dashed error bars and lines are predictions from a Gaussian generalised linear mixed model, with upper and lower bars representing a 95% confidence interval. Predictions were made using ggeffects, accounting for random effects and a zero-inflation term (Lüdecke, 2018).*

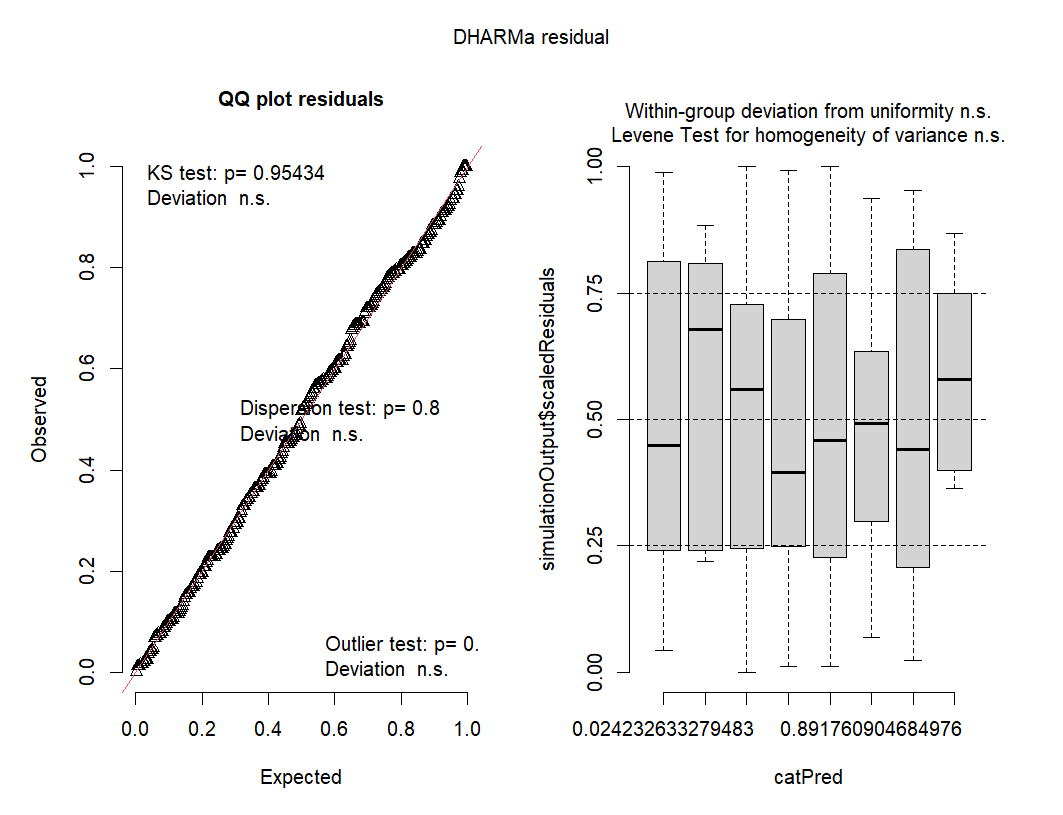

*Figure A31: Residuals from a Gaussian GLMM of the effect of pair status and widowing on opposite sex proportion in the weekend before and after widowing, as calculated by DHARMa (Hartig, 2018).*

Flock size - period before/after

*Table A19: Fixed effect coefficient estimates from a lognormal GLMM of the effect of pair status and widowing on flock size. Significant coefficients are indicated in italics.*

| Coefficient | Estimate | Standard error | *p*-value |
| --- | --- | --- | --- |
| *Intercept* | *11.970* | *0.852* | *<0.001* |
| Pair status Divorced | 0.526 | 0.602 | 0.382 |
| *Pair status Divorced (widowed)* | *3.282* | *1.240* | *0.008* |
| Pair status Faithful (widowed) | 1.072 | 0.836 | 0.200 |
| Post-widowing | -0.302 | 0.295 | 0.305 |
| Pair status Divorced:Post-widowing | 0.913 | 0.579 | 0.115 |
| Pair status Divorced (widowed):Post-widowing | 0.226 | 1.272 | 0.859 |
| Pair status Faithful (widowed):Post-widowing | -0.354 | 0.896 | 0.693 |

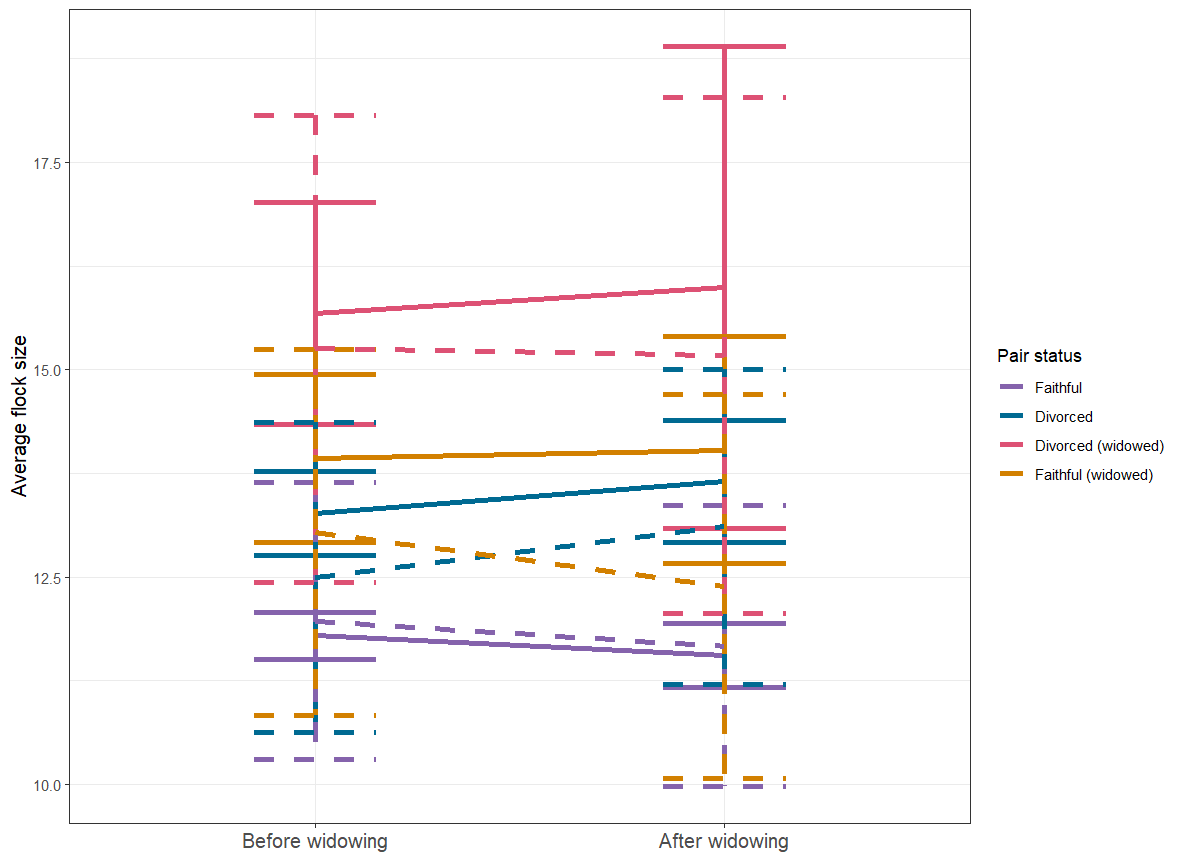

*Figure A32: Predicted and observed difference between average flock size before and after the loss of a bird’s mate (widowing). Solid points and lines are averages and standard error from the raw data, while dashed error bars and lines are predictions from a lognormal generalised linear mixed model, with upper and lower bars representing a 95% confidence interval. Predictions were made using ggeffects, accounting for random effects and a zero-inflation term (Lüdecke, 2018).*

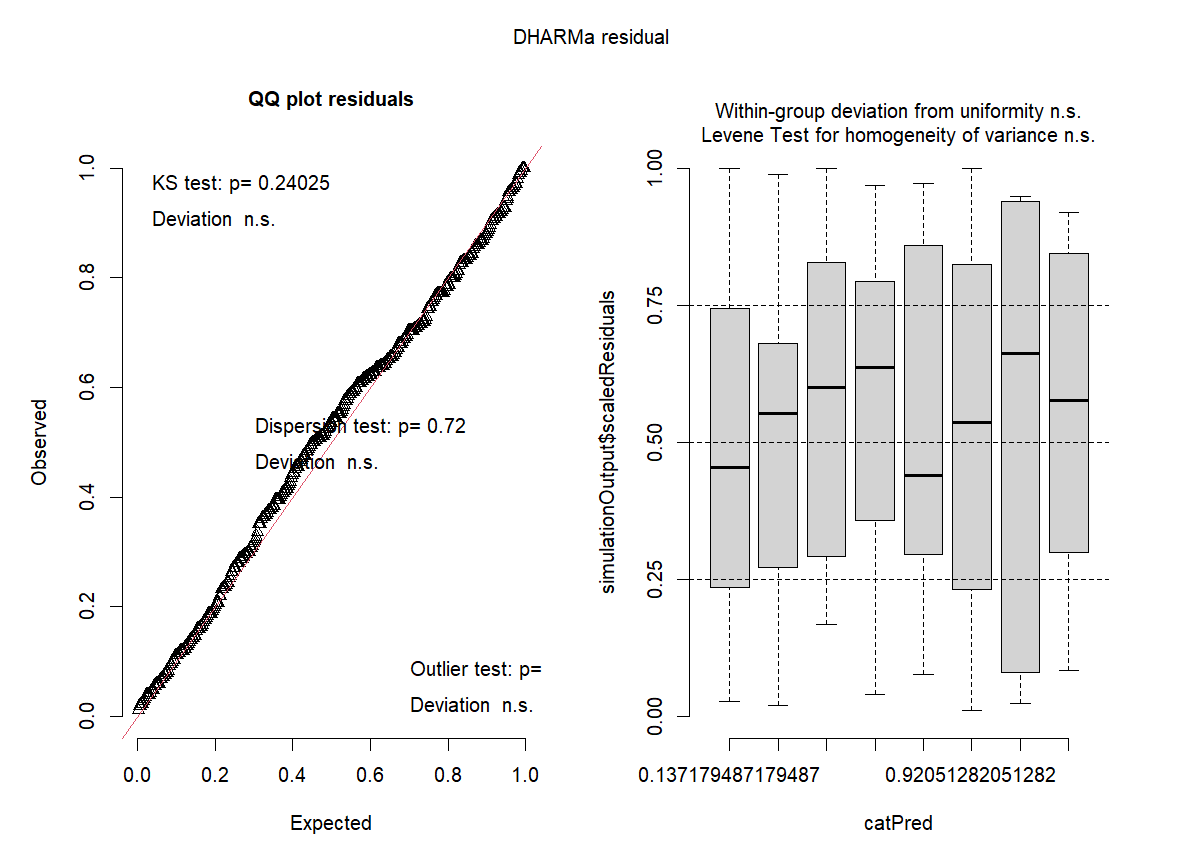

*Figure A33: Residuals from a lognormal GLMM of the effect of pair status and widowing on flock size, as calculated by DHARMa (Hartig, 2018).*

Flock size - weekend before/after

*Table A20: Fixed effect coefficient estimates from a lognormal GLMM of the effect of pair status and widowing on average flock size in the weekend before and after widowing. Significant coefficients are indicated in italics.*

| Coefficient | Estimate | Standard error | *p*-value |
| --- | --- | --- | --- |
| *Intercept* | *12.915* | *0.819* | *<0.001* |
| Pair status Divorced | 1.166 | 0.937 | 0.214 |
| *Pair status Divorced (widowed)* | *3.754* | *1.910* | *0.049* |
| Pair status Faithful (widowed) | 1.207 | 1.344 | 0.369 |
| *Post-widowing* | *-1.244* | *0.529* | *0.019* |
| Pair status Divorced:Post-widowing | 0.244 | 1.106 | 0.825 |
| Pair status Divorced (widowed):Post-widowing | -1.512 | 2.148 | 0.482 |
| Pair status Faithful (widowed):Post-widowing | 1.944 | 1.696 | 0.252 |

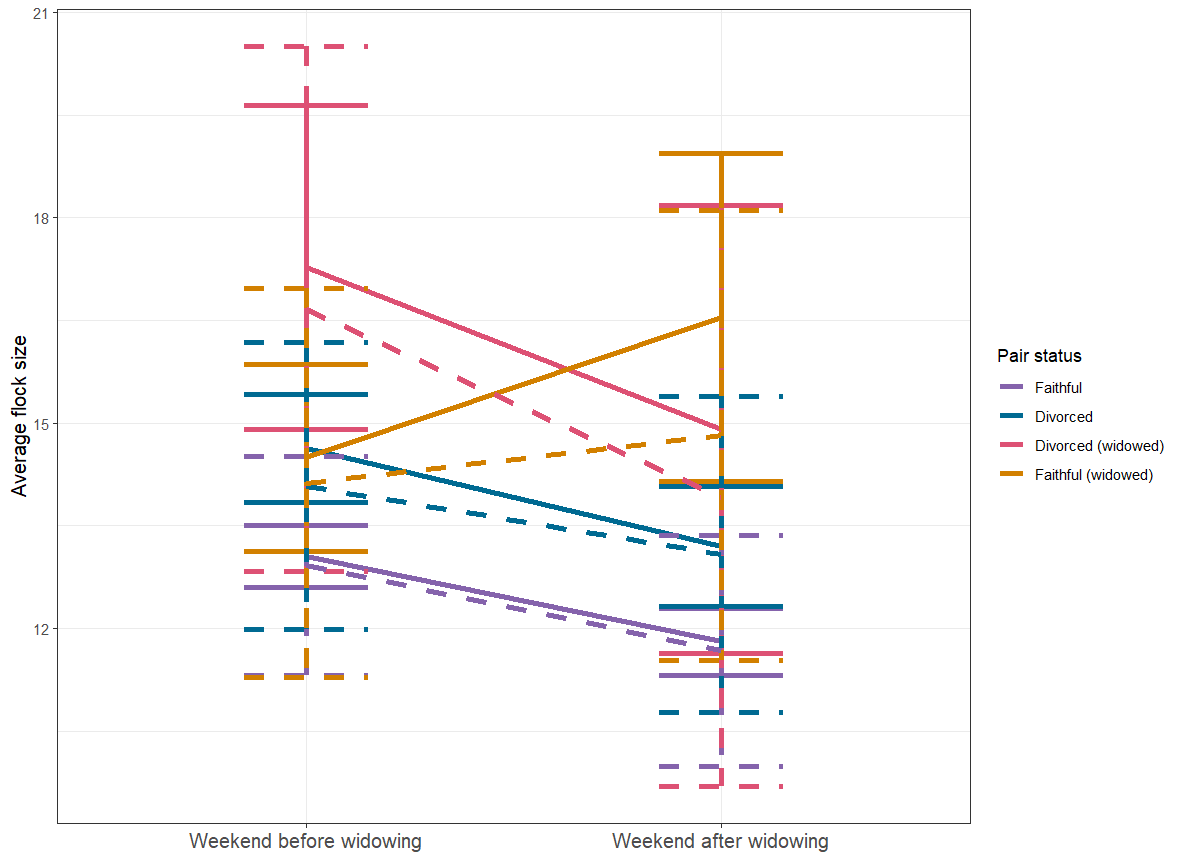

*Figure A34: Predicted and observed difference between average flock size in the weekend before and after the loss of a bird’s mate (widowing). Solid points and lines are averages and standard error from the raw data, while dashed error bars and lines are predictions from a lognormal generalised linear mixed model, with upper and lower bars representing a 95% confidence interval. Predictions were made using ggeffects, accounting for random effects and a zero-inflation term (Lüdecke, 2018).*

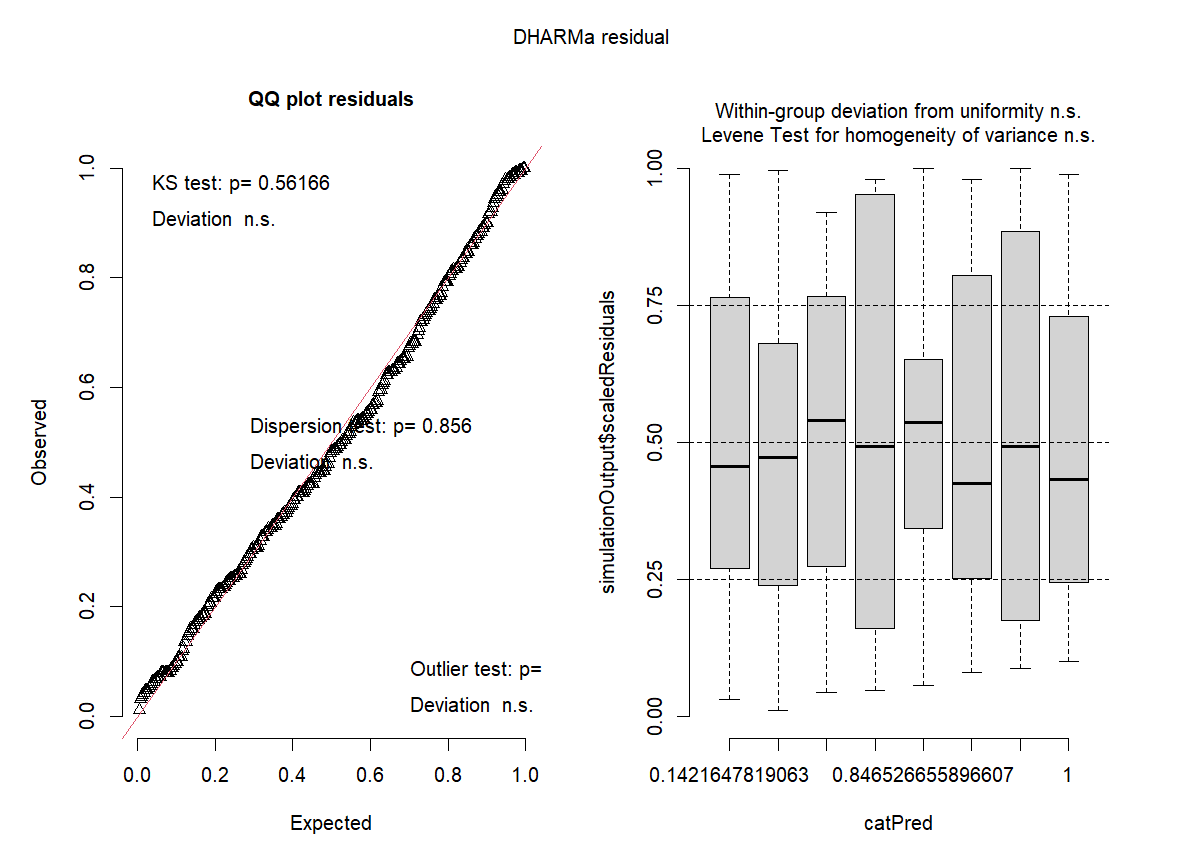

*Figure A35: Residuals from a lognormal GLMM of the effect of pair status and widowing on average flock size in the weekend before and after widowing, as calculated by DHARMa (Hartig, 2018).*

Feeder visitation rate - period before/after

*Table A21: Fixed effect coefficient estimates from a lognormal GLMM of the effect of pair status and widowing on feeder visitation rate. Significant coefficients are indicated in italics.*

| Coefficient | Estimate | Standard error | *p*-value |
| --- | --- | --- | --- |
| *Intercept* | *2.563* | *0.076* | *<0.001* |
| Pair status Divorced | 0.114 | 0.082 | 0.167 |
| Pair status Divorced (widowed) | 0.078 | 0.216 | 0.720 |
| Pair status Faithful (widowed) | 0.105 | 0.120 | 0.381 |
| *Post-widowing* | *0.163* | *0.037* | *<0.001* |
| Pair status Divorced:Post-widowing | 0.099 | 0.159 | 0.532 |
| Pair status Divorced (widowed):Post-widowing | 0.099 | 0.159 | 0.532 |
| Pair status Faithful (widowed):Post-widowing | -0.067 | 0.100 | 0.503 |

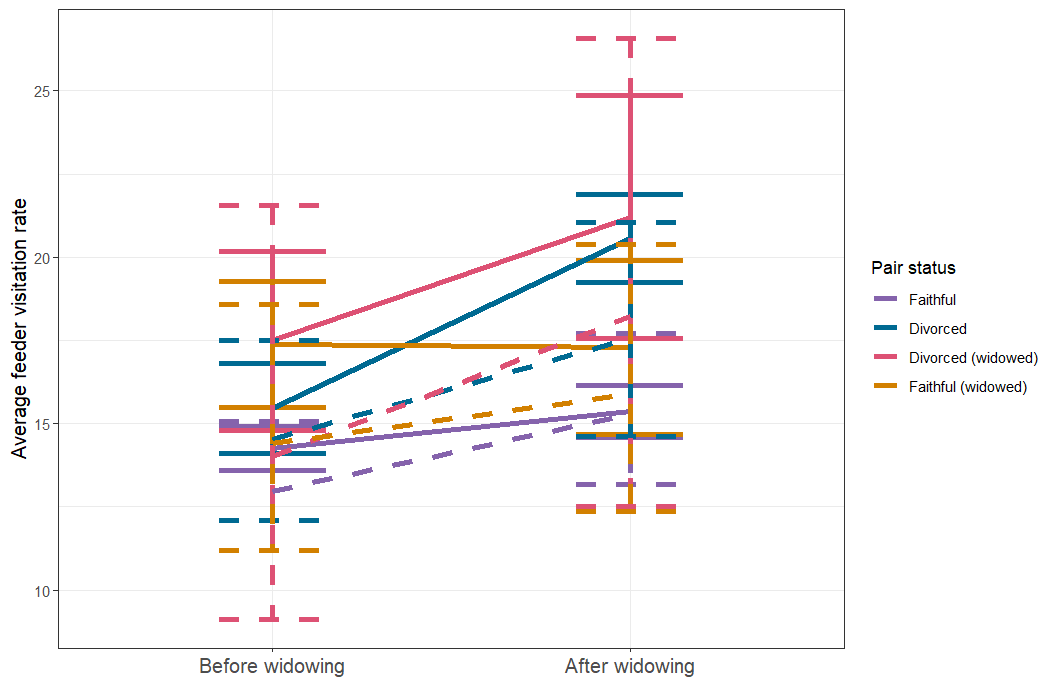

*Figure A36: Predicted and observed difference between average feeder visitation rate before and after the loss of a bird’s mate (widowing). Solid points and lines are averages and standard error from the raw data, while dashed error bars and lines are predictions from a lognormal generalised linear mixed model, with upper and lower bars representing a 95% confidence interval. Predictions were made using ggeffects, accounting for random effects and a zero-inflation term (Lüdecke, 2018).*

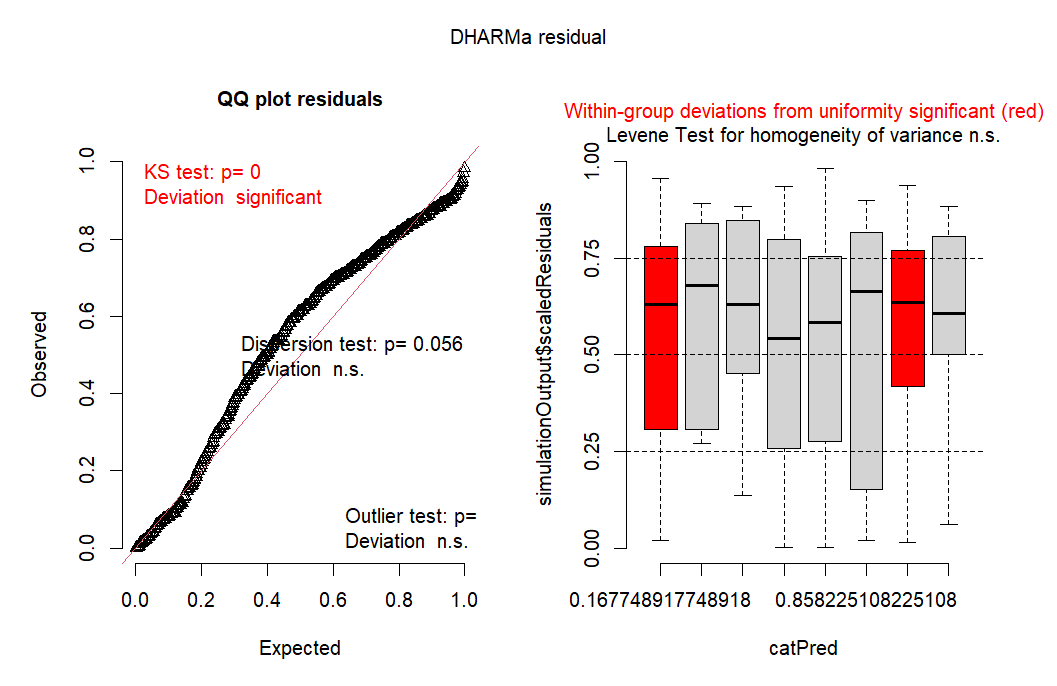

*Figure A37: Residuals from a lognormal GLMM of the effect of pair status and widowing on feeder visitation rate, as calculated by DHARMa (Hartig, 2018).*

Feeder visitation rate - weekend before/after

*Table A22: Fixed effect coefficient estimates from a lognormal GLMM of the effect of pair status and widowing on feeder visitation rate in the weekend before and after widowing. Significant coefficients are indicated in italics.*

| Coefficient | Estimate | Standard error | *p*-value |
| --- | --- | --- | --- |
| *Intercept* | *2.874* | *0.056* | *<0.001* |
| *Pair status Divorced* | *0.224* | *0.087* | *0.010* |
| Pair status Divorced (widowed) | 0.166 | 0.166 | 0.317 |
| Pair status Faithful (widowed) | 0.123 | 0.119 | 0.298 |
| *Post-widowing* | *-0.130* | *0.048* | *0.006* |
| Pair status Divorced:Post-widowing | -0.161 | 0.088 | 0.066 |
| Pair status Divorced (widowed):Post-widowing | 0.002 | 0.160 | 0.991 |
| Pair status Faithful (widowed):Post-widowing | 0.007 | 0.127 | 0.957 |

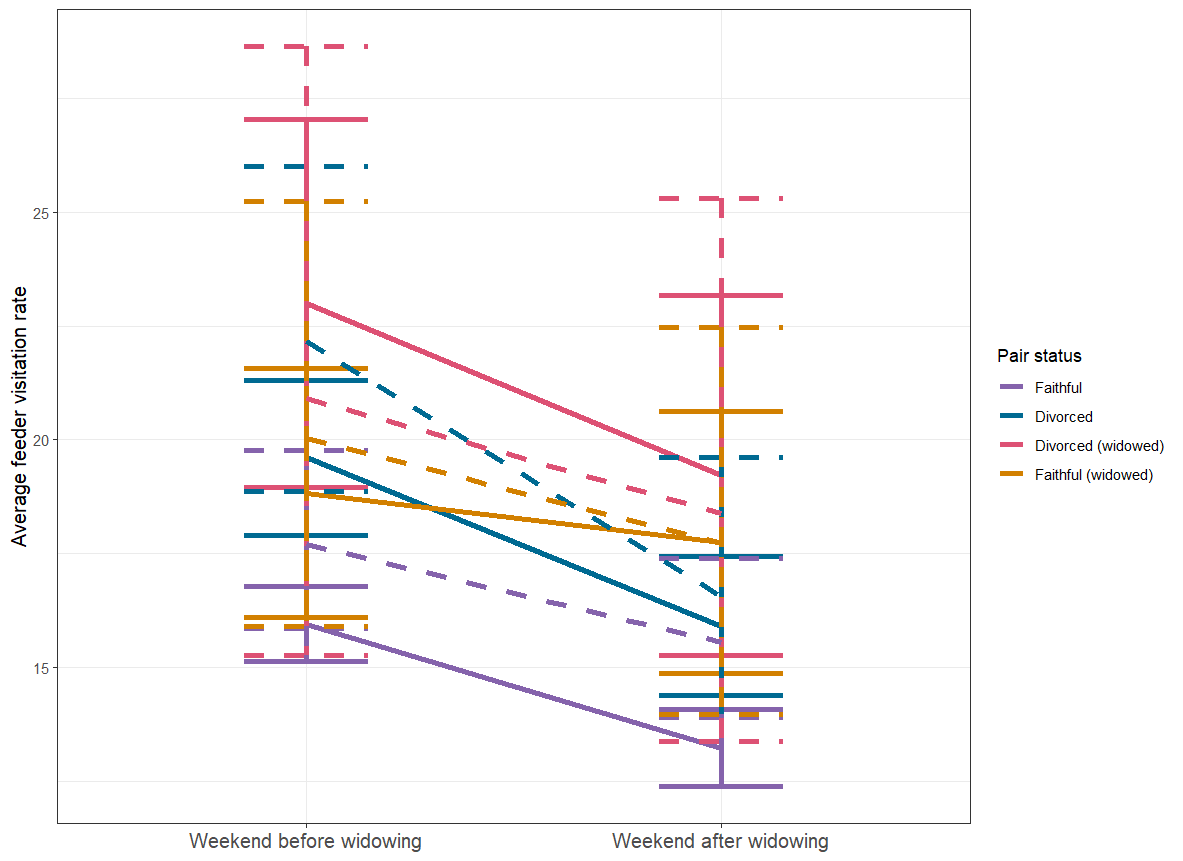

*Figure A38: Predicted and observed difference between average feeder visitation rate in the weekend before and after the loss of a bird’s mate (widowing). Solid points and lines are averages and standard error from the raw data, while dashed error bars and lines are predictions from a lognormal generalised linear mixed model, with upper and lower bars representing a 95% confidence interval. Predictions were made using ggeffects, accounting for random effects and a zero-inflation term (Lüdecke, 2018).*

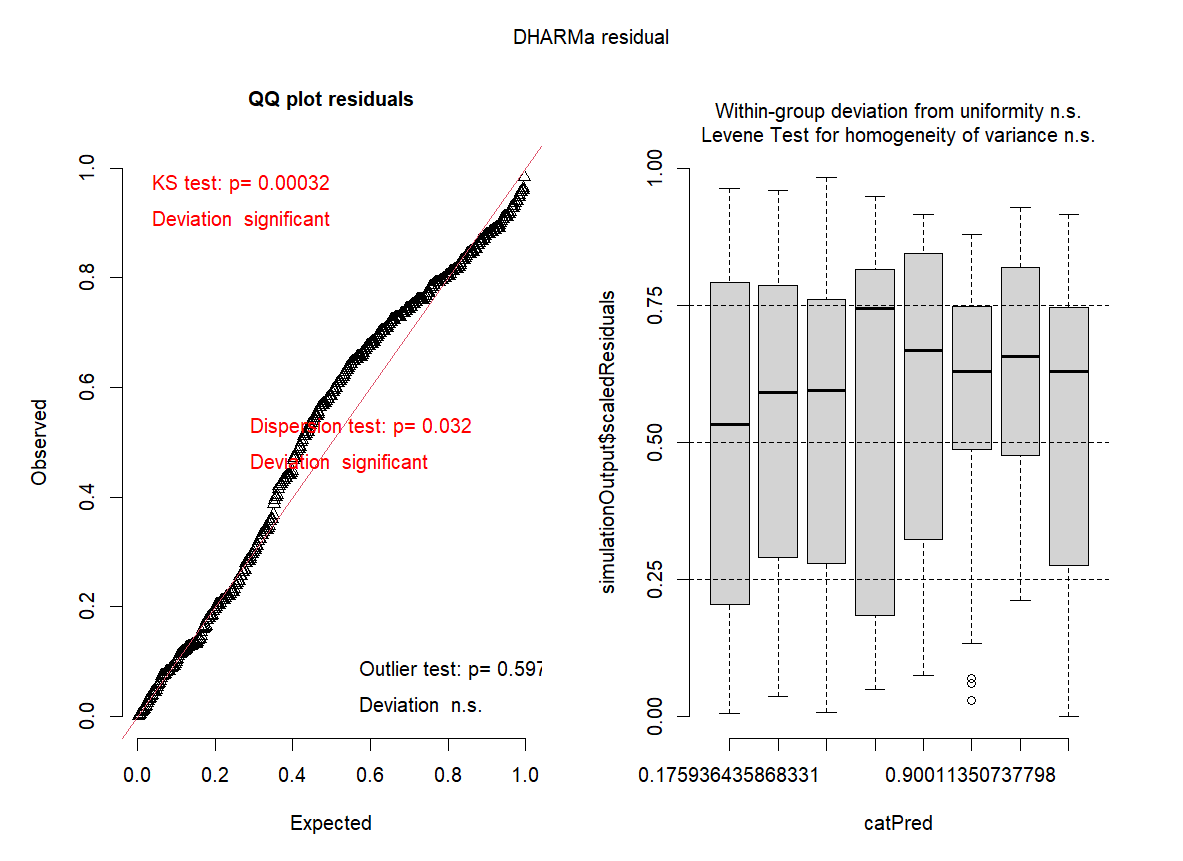

*Figure A39: Residuals from a lognormal GLMM of the effect of pair status and widowing on feeder visitation rate in the weekend before and after widowing, as calculated by DHARMa (Hartig, 2018).*

Preferred feeder location - period before/after

*Table A23: Fixed effect coefficient estimates from a binomial GLMM of the effect of pair status on the chance of moving preferred feeder after widowing. Significant coefficients are indicated in italics.*

| **Coefficient** | **Estimate** | **Standard error** | ***p*-value** |
| --- | --- | --- | --- |
| *Intercept* | *1.485* | *0.226* | *<0.001* |
| Pair status Divorced | -0.328 | 0.413 | 0.427 |
| Pair status Divorced (widowed) | -0.387 | 0.847 | 0.648 |
| Pair status Faithful (widowed) | 0.386 | 0.793 | 0.626 |

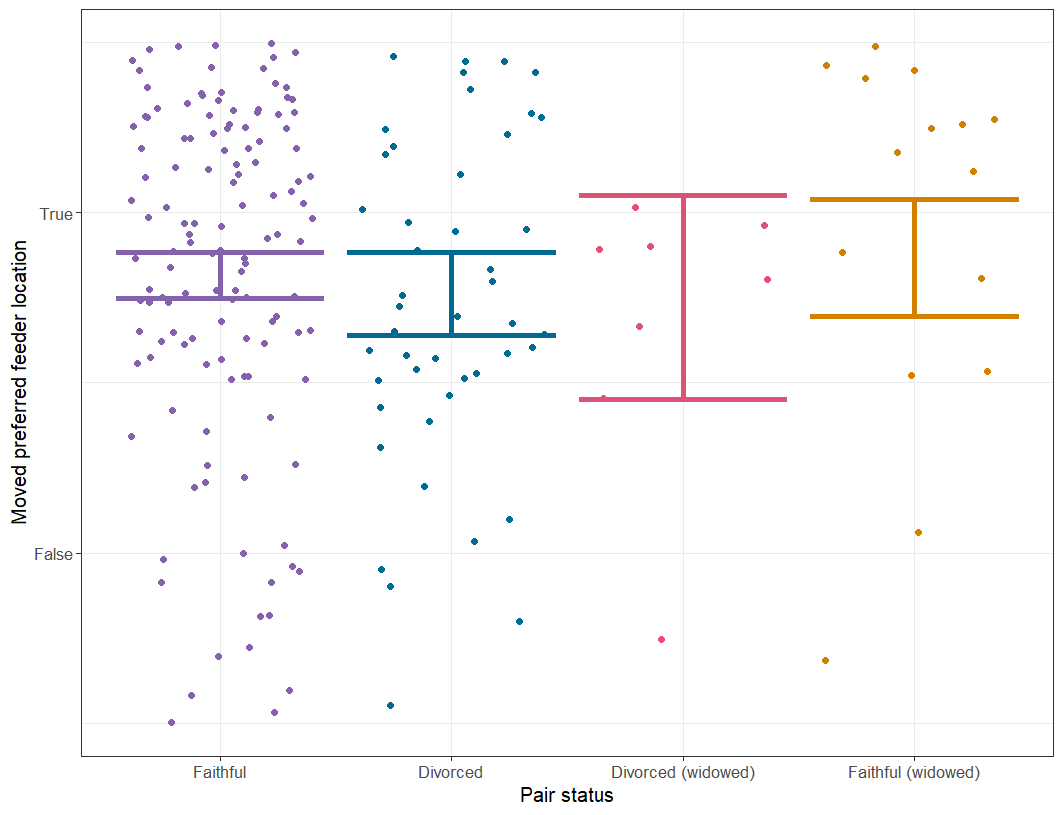

*Figure A40: Predicted and observed chance of moving preferred feeder location after the loss of a bird’s mate (widowing). Solid points and lines are averages and standard error from the raw data, while dashed error bars and lines are predictions from a lognormal generalised linear mixed model, with upper and lower bars representing a 95% confidence interval. Predictions were made using ggeffects, accounting for random effects (Lüdecke, 2018).*

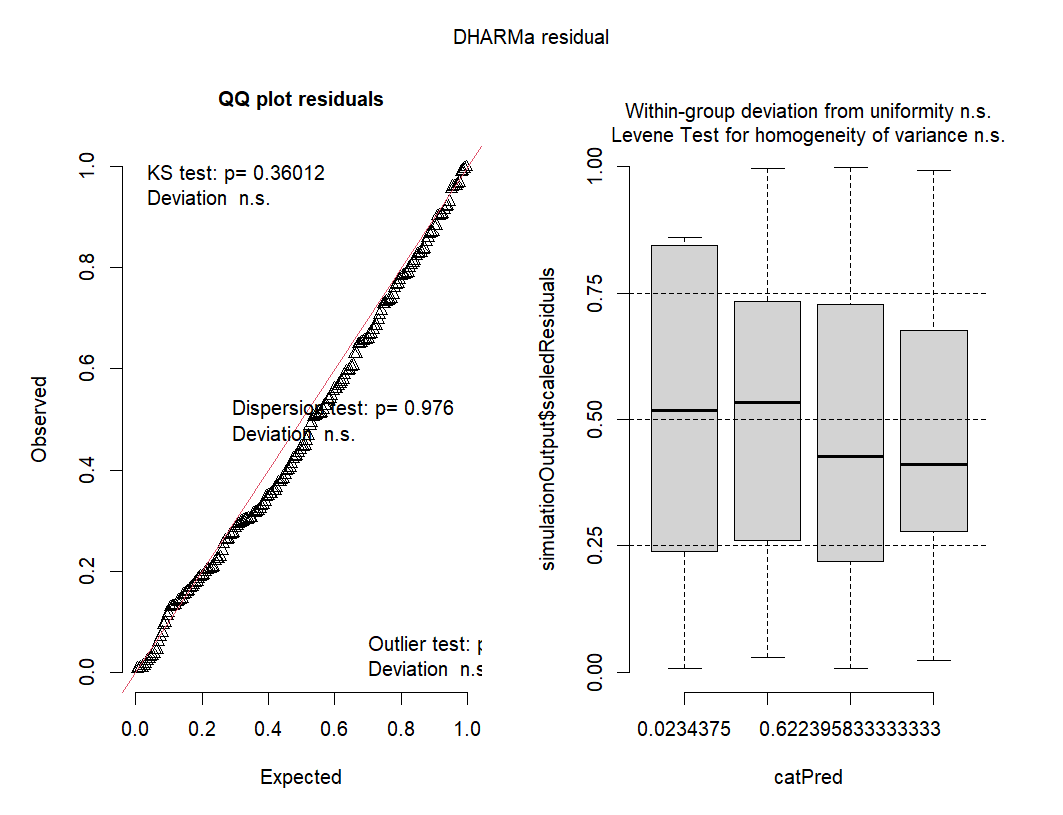

*Figure A41: Residuals from a binomial GLMM of the chance of moving preferred feeder location after widowing, as calculated by DHARMa (Hartig, 2018).*

Preferred feeder location - weekend before/after

*Table A24: Fixed effect coefficient estimates from a binomial GLMM of the effect of pair status on the chance of moving preferred feeder between the weekend before and after widowing. Significant coefficients are indicated in italics.*

| **Coefficient** | **Estimate** | **Standard error** | ***p*-value** |
| --- | --- | --- | --- |
| *Intercept* | *1.335* | *0.225* | *<0.001* |
| Pair status Divorced | -0.204 | 0.428 | 0.634 |
| Pair status Divorced (widowed) | -0.236 | 0.847 | 0.780 |
| Pair status Faithful (widowed) | 0.370 | 0.801 | 0.644 |

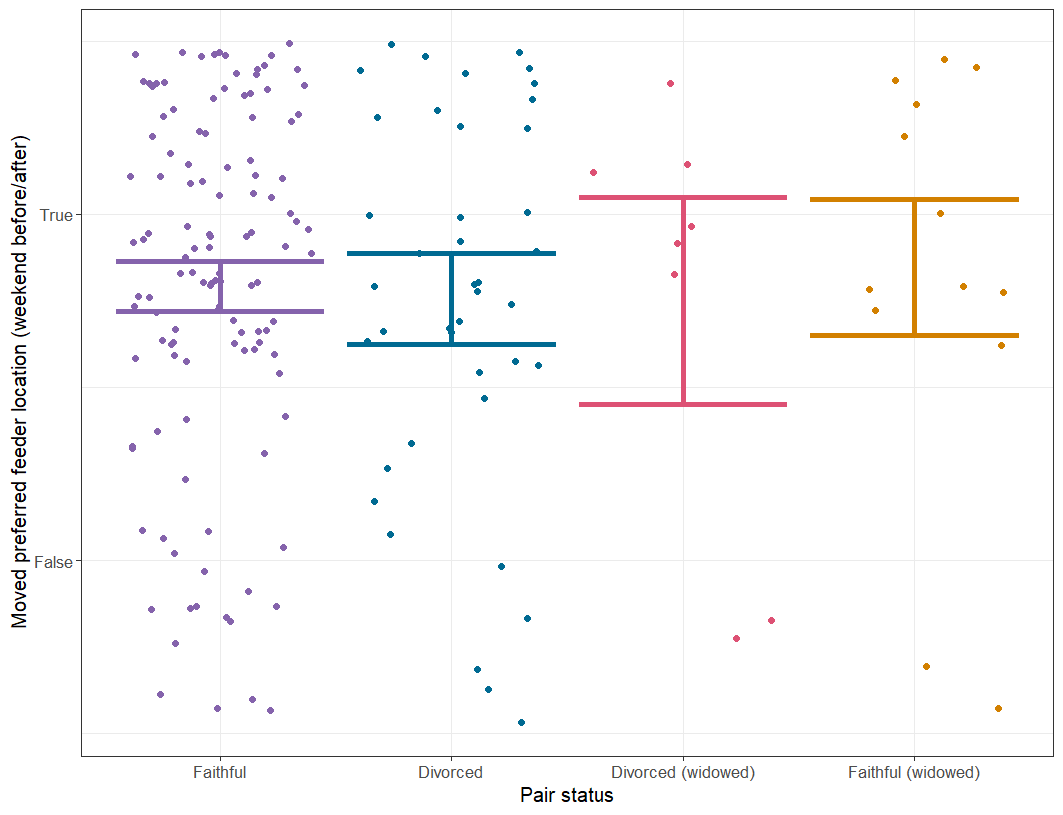

*Figure A42: Predicted and observed chance of moving preferred feeder location between the weekend before and after the loss of a bird’s mate (widowing). Solid points and lines are averages and standard error from the raw data, while dashed error bars and lines are predictions from a lognormal generalised linear mixed model, with upper and lower bars representing a 95% confidence interval. Predictions were made using ggeffects, accounting for random effects (Lüdecke, 2018).*

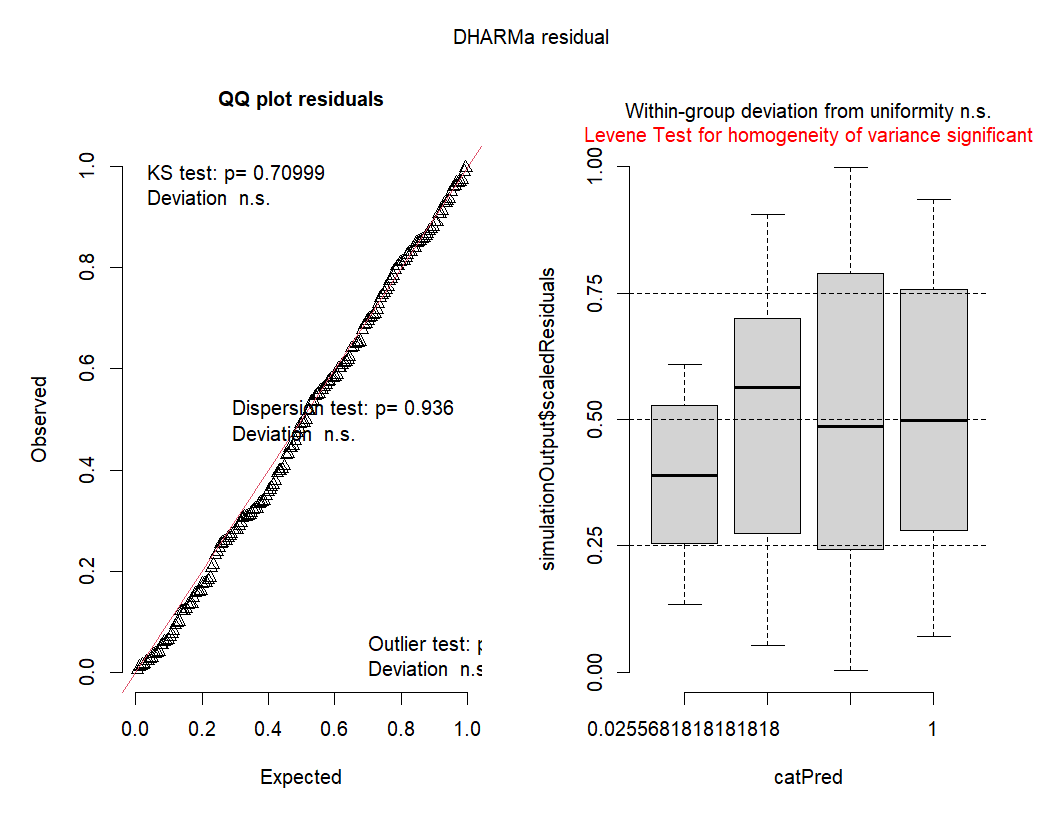

*Figure A43: Residuals from a binomial GLMM of the chance of moving preferred feeder location between the weekend before and after widowing, as calculated by DHARMa (Hartig, 2018).*

Top association score - period before/after

*Table A25: Fixed effect coefficient estimates from a lognormal GLMM of the effect of pair status and widowing on top winter association score. Significant coefficients are indicated in italics.*

| Coefficient | Estimate | Standard error | *p*-value |
| --- | --- | --- | --- |
| *Intercept* | *-1.222* | *0.044* | *<0.001* |
| *Pair status Divorced* | *-0.134* | *0.052* | *0.010* |
| Pair status Divorced (widowed) | -0.054 | 0.107 | 0.615 |
| Pair status Faithful (widowed) | 0.104 | 0.070 | 0.141 |
| *Post-widowing* | *0.156* | *0.027* | *<0.001* |
| Pair status Divorced:Post-widowing | 0.049 | 0.053 | 0.349 |
| Pair status Divorced (widowed):Post-widowing | 0.017 | 0.106 | 0.876 |
| *Pair status Faithful (widowed):Post-widowing* | *-0.191* | *0.084* | *0.023* |

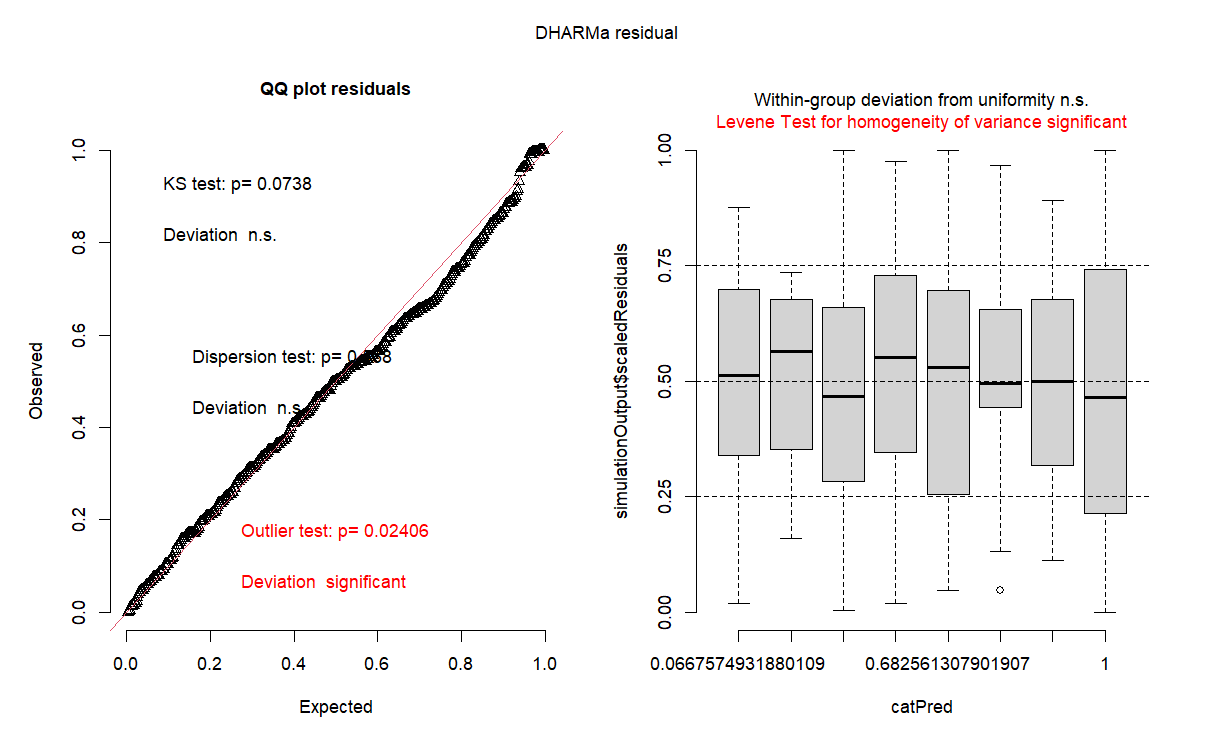

*Figure A44: Residuals from a lognormal GLMM of the effect of pair status and widowing on top winter association score, as calculated by DHARMa (Hartig, 2018).*

Top association score - weekend before/after

*Table A26: Fixed effect coefficient estimates from a lognormal GLMM of the effect of pair status and widowing on top association score in the weekend before and after widowing. Significant coefficients are indicated in italics.*

| Coefficient | Estimate | Standard error | *p*-value |
| --- | --- | --- | --- |
| *Intercept* | *-1.143* | *0.042* | *<0.001* |
| Pair status Divorced | -0.069 | 0.065 | 0.284 |
| Pair status Divorced (widowed) | -0.084 | 0.134 | 0.529 |
| Pair status Faithful (widowed) | 0.068 | 0.096 | 0.480 |
| Post-widowing | 0.049 | 0.040 | 0.223 |
| Pair status Divorced:Post-widowing | 0.020 | 0.076 | 0.791 |
| Pair status Divorced (widowed):Post-widowing | 0.061 | 0.161 | 0.702 |
| Pair status Faithful (widowed):Post-widowing | -0.137 | 0.122 | 0.259 |

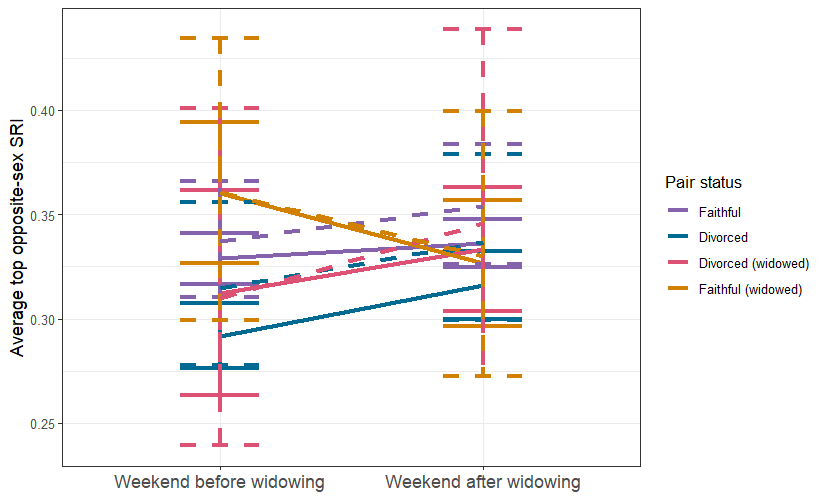

*Figure A45: Predicted and observed difference between average top association score in the weekend before and after the loss of a bird’s mate (widowing). Solid points and lines are averages and standard error from the raw data, while dashed error bars and lines are predictions from a lognormal generalised linear mixed model, with upper and lower bars representing a 95% confidence interval. Predictions were made using ggeffects, accounting for a zero-inflation term (Lüdecke, 2018).*

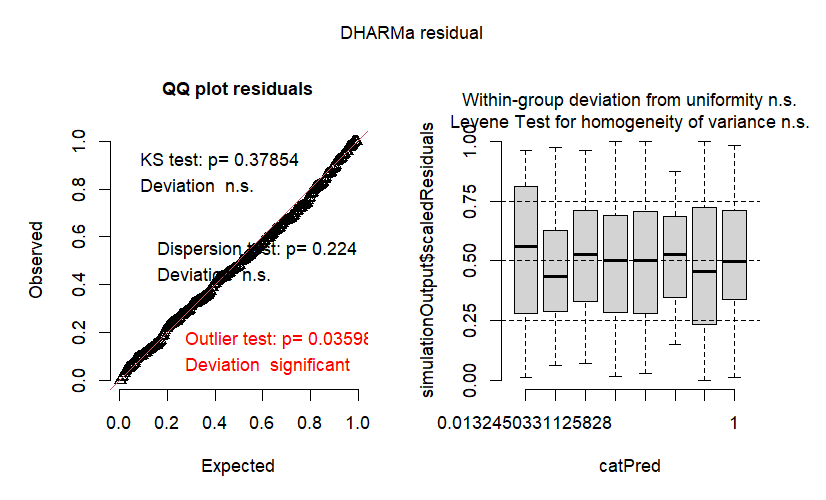

*Figure A46: Residuals from a lognormal GLMM of the effect of pair status and widowing on top association score in the weekend before and after widowing, as calculated by DHARMa (Hartig, 2018).*

Top visit adjacency index - period before/after

*Table A27: Fixed effect coefficient estimates from a lognormal GLMM of the effect of pair status and widowing on top visit adjacency index (with random effects included). Significant coefficients are indicated in italics.*

| Coefficient | Estimate | Standard error | *p*-value |
| --- | --- | --- | --- |
| *Intercept* | *-6.183* | *0.057* | *<0.001* |
| Pair status Divorced | 0.046 | 0.065 | 0.478 |
| Pair status Divorced (widowed) | -0.074 | 0.133 | 0.579 |
| Pair status Faithful (widowed) | -0.035 | 0.095 | 0.716 |
| Post-widowing | -0.070 | 0.046 | 0.129 |
| Pair status Divorced:Post-widowing | -0.124 | 0.090 | 0.169 |
| Pair status Divorced (widowed):Post-widowing | 0.023 | 0.194 | 0.904 |
| Pair status Faithful (widowed):Post-widowing | 0.000 | 0.142 | 1.000 |

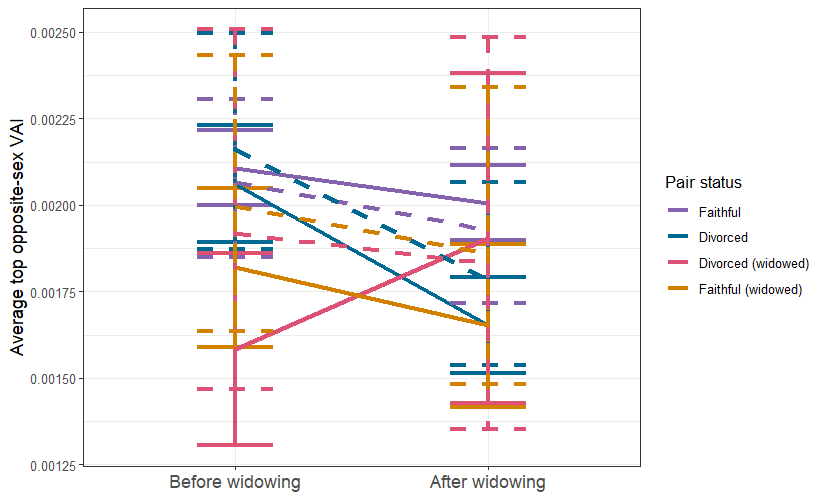

*Figure A47: Predicted and observed difference between average top visit adjacency index in before and after the loss of a bird’s mate (widowing). Solid points and lines are averages and standard error from the raw data, while dashed error bars and lines are predictions from a lognormal generalised linear mixed model, with upper and lower bars representing a 95% confidence interval. Predictions were made using ggeffects, accounting for a zero-inflation term (Lüdecke, 2018).*

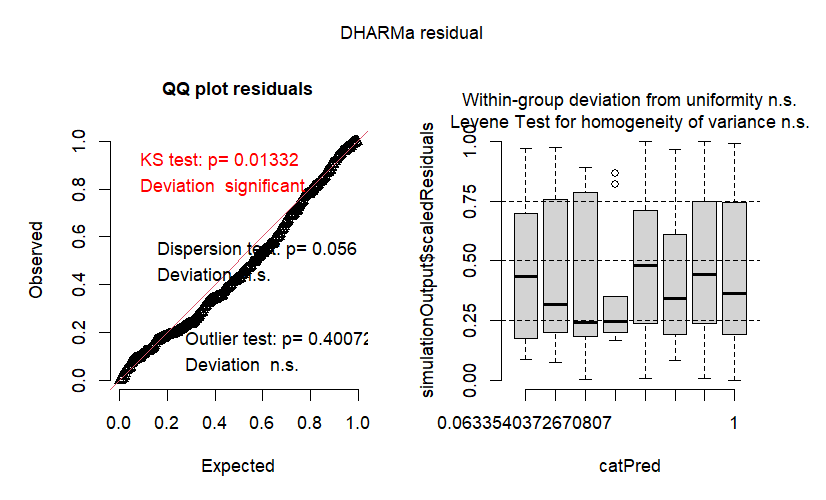
 *Figure A48: Residuals from a lognormal GLMM (with random effects included) of the effect of pair status and widowing on top visit adjacency index, as calculated by DHARMa (Hartig, 2018).*

Top visit adjacency index - weekend before/after

*Table A28: Fixed effect coefficient estimates from a lognormal GLMM of the effect of pair status and widowing on top visit adjacency index for the weekend before and after widowing (with random effects included). Significant coefficients are indicated in italics.*

| Coefficient | Estimate | Standard error | *p*-value |
| --- | --- | --- | --- |
| *Intercept* | *-6.314* | *0.075* | *<0.001* |
| Pair status Divorced | -0.030 | 0.083 | 0.713 |
| Pair status Divorced (widowed) | 0.152 | 0.147 | 0.301 |
| Pair status Faithful (widowed) | -0.055 | 0.131 | 0.676 |
| Post-widowing | 0.022 | 0.059 | 0.706 |
| Pair status Divorced:Post-widowing | 0.104 | 0.112 | 0.356 |
| Pair status Divorced (widowed):Post-widowing | -0.124 | 0.215 | 0.563 |
| Pair status Faithful (widowed):Post-widowing | 0.170 | 0.181 | 0.347 |

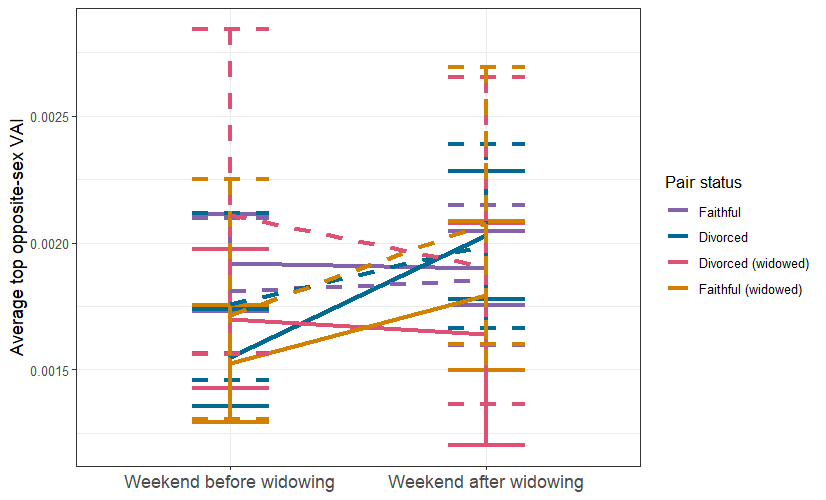

*Figure A49: Predicted and observed difference between average top visit adjacency index in the weekend before and after the loss of a bird’s mate (widowing). Solid points and lines are averages and standard error from the raw data, while dashed error bars and lines are predictions from a lognormal generalised linear mixed model, with upper and lower bars representing a 95% confidence interval. Predictions were made using ggeffects, accounting for a zero-inflation term (Lüdecke, 2018).*

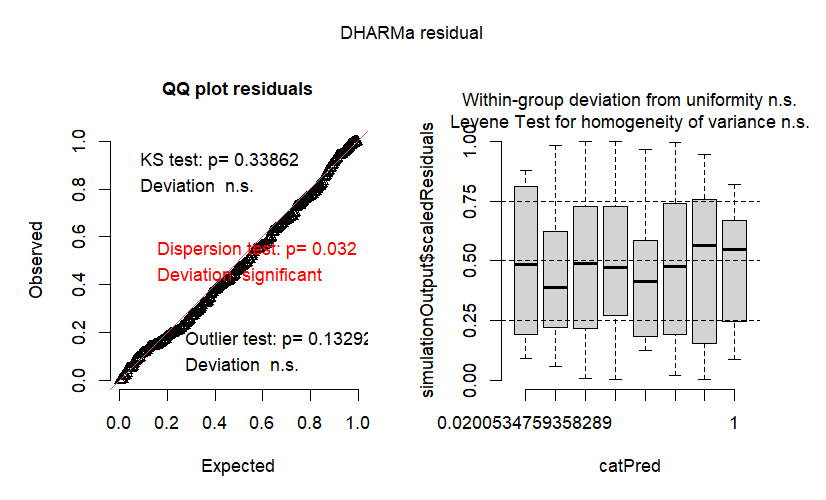
 *Figure A50: Residuals from a lognormal GLMM (with random effects included) of the effect of pair status and widowing on top visit adjacency index the weekend before and after widowing, as calculated by DHARMa (Hartig, 2018).*

Top association score - end of season

*Table A29: Fixed effect coefficient estimates from a lognormal GLMM of the effect of pair status on the top winter association score in the final weekend of data collection. Significant coefficients are indicated in italics.*

| **Coefficient** | **Estimate** | **Standard error** | ***p*-value** |
| --- | --- | --- | --- |
| *Intercept* | *-0.986* | *0.049* | *<0.001* |
| Pair status Divorced | -0.067 | 0.059 | 0.255 |
| Pair status Divorced (widowed) | 0.038 | 0.114 | 0.740 |
| Pair status Faithful (widowed) | -0.158 | 0.099 | 0.110 |

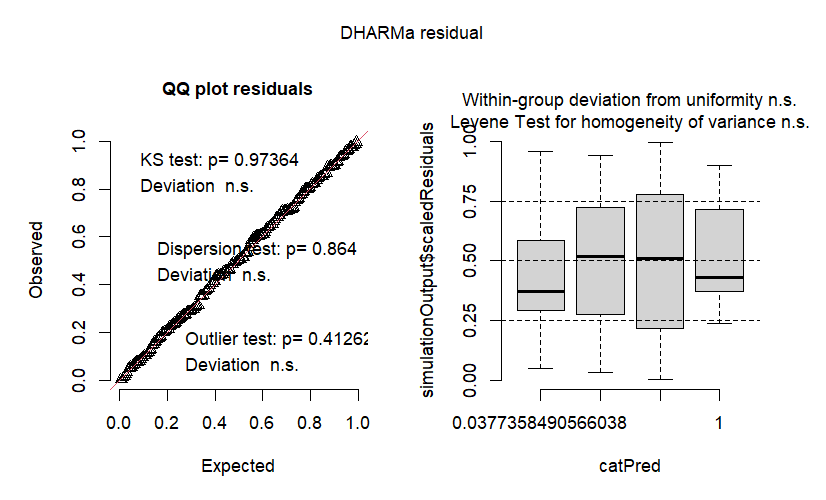

*Figure A51: Residuals from a lognormal GLMM of the effect of pair status on top winter association score in the final weekend of data collection, as calculated by DHARMa (Hartig, 2018).*

#### Sensitivity of non-widowed ‘widowing date’

In order to assess whether the particular distribution of randomly generated widowing dates used for non-widowed birds in this analysis may have led to the observed results, we conducted a sensitivity analysis. We regenerated widowing dates then reran the models 500 times. The z-scores of the faithful (widowed):predeath coefficient are presented below (Figure A52). As can be seen, most of the models showed that the model results are robust to differences in widowing dates used. The reported result for opposite sex degree (weekend after) is at the edge of the distribution of possible results. With a two-sided p-value, the result does not appear significant, but it is worth considering when interpreting results.

Above, in the permutation test of faithful/divorced statuses, a significant *p* value for permuted models of top association score reinforces our significant finding. However, here, a non-significant *p* does the same. This is due to the difference in what we are permuting. In the permutation test of faithful/divorced status, we are attempting to show that this particular distribution of pair statuses generates *unique* results. If true, it would show that the real distribution of statuses has qualities that would not be observed were the distribution different, suggesting a real effect. In the permutation of widowing date however, we are attempting to show that the randomly generated distribution of fake widowing dates does *not* generate unique results, as this would suggest that we have happened upon a particular random generation that changes the results we observe. This isn’t desirable, as the widowing dates were generated randomly simply to allow comparison between groups.

Overall, this sensitivity analysis has confirmed that the results of our analysis are not due only to methods used within the model building process.

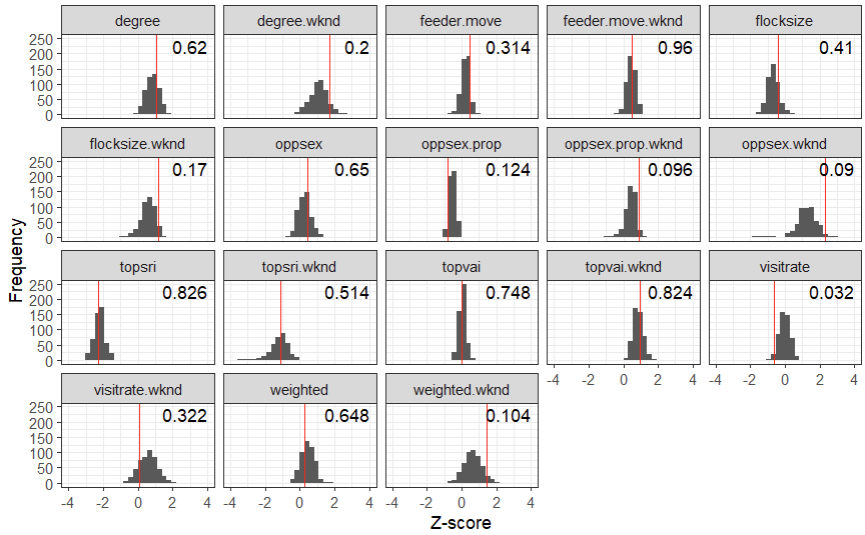
 *Figure A52: Permutation test of analysis models, generated by rerunning the models 500 times with widowing dates randomly assigned between pairs. Red lines represent the Z-score of the ‘Faithful (widowed):Pre-widowing’ coefficient in the original model, as presented in the main text. Text in the right hand side is the two-sided p-value of that original Z-score given the distribution of permuted results.*

#### Split sex models

We also produced each of the models reported in the main text but split by sex. The data was split in order to avoid the use of three way interaction terms, but presented only in the Supplementary Information as splitting the data by sex reduces the sample size significantly. The final sample sizes for each pair status split by sex are as follows:

*Table A30: Sample sizes of pair status and sex of individuals used in the analysis.*

|  | Females | Males |
| --- | --- | --- |
| Faithful | 77 | 77 |
| Divorced | 26 | 26 |
| Faithful (widowed) | 10 | 8 |
| Divorced (widowed) | 6 | 3 |

As these were preliminary supporting analyses with low sample sizes, only relevant coefficients are provided below, as most models were run with identical structure to the model that was not split by sex, only with separated data. Preferred feeder location was able to be modelled with an interaction term but had complete separation - in both the weekend and full season model, all male faithful (widowed) birds moved, preventing the calculation of accurate coefficients. Top association score at the end of the season was also modelled with an interaction term.

A table of faithful (widowed):postdeath coefficients for the models are presented below (Table A31). The only significant sex based effect was found in opposite sex degree in the weekend after widowing, where female widows showed a significant increase, but the same result was not observed for males.

*Table A31: Coefficient estimates, standard error, and p-values for models of various behavioural metrics, modelled separately for each sex. Reported coefficients are from the faithful (widowed):postdeath coefficient. Significant values are in bold and italics.*

|  | **Sex** | **Estimate** | **Std. error** | ***p*-value** |
| --- | --- | --- | --- | --- |
| **Daily degree** | Female | 0.177 | 0.124 | 0.154 |
|  | Male | -0.073 | 0.148 | 0.621 |
| **Daily degree (weekend)** | Female | 0.252 | 0.174 | 0.148 |
|  | Male | 0.149 | 0.195 | 0.443 |
| **Weighted degree** | Female | 0.116 | 0.114 | 0.312 |
|  | Male | -0.132 | 0.163 | 0.416 |
| **Weighted degree (weekend)** | Female | 0.192 | 0.171 | 0.261 |
|  | Male | 0.097 | 0.180 | 0.590 |
| **Opposite sex degree** | Female | 0.119 | 0.088 | 0.177 |
|  | Male | -0.094 | 0.152 | 0.536 |
| **Opposite sex degree (weekend)** | ***Female*** | ***0.309*** | ***0.154*** | ***0.045*** |
|  | Male | 0.220 | 0.210 | 0.295 |
| **Opposite sex proportion** | Female | -0.019 | 0.023 | 0.419 |
|  | Male | -0.000 | 0.018 | 0.979 |
| **Opposite sex proportion (weekend)** | Female | 0.020 | 0.031 | 0.534 |
|  | Male | 0.002 | 0.027 | 0.940 |
| **Flocksize** | Female | 0.107 | 1.174 | 0.928 |
|  | Male | -1.224 | 1.345 | 0.363 |
| **Flocksize (weekend)** | Female | 2.731 | 2.456 | 0.266 |
|  | Male | 0.658 | 2.228 | 0.768 |
| **Visitation rate** | Female | 0.006 | 0.140 | 0.966 |
|  | Male | -0.148 | 0.144 | 0.304 |
| **Visitation rate (weekend)** | Female | 0.264 | 0.162 | 0.103 |
|  | Male | -0.385 | 0.247 | 0.120 |
| **Top association score** | Female | -0.177 | 0.112 | 0.115 |
|  | Male | -0.235 | 0.130 | 0.071 |
| **Top association score (weekend)** | Female | -0.094 | 0.170 | 0.578 |
|  | Male | -0.197 | 0.180 | 0.274 |
| **Top visit adjacency** | Female | -0.039 | 0.201 | 0.846 |
|  | Male | 0.081 | 0.197 | 0.680 |
| **Top visit adjacency (weekend)** | Female | 0.128 | 0.267 | 0.630 |
|  | Male | 0.197 | 0.248 | 0.428 |
| **Top association score (end of season)** | Female | -0.041 | 0.132 | 0.759 |
|  | Male | -0.204 | 0.193 | 0.289 |

As results for opposite sex degree (weekend) was significant, full model residuals and coefficient tables are provided below. Additionally, we conducted further permutation tests of randomised pair statuses, to investigate whether the split sex results were due only to the smaller sample sizes for each category. Results for these are provided below.

Opposite sex degree (weekend before/after) - females

*Table A32: Fixed effect coefficient estimates from a lognormal GLMM of the effect of pair status and widowing on the opposite sex degree of females in the weekend before and after widowing. Significant coefficients are indicated in italics.*

| Coefficient | Estimate | Standard error | *p*-value |
| --- | --- | --- | --- |
| *Intercept* | *-3.450* | *0.069* | *<0.001* |
| Pair status Divorced | 0.151 | 0.133 | 0.257 |
| *Pair status Divorced (widowed)* | *0.539* | *0.178* | *0.003* |
| Pair status Faithful (widowed) | 0.082 | 0.174 | 0.636 |
| Post-widowing | -0.117 | 0.071 | 0.101 |
| Pair status Divorced:Post-widowing | 0.121 | 0.148 | 0.411 |
| Pair status Divorced (widowed):Post-widowing | -0.335 | 0.202 | 0.097 |
| *Pair status Faithful (widowed):Post-widowing* | *0.309* | *0.154* | *0.045* |

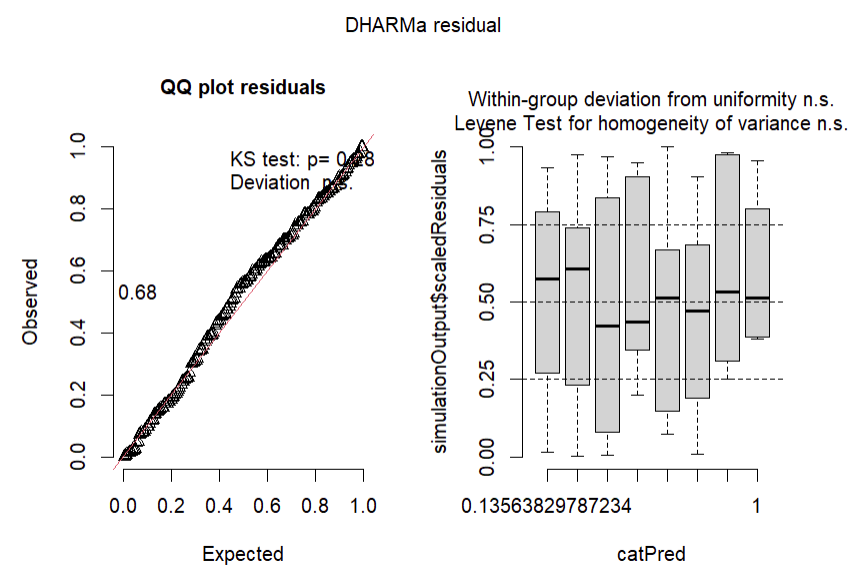

*Figure A53: Residuals from a lognormal GLMM of the effect of pair status and widowing on the opposite sex degree of females in the weekend before and after widowing, as calculated by DHARMa (Hartig, 2018).*

Opposite sex degree (weekend before/after) - males

*Table A33: Fixed effect coefficient estimates from a lognormal GLMM of the effect of pair status and widowing on the opposite sex degree of males in the weekend before and after widowing. Significant coefficients are indicated in italics.*

| Coefficient | Estimate | Standard error | *p*-value |
| --- | --- | --- | --- |
| *Intercept* | *-3.482* | *0.070* | *<0.001* |
| Pair status Divorced | 0.024 | 0.113 | 0.835 |
| Pair status Divorced (widowed) | -0.019 | 0.343 | 0.955 |
| Pair status Faithful (widowed) | 0.135 | 0.193 | 0.485 |
| Post-widowing | -0.079 | 0.066 | 0.233 |
| Pair status Divorced:Post-widowing | 0.072 | 0.133 | 0.588 |
| Pair status Divorced (widowed):Post-widowing | 0.372 | 0.343 | 0.279 |
| Pair status Faithful (widowed):Post-widowing | 0.220 | 0.211 | 0.295 |

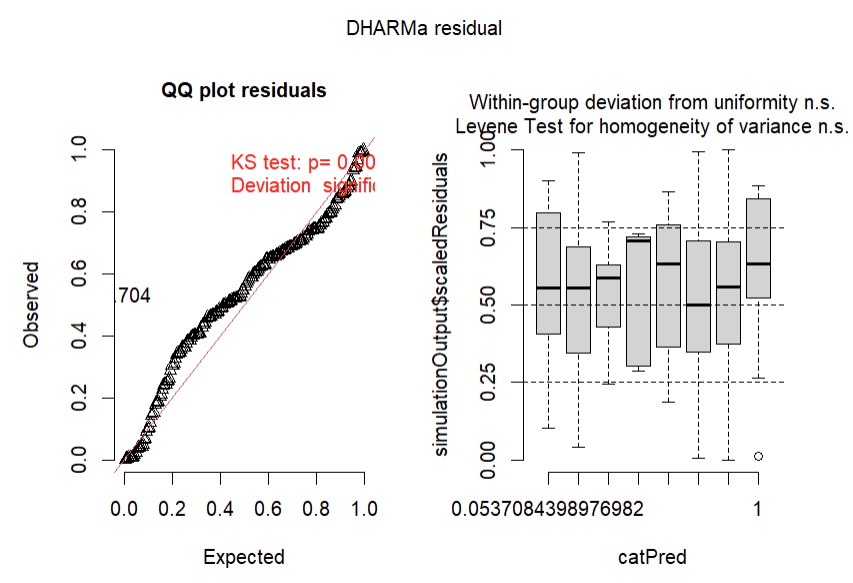

*Figure A54: Residuals from a lognormal GLMM of the effect of pair status and widowing on the opposite sex degree of males in the weekend before and after widowing, as calculated by DHARMa (Hartig, 2018).*

Permutation tests

For the model which found significant results when split by sex, we ran a permutation test, rerunning each model 1000 times, with pair status randomly assigned, while retaining the same numbers of each pair status (i.e 6 divorced widowed females, 3 divorced widowed males, 77 faithful females, etc.). This was in order to investigate whether the patterns observed in the models were significantly different from what we would expect to observe by chance, given the smaller sample sizes in the models. As seen in Figure A55, our observed results for the faithful (widowed):post-widowing coefficient Z-score were not significantly different from the results observed when the pair statuses were randomly assigned. Based on these permutation tests, we concluded that we do not have an appropriate amount of data to draw conclusions about sex effects on the results presented in the main text.

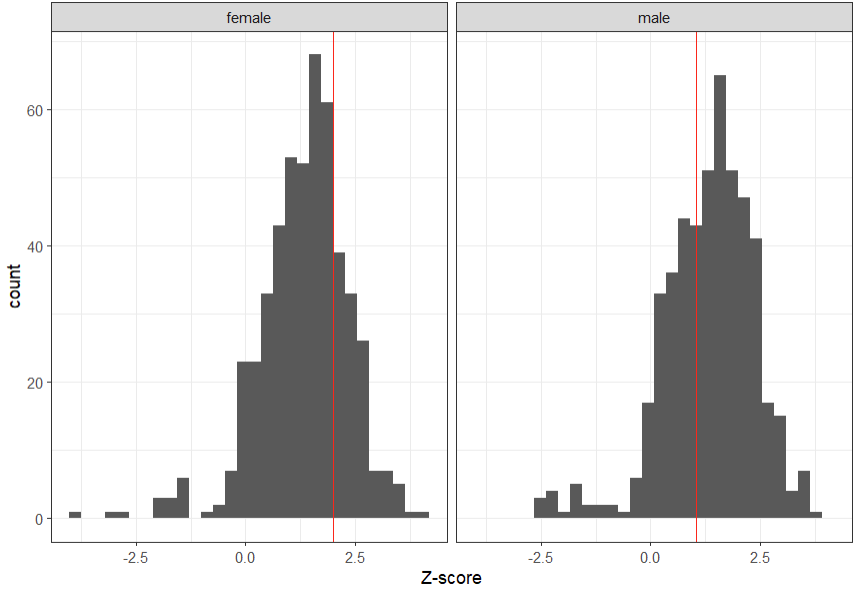

*Figure A55: Permutation test of sex split analysis models, generated by rerunning the models 500 times with sexes randomly assigned according to sample sizes from the original model. Red lines represent the Z-score of the ‘Faithful (widowed):Pre-widowing’ coefficient in the original model, as presented above.*
